# PreSMA population dynamics keep the beat atop the primate audiomotor hierarchy

**DOI:** 10.64898/2026.11.16.767170

**Authors:** Pablo Marquez, Yaneri Ayala, Luis Prado, Germán Mendoza, Hugo Merchant

## Abstract

Humans move in time with musical beats, yet how the brain builds the internal beat that guides movement is unknown. We recorded 4,719 neurons in six nodes of the audiomotor circuit while monkeys tapped in synchrony with auditory metronomes at different tempos. We found a clear functional dissociation in the temporal-frontal axis. Auditory cortex detected metronome clicks and contained local clocks that tiled each interstimulus interval, whereas parietal, striatal, and motor areas, most strongly the pre-supplementary motor area (preSMA), generated movement-aligned neural sequences that rescaled with tempo. Only in preSMA did population activity formed a cone-shaped manifold that jointly encoded tempo in its height, the produced interval in its width, and the timing of each tap relative to the beat in its position along the trajectory. During timing errors, preSMA activity kept conical shape but drifted toward shorter or longer tempos, revealing a hierarchical neural clock for beat synchronization.

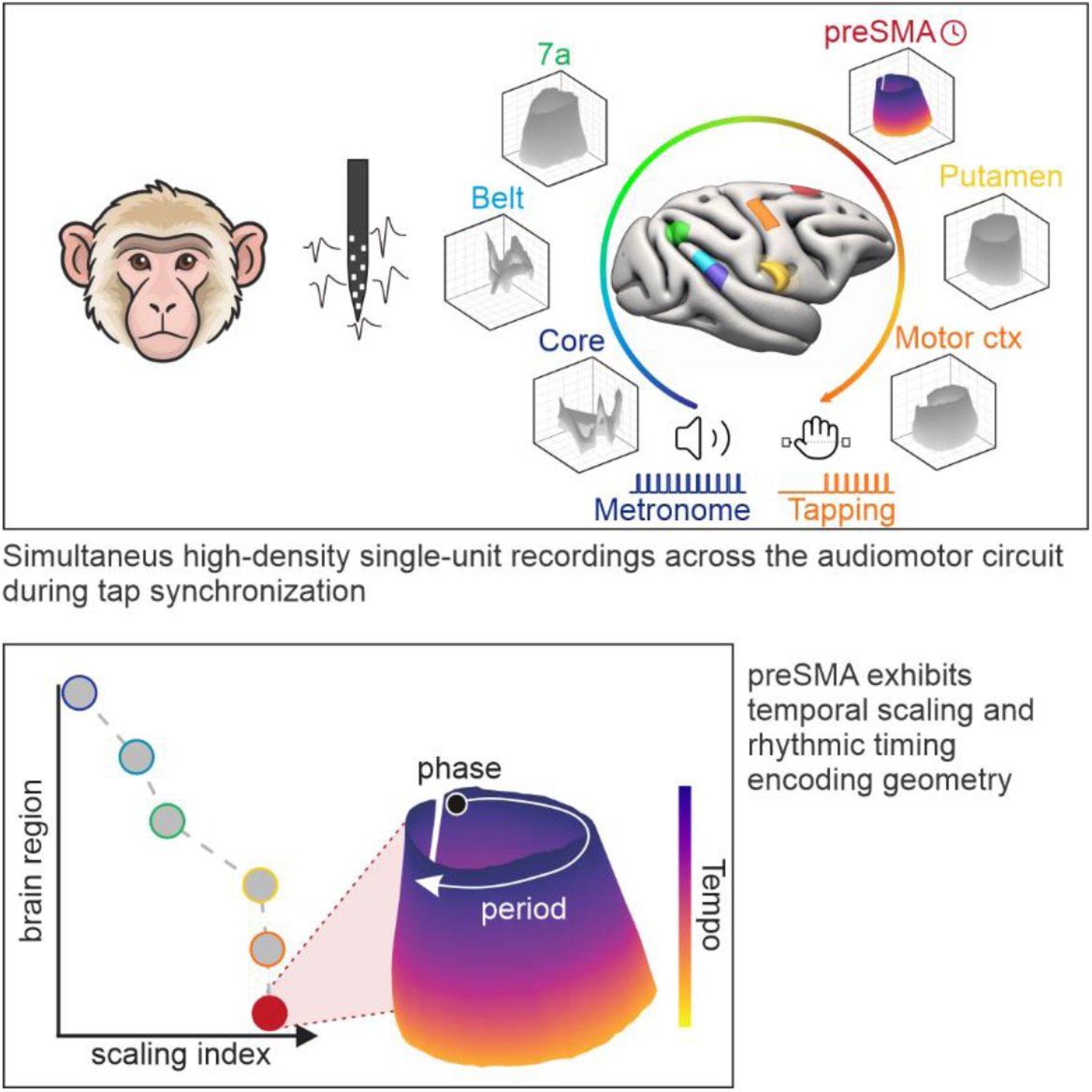
Graphical abstract.

**In brief:** A distinctive neural population geometry in macaque preSMA encodes metronome tempo and tap timing during rhythmic synchronization. This forms a behaviorally informative hierarchical manifold that is not found elsewhere in the audiomotor circuit.

**Highlights:**

- Timing signals are segregated and strengthen gradually from auditory to motor areas
- At the top of the hierarchy medial premotor activity forms a manifold that encodes the metronomés tempo and the period and phase of taps
- PreSMA manifold drift toward shorter or longer tempos before timing errors

## Introduction

Music cognition is an integral part of human life ^1^. Crucially, music develops in time, and humans spontaneously extract the beat of music and move in synchrony with it ^2,3^. Doing so requires extracting periodic events from a stream of sounds, building an internal representation of the beat, and using it both to predict the next beat and to launch an anticipatory movement that coincides with it ^4^. Functional imaging places these computations in an audiomotor circuit that includes the primary auditory cortex (AC) and the posterior auditory belt, the posterior parietal cortex, the medial premotor cortex (MPC: supplementary motor area [SMA] and presupplementary motor area [preSMA]), the basal ganglia, and the primary motor cortex ^5–8^. A leading hypothesis is that the internal beat is generated by the motor system^9–12^: beat perception engages the MPC and the putamen with or without movement ^13,14^, and motor areas could send a predictive signal back to the auditory system that modulates auditory gain and generates error signals to optimize synchronization ^15–18^. How this computation is divided among the nodes and whether there is a hierarchical structure within the circuit, however, remains unknown.

Rhesus monkeys are an excellent model to address this question. They synchronize predictively and flexibly to isochronous metronomes^19–22^ and tap to the subjective beat of music^4^. Their tapping combines a stereotyped movement, whose kinematics do not change across tempos or serial positions, with a dwell between movements whose duration increases with tempo ^22,23^. Monkeys, therefore, control the period of synchronization through the dwell and its phase through the timing of the stereotyped movement.

In the MPC, single neurons systematically increase or decrease their discharge rate as a function of time during rhythmic tapping^24–27^ but the beat clock is thought to depend on the interaction of neural populations ^28–30^. When MPC activity is projected into a low-dimensional state space, it follows circular dynamics that regenerate for every produced interval ^31–34^. Trajectories converge on a similar region of state space at tap times, which could define the phase of synchronization and provide a top-down predictive signal to sensory areas ^35^, and they increase in amplitude and decrease in speed as a function of tempo, encoding the dwell and the period of the metronome ^33,34^. This tempo code generalizes across auditory and visual metronomes ^33^. Together, these properties define a concise neural population chronometer ^36^.

Population activity often evolves within low-dimensional neural manifolds ^37–39^, whose geometry can carry behavioral variables such as spatial position or accumulated evidence ^40,41^. In single-interval timing, tonic inputs shift trajectories to different regions of state space and thereby set the speed at which they unfold and the interval they encode ^42^, which could give the MPC flexibility to time different intervals ^28^. Whether a comparable manifold underlies beat synchronization, and where in the audiomotor circuit it arises, is unknown.

Here, we recorded single neurons in six nodes of the macaque audiomotor circuit while monkeys synchronized their taps to auditory metronomes of four tempos or listened passively to the same metronomes. We asked how timing properties of cells change along the circuit and whether the geometry of population activity in any node captures the period and phase of synchronization, including when synchronization fails, to distinguish between a hierarchical or a distributed beat clock ^43,44^. We found a gradient of timing properties from auditory to motor areas that culminates in preSMA, where population activity forms a cone-shaped manifold that encodes tempo, the period of the produced interval, the phase of each tap, and the drift that precedes timing errors. preSMA is the only area within the recorded nodes of the audiomotor system where the population dynamics explains the tempo of the auditory metronome, the period and phase of the produced intervals, and the early and late errors, supporting the notion of a hierarchical organization in the neural mechanism that governs rhythmic synchronization.

## Results

### Monkeys set the period and adjust the phase of synchronization tap by tap

Two monkeys were trained on a perception-synchronization task (PST; Figure 1). Each animal held a lever during four metronome stimuli (ts_p1-4_ in Figure 1C; perception epoch, three intervals) and then produced seven fast forward-backward movements over a plate with infrared sensors in synchrony with the metronome (ts_s1-7_; synchronization epoch), yielding six produced intervals (PI_1-6_; Figure 1C; see Methods). The movements were spontaneously adopted by the monkeys, produced no sound, and had kinematics similar to those of push-button tapping ^23,33^; we therefore refer to them as taps. The interonset or target interval (TI) of the metronome was 550, 650, 750, or 850 ms, presented pseudorandomly. In a passive condition, the monkeys listened to the same metronomes without access to the lever, the sensor plate, or the reward tube, and did not move (Figure 1D). Thus, the PST is a complex and demanding task with an initial epoch where monkeys needed to pay attention to the tempo of the metronome without moving and then synchronize their movements to the periodic stimuli. In contrast in the passive condition the animals did not need to pay attention or to respond to the same isochronous auditory sequence, allowing to distinguish between the auditory input neural responses from the encoding of an internal predictive clock used to synchronize the movement.

**Figure 1.**
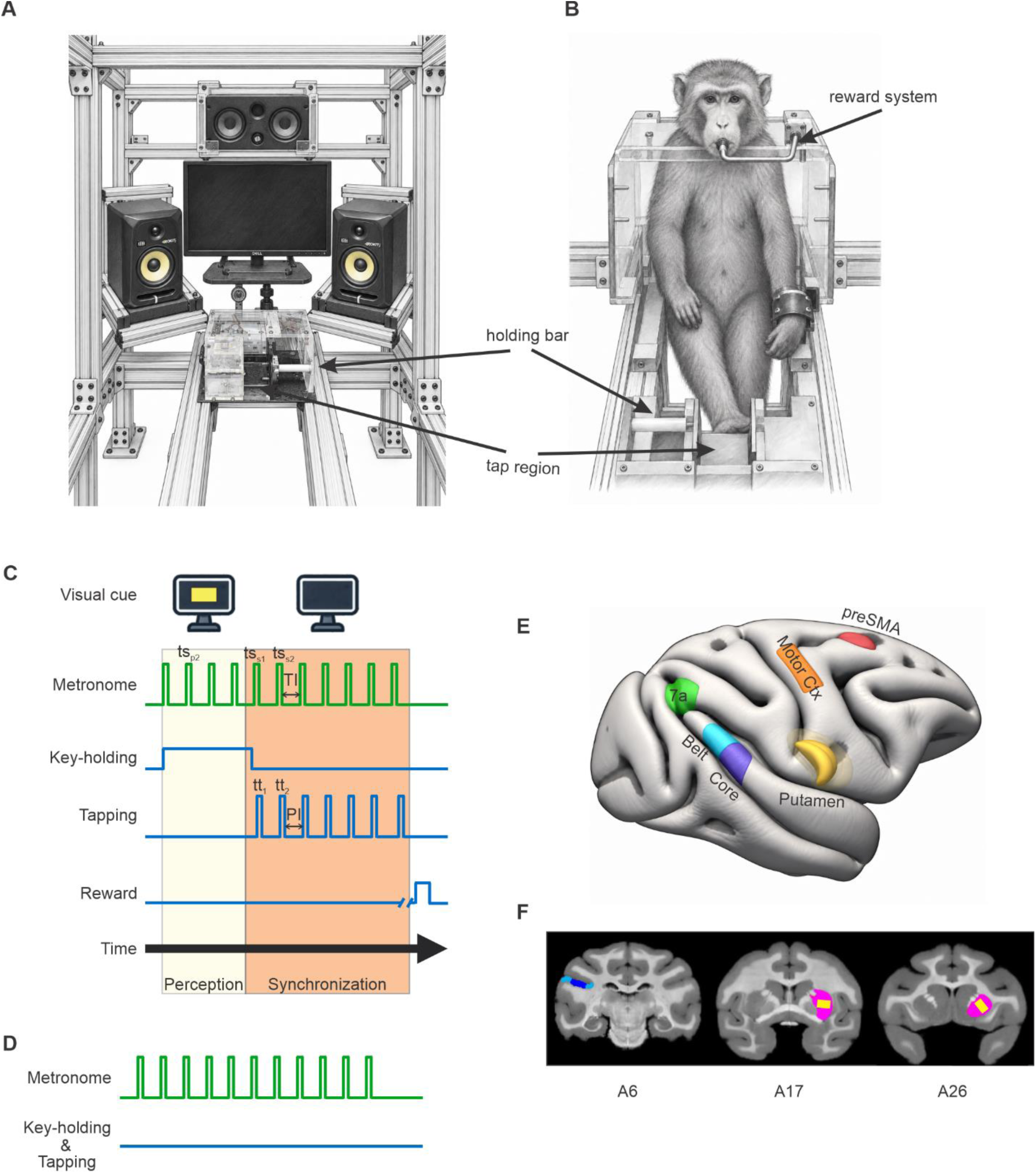
Experimental setup. (A) System for auditory stimulus presentation, including a screen for the visual *go* cue. (B) Monkey seated in a primate chair compatible with the system, with a tube for reward delivery. (C) Perception-synchronization task (PST). A trial started when the monkey held a lever during the presentation of a yellow background and of short clicks that acted as a metronome. The disappearance of the yellow background was the go signal to move in synchrony with the metronome by crossing an optical sensor with a forward-backward movement, producing six intervals (PI1 to PI6). The target interval (TI) of the metronome was pseudorandomly selected from four intervals (550–850 ms). (D) Passive condition. The monkeys listened to the same metronomes without access to the lever, the sensor plate, or the reward tube, and did not move. (E) Cortical recording areas, recorded with high-density silicon probes: posterior auditory core (marine blue), posterior belt (cyan), area 7a of the posterior parietal cortex (green), preSMA (red), and primary motor cortex (orange). (F) Coronal MRI sections of monkey 1 showing the recording sites in the medial and lateral posterior belt (cyan) and the posterior core (marine blue) (left), and in the putamen (yellow corresponds to the recorded area and pink depicts the contour of all the putamen; middle and right). The anterior coordinate of each section relative to the interaural zero is shown at the bottom (Saleem and Logothetis, 2012).

In the PST we quantified the phase of synchronization with the asynchrony (tap minus stimulus time), its accuracy with the constant error (PI minus TI), and its precision with the temporal variability (SD of the PI). Asynchronies were close to zero, slightly positive for short TIs and negative for the longest TI (Figure 2A; one-way ANOVA, effect of TI: F(3, 8606) = 1,198.1 for monkey 1 and F(3, 9005) = 2,323.5 for monkey 2, both p < 0.001). Produced intervals formed largely non-overlapping distributions that followed the TI (Figure 2B; F(3, 8606) = 49,686 and F(3, 9005) = 43,729, both p < 0.001), and constant errors were small but varied with TI (Figure 2C; F(3, 8606) = 11.22 and F(3, 9005) = 71.06, both p < 0.001). Bonferroni-corrected post hoc comparisons were significant for all pairs of TIs, except for the constant error of monkey 1 at 550 vs. 650 ms, 550 vs 750 ms and 750 vs. 850 ms; monkey 2 550 vs 650 ms and 750 vs 850 ms. Thus, both monkeys synchronized predictively and accurately across tempos.

**Figure 2.**
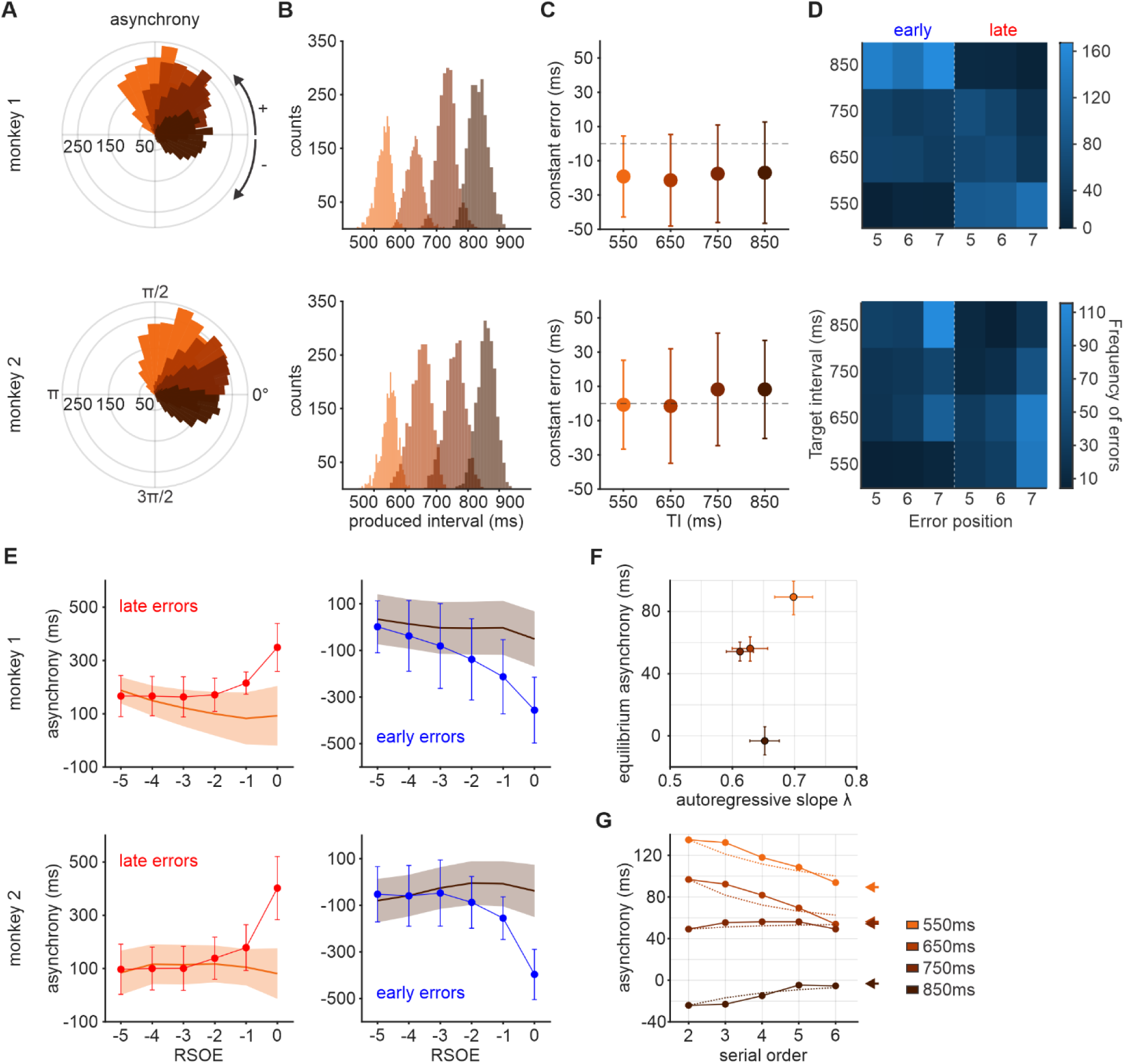
Behavior during the PST. (A–E) Upper and lower rows correspond to monkey 1 and monkey 2, respectively. The number of correct trials for each TI (550, 650, 750, and 850 ms) was 2148, 2130, 2148, and 2184 for monkey 1, and 2250, 2255, 2249, and 2255 for monkey 2. (A) Polar histograms of the mean asynchrony per trial for each TI, expressed as phase relative to the TI (positive, counterclockwise; negative, clockwise). The radial axis represents trial counts. (B) Distributions of produced intervals for each TI. (C) Constant error for each TI (mean ± SD across trials). (D) Frequency of early and late errors for each TI and serial order of occurrence (5–7). (E) Mean asynchrony as a function of the relative serial order to error (RSOE; 0, error tap; negative values, preceding taps) for late errors at 550 ms (red) and early errors at 850 ms (blue), the TIs at which each error type was most frequent. Solid lines and shaded bands show the mean ± SD of correct trials; circles and error bars show the mean ± SD of error trials. (F) Autoregressive slope λ versus equilibrium point A*, from the fitted relation between successive asynchronies for both monkeys and each TI (see Supplementary Figure 2); error bars, 95% session-bootstrap confidence intervals. Correction strength was similar across TIs, whereas A* was strongly displaced across TIs. (G) Mean asynchrony as a function of serial order (solid lines) and profile predicted from the tap-to-tap asynchrony dynamics (dotted lines), using λ and A* estimated independently of serial order and anchored to the first observed tap; arrowheads indicate A* (linear fits, p < 0.0001). TI color code as in (C).

Monkeys 1 and 2 nevertheless made errors on 17% and 9.5% of trials, respectively; a trial was aborted when the asynchrony of any tap exceeded ±280 ms. Early errors were most frequent at the longest TI (850 ms) and late errors at the shortest TI (550 ms) (Figure 2D), so each error type was analyzed at that TI. Aligning errors occurring at serial orders 5–7 to the error tap (relative serial order to error, RSOE = 0) showed that asynchronies deviated progressively from those of correct trials over the three preceding taps, culminating in an abrupt loss of synchrony (Figure 2E; two-way mixed ANOVAs, trial type × RSOE interaction for early and late errors in both monkeys, all p < 0.001).

To determine how the monkeys controlled period and phase, we decomposed the variance of produced intervals and asynchronies into contributions of TI, serial order, and trial-wise offset fluctuations. The TI explained almost all the variance of the produced intervals, whereas TI, serial order, trial offset, and a large residual all contributed to the variance of the asynchronies (Supplementary Figure 1A). Both parameters showed a strong lag-1 autocorrelation (≈0.7), but only the asynchrony retained it after the deterministic task structure was removed (Supplementary Figure 1B). Thus, the period was predefined, presumably during the perception epoch, whereas the phase was controlled dynamically during each trial.

Consecutive asynchronies followed a linear relation, async_k+1_ = c + λ·async_k_, with λ below one for all TIs (Figure 2F; Supplementary Figure 2): a positive asynchrony tended to be followed by a less positive one and a negative asynchrony by a less negative one. The correction strength λ was similar across TIs, whereas the equilibrium point A* = c/(1 − λ) was positive for short TIs and negative for the 850-ms TI (Figure 2F). A recursive model parametrized only by λ and A* reproduced the systematic drift of the asynchrony toward A* along the sequence (Figure 2G). Hence, the monkeys set the period of synchronization in advance and controlled its phase tap by tap, consistent with our asynchrony-based reward contingencies.

## Synchronization engages the audiomotor circuit along a functional gradient

We recorded 4,719 neurons with high-density silicon probes in six areas of the audiomotor circuit (Figures 1E and 1F; Supplementary Figure 3; Supplementary Table 1): the posterior auditory core, the posterolateral belt, area 7a of the posterior parietal cortex, the putamen, the primary motor cortex (M1), and preSMA. Because response properties were very similar in both monkeys, neurons were pooled across animals.

Core and belt neurons were mainly aligned to the stimuli, whereas parietal, putamen, M1, and preSMA neurons were aligned to the taps (Supplementary Figure 4), as determined with a warping analysis ^45^. We therefore analyzed auditory cortex in absolute time and the other areas in time warped to the taps and grouped the neurons of each area into response clusters using UMAP followed by k-means clustering (see Methods; Supplementary Table 2).

Auditory cortex showed a rich repertoire of responses (Supplementary Figures 5 and 7; Supplementary Table 3). Besides short-latency responses phase-locked to the stimuli, the core (six clusters) and belt (seven clusters) contained clusters whose activity covered the interval between stimuli, most clearly core cluster 5 (an inhibition followed by a rebound) and belt cluster 6 (a neural sequence), as well as tap-aligned clusters that could carry proprioceptive or top-down beat signals. In contrast, parietal cortex, putamen, M1, and preSMA each showed three tap-aligned clusters that divided the interval between taps into roughly three equal parts, forming neural sequences that tiled the produced interval at all tempos (Supplementary Figures 5 and 6); in area 7a, the third cluster was phase-shifted, with peaks before and after the tap.

Importantly, a permutation test confirmed that both phase-locked responses and neural sequences were consistent across trial repetitions and did not result from sorting the activity by response latency (Supplementary Figure 8). Notably, these finding supports the notion of a functional segregation, with most of the neurons in the core and belt encoding the presence of each auditory stimuli in absolute time, while the frontal areas focus on the relative timing between the rhythmic taps.

We next compared the activity of each cluster between the perception and synchronization epochs (Figures 3A–3C; temporal cluster-based sign-flip permutation test). Figure 3A depicts the difference in the integrated firing rate between the two epochs, where the green bars denote the task epoch with the largest difference in discharge rate (bars to the left indicate larger activity during perception; bars to the right, during synchronization) and the size of the bar corresponds to the magnitude of the difference between epochs. In addition, we included a gray bar when there was a structured activation in the other epoch, but with a magnitude smaller than that of the green bar epoch. For example, at the top of Figure 3B with the arrow 1, we depict the difference in the profile of activation between the synchronization and perception epochs for the stimulus phase-locked responses of cluster 1 of the core. The overall discharge rate is larger in the perception epoch, as indicated by the large negative green density in the synchronization-minus-perception difference across the 850-ms interval (Figure 3B top). In addition, this cluster has a large, short-duration activation that is phase-locked to the stimuli, depicted in gray, which has a slightly larger magnitude for the synchronization epoch. Figure 3C (number 1 on the top) shows the average discharge rate for one neuron of this cluster aligned to the stimuli for the three intervals of the perception and the six intervals of the synchronization, recapitulating what we have just described. All core and belt clusters except core cluster 5 were more active during perception, although the phase-locked response of core cluster 1 was slightly larger during synchronization (arrow 1). This suggests that a tonic inhibitory input during tapping suppresses ongoing auditory activity and sharpens stimulus-evoked responses. In contrast, the putamen, M1, and especially preSMA were strongly biased toward synchronization, with cyclic activations locked to the taps that were absent, or even slightly inhibited, during perception (arrows 4 and 5). Parietal clusters were balanced between epochs (arrow 3), consistent with a role in sensorimotor integration.

**Figure 3.**
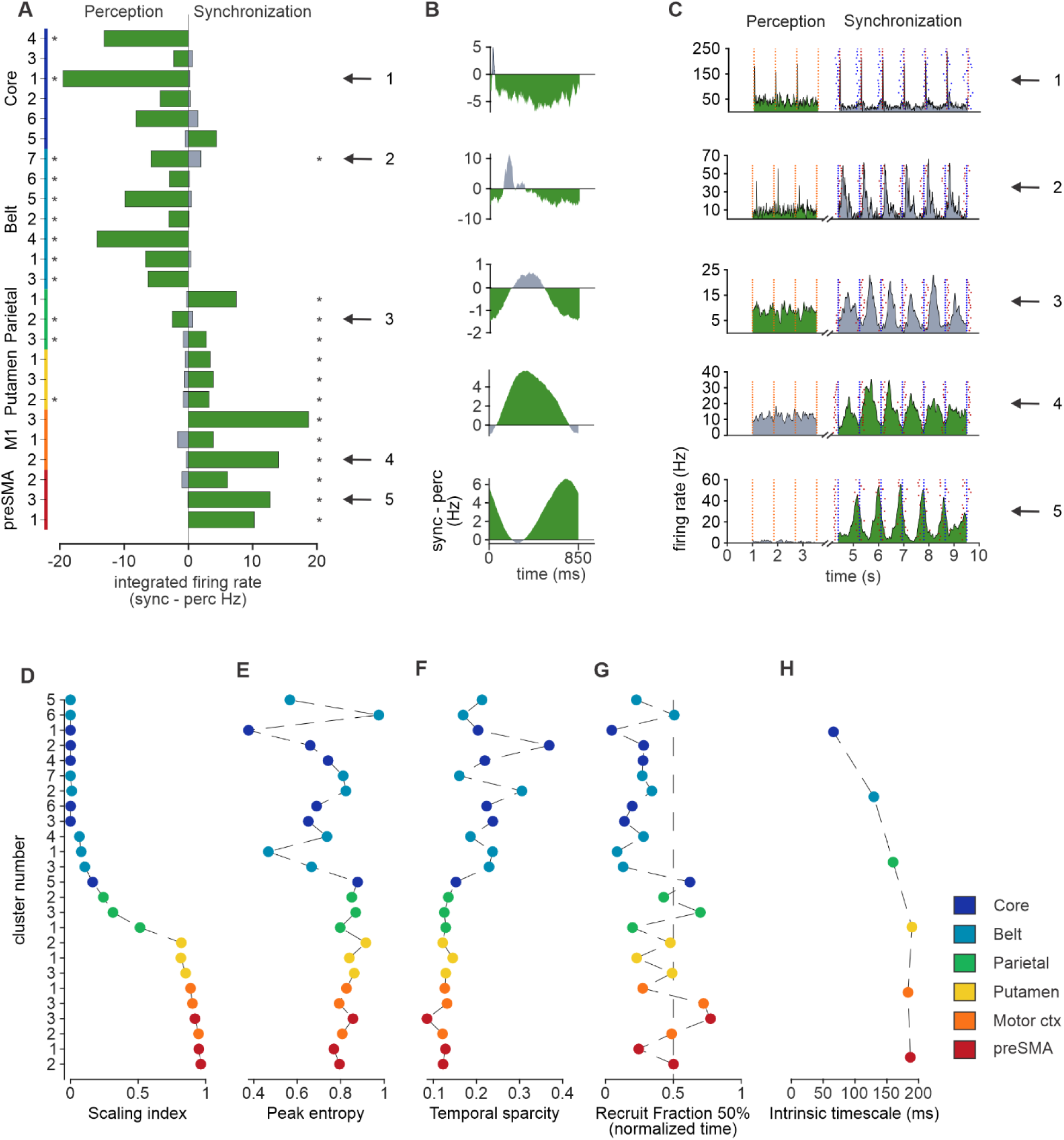
Task-epoch dependence and intrinsic timing properties across areas. (A) Difference in integrated firing rate between the perception and synchronization epochs for each area and cluster. Green bars indicate the epoch with the larger activity (perception, left; synchronization, right), and their length the magnitude of the difference; gray bars indicate structured activity of smaller magnitude in the other epoch. Asterisks, significant difference from zero (temporal cluster-based sign-flip permutation test). (B) Synchronization-minus-perception difference in mean firing rate as a function of time for the 850-ms TI, for the clusters marked by arrows in (A). Negative densities (arrows 1 and 2) indicate larger activity during perception; positive densities (arrows 4 and 5), larger activity during synchronization. (C) Firing rate of prototypical neurons from the clusters marked in (A) during perception (left) and synchronization (right) for the 850-ms TI. Traces are green for the epoch with the larger integrated firing rate. Orange dots, stimuli during perception; red dots, stimuli during synchronization; blue dots, taps. For arrows 3–5, activity during synchronization was aligned to the taps using dynamic time warping. (D) Temporal scaling index for the clusters of the six areas (color coded). (E) Peak entropy. (F) Temporal sparsity. (G) Recruitment fraction at 50% of the active population; the dashed line marks half of the population recruited at half of the normalized produced interval (425 ms). (E–G) Same format as (D). (H) Intrinsic timescale for each of the six areas.

During passive listening, phase-locked auditory clusters (core clusters 1, 3, 4, and 5; belt clusters 1–6) still responded to the stimuli, but their responses were smaller and had longer latencies than during synchronization (Supplementary Figures 7, 9, and 10). Clusters in all other areas were active almost exclusively during synchronization, and the neural sequences of parietal cortex, putamen, M1, and preSMA lost their consistency across trials during passive listening (Supplementary Figure 8). Thus, auditory areas access metronome information regardless of behavior, whereas the entire audiomotor circuit is engaged when rhythmic input drives movement.

## Timing properties increase gradually along the circuit

Temporal scaling, the stretching or compression of responses with the timed interval, is a hallmark of timing circuits ^22,27,46,47^. Scaling indices were low in phase-locked auditory clusters, higher in the remaining core and belt clusters, increased in parietal cortex, and were highest in the putamen, M1, and especially preSMA, with values close to 1 (Figure 3D, see Table S4).

Peak entropy, which approaches 1 when the peak times of a population uniformly tile the interval ^48^, followed the same gradient: it was low in phase-locked auditory clusters, high in core cluster 5 and belt cluster 6, and increased from parietal cortex through putamen and M1 to values close to 1 in preSMA (Figure 3E). In addition, the temporal sparsity reflects how concentrated population activity is across neurons at each moment in time, with larger values indicating that activity is dominated by fewer neurons and, consequently, less temporal overlap among neuronal responses. Temporal sparsity peaked in cluster 2 of the core and belt (Figure 3F) and did not coincide with stronger timing properties ^48^.

The recruitment fraction, the normalized time at which half of the neurons in a sequence reached their peak, was close to 0.5 for core cluster 5 and belt cluster 6, indicating a progression symmetrical to the timed interval, but close to zero for most other auditory clusters (Figure 3G). In parietal cortex, putamen, M1, and preSMA, clusters 1, 2, and 3 reached this point at about one third, one half, and two thirds of the interval, respectively, with preSMA cluster 2 lying exactly at 0.5. Thus, core cluster 5 and belt cluster 6 are the only auditory clusters that represent elapsed time between stimuli in relative terms, whereas preSMA neurons are recruited in rapid succession and flexibly fill the produced interval at all tempos.

Finally, the intrinsic timescale of baseline activity, estimated as in^48^ Murray et al., 2014, increased systematically along the circuit, from core and belt to parietal cortex and then to the putamen, M1, and preSMA (Figure 3H). The gradient in timing properties is thus accompanied by a gradient in intrinsic timescales.

## Auditory cortex contains local clocks for the interval between stimuli

Due to their timing properties, we next asked whether core cluster 5 and belt cluster 6 behave as population clocks, comparing them with the phase-locked core cluster 1 and with preSMA cluster 3. Core cluster 1 showed short-latency responses with a narrow peak-time distribution, consistent with classical phase-locked responses in AC (Figures 4A and 4B). In contrast, the richer peak-time distributions of core cluster 5 and belt cluster 6 were modulated by the TI (Figures 4A and 4B). Wasserstein distances between peak-time distributions increased with TI but changed little across serial orders (Figure 4C), indicating that these auditory clocks were renewed in every interval of the sequence. For both clusters, the Wasserstein distance grew monotonically with the difference between TIs, |Δ*TI*| (e.g., |Δ*TI*| = 300 ms for 850 vs. 550 ms; Figure 4D), whereas core cluster 1 carried no TI information; preSMA cluster 3 showed even larger distances as a function of |Δ*TI*|, together with strong temporal scaling (Figures 4E and 4F). Hence, auditory cortex is not a simple relay: it contains sparse sensory clocks whose peak times tile the interval between stimuli across tempos and regenerate with every interval.

**Figure 4.**
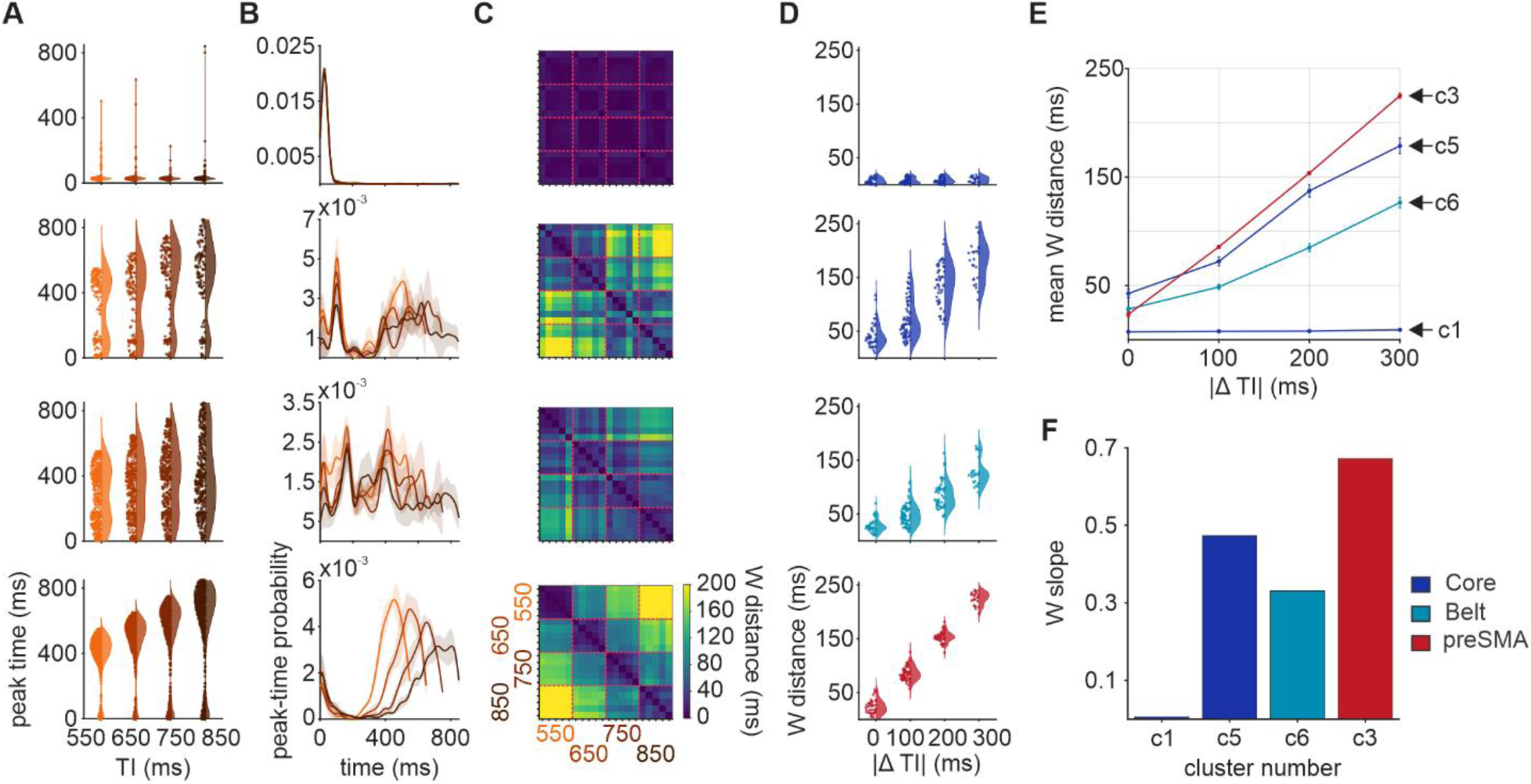
Target-interval-dependent organization of peak-time distributions across the audiomotor circuit. (A) Peak times of significant activity for the four TIs, pooled across serial orders, for core cluster 1 (C1), core cluster 5 (C5), belt cluster 6 (C6), and preSMA cluster 3 (C3). (B) Smoothed peak-time densities for each TI; thick lines and shaded regions show the mean ± SD across serial orders; densities were normalized to unit integral. (C) Pairwise Wasserstein-1 distance matrices between TI × serial-order peak-time distributions; red dashed lines delimit comparisons between serial orders within the same TI. (D) Distribution of Wasserstein distances as a function of the difference between TIs (|Δ*TI*|). (E) Mean Wasserstein distance as a function of |Δ*TI*| (error bars, SEM). (F) Slope of the relation between mean Wasserstein distance and |Δ*TI*|. Peak-time distributions differed from the pooled baseline for core C5 and preSMA C3 at all TIs, and for belt C6 at 550, 750, and 850 ms (Cramér–von Mises permutation tests, BH-FDR, *q* < 0.05). Wasserstein distances increased above the within-target baseline ((*W*0)) and the Wasserstein slopes were significant for core C5, belt C6, and preSMA C3 (*q* = 2.67 × 10^−4^; permutation tests with BH-FDR correction).

Nevertheless, the cells of core cluster 5 and belt cluster 6 are less than 25 percent of the responding cells in these areas and they are aligned to the auditory metronome suggesting that they form auditory chronometers for local computations.

## preSMA population activity forms a cone-shaped manifold

We projected the activity of the 980 preSMA neurons recorded during the PST onto its principal components (PCs). Consistent with previous work, PC1 and PC2 were sinusoidal with a period that increased with TI (Figures 5A and 5B), and in the PC1–PC2 plane the trajectories formed an elliptical loop for each produced interval that was reset at the tap (Figure 5E). Taps (green circles) fell on the top of the loops, where trajectories for all TIs converged as tangent circles; we propose that each passage through this zone generates a tick of the internal beat. The duration of the interval depended on both an increase in amplitude and a decrease in speed of the trajectories with TI during the dwell (Figures 5F and 5G), with amplitude changes slightly preceding speed changes (Supplementary Figure 11). Notably, PC3 had a magnitude that decreased with TI but varied little over time (Figures 5C and 5D), as expected for a tempo-specific control signal.

**Figure 5.**
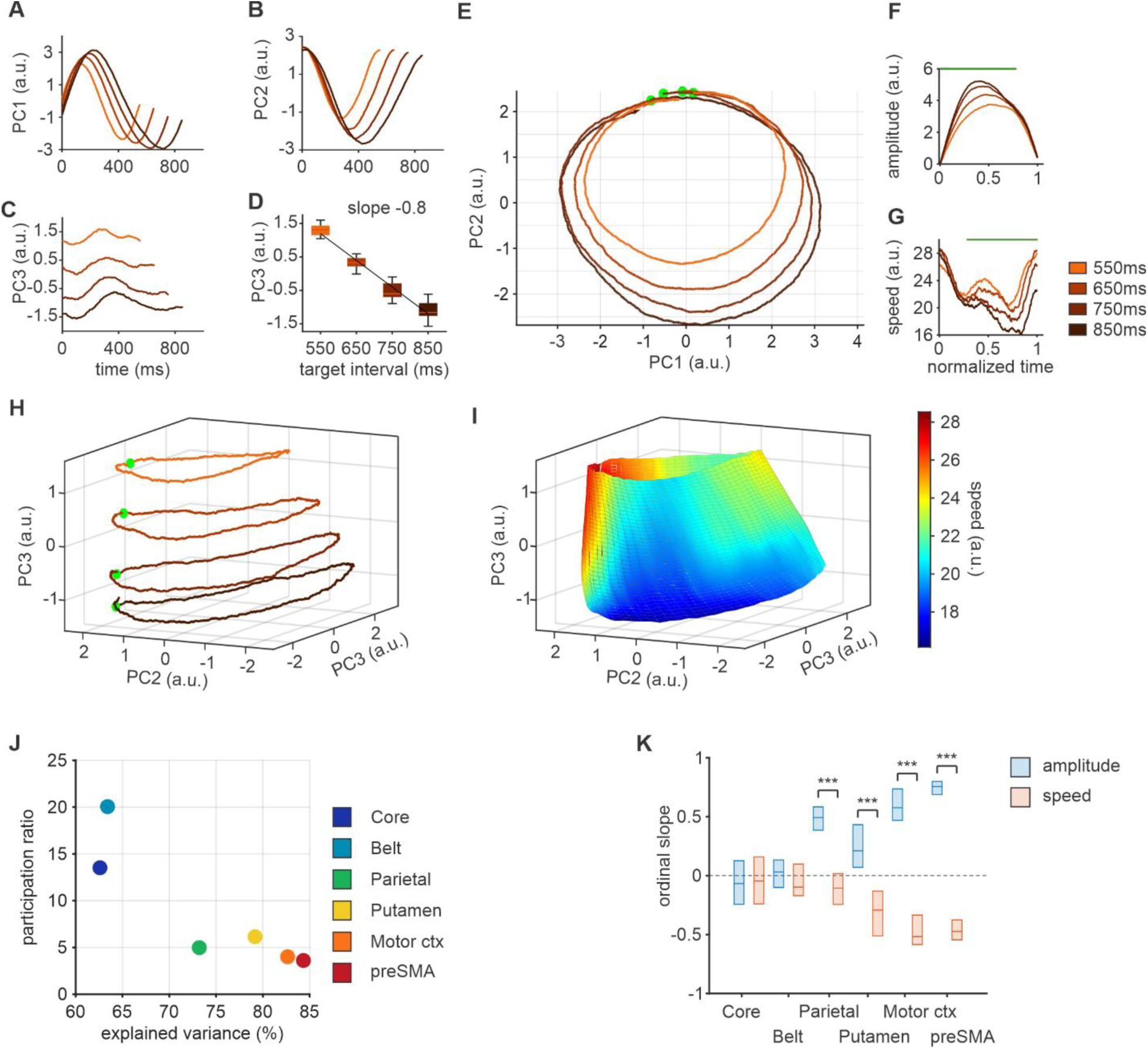
preSMA population trajectories form a cone-shaped manifold that encodes tempo. (A–C) First. (A), second (B), and third (C) principal components (PCs) as a function of produced time for the four TIs (color code on the right of G). (D) Interquartile boxplots of PC3 values as a function of TI, showing a large linear decrease across TIs. (E) preSMA population activity during the PST projected onto PC1 and PC2. Each produced interval corresponds to a loop whose radius increases with TI; taps (green circles) concentrate at the top, where trajectories for different TIs converge as tangent circles. (F and G) Amplitude (F) and speed (G) of the trajectories as a function of normalized interval. Amplitude increases and speed decreases with TI in the middle of the interval, during the dwell (significant time bins in green). (H) Projection onto the first three PCs; rings for different TIs lie at different heights along PC3. Conventions as in (E). (I) Manifold interpolated from the trajectories in (H), shaped as a truncated cone flattened around the tap, with a larger diameter for longer TIs during the dwell only. Surface color indicates trajectory speed (as in G), which is high near the tap and decreases with TI during the dwell. (J) Participation ratio as a function of cumulative explained variance for each area. Auditory areas show high dimensionality and low explained variance, whereas preSMA and motor cortex show low dimensionality and high explained variance. (K) Distributions of ordinal slopes of trajectory amplitude (blue) and speed (orange) as a function of TI across areas; positive values indicate increases with longer TIs. Slopes were tested against zero with sign-flip permutation tests with FDR correction. Amplitude slopes differed from zero in parietal cortex, putamen, M1, and preSMA, but not in core or belt, and speed slopes followed the same pattern. Asterisks indicate significant paired amplitude–speed differences (sign-flip permutation tests on within-area slope differences, FDR corrected across areas).

In the space of the first three PCs, the circular trajectories were stacked along PC3 by tempo, with the shortest TI on top and the longest at the bottom (Figure 5H). Interpolating between them by spatial lofting yielded a manifold shaped as a truncated cone, flattened around the tap, whose diameter increased with TI only during the dwell and whose speed was high near the tap and decreased with TI during the dwell (Figure 5I; Supplementary Figure 12). This manifold has all the properties to a population mechanism that governs beat timing. An ideal observer could read relative elapsed time from the angular position on the circular plane, and absolute time by integrating the speed and amplitude of the trajectory during the dwell. In addition, the height of the cone could thus act as a dynamic control signal that sets the amplitude and speed of the trajectory, and thereby the period, for each tempo.

We then compared population dynamics across areas. Measured with the participation ratio ^50^, auditory areas showed high dimensionality, but low total explained variance, whereas parietal cortex, putamen, M1, and preSMA showed low dimensionality and high explained variance (Figure 5J), in line with the number and diversity of their clusters. In core and belt, trajectories moved rapidly after each stimulus and then lingered in a small region of state space for the rest of the interval, with no evident geometry for encoding timing. In the other four areas, they formed elliptical loops with homogeneous densities for every produced interval (Supplementary Figure 13).

Trajectory amplitude increased and speed decreased with TI in preSMA and M1, with ordinal slopes significantly different from zero for both features and significantly different from each other (Figure 5K). Parietal cortex encoded TI only in amplitude and the putamen only in speed, whereas neither feature encoded TI in core or belt (Figure 5K). Thus, the low-dimensional population dynamics of preSMA and M1 encode the TI in their geometry and kinematics, whereas the high-dimensional dynamics of auditory cortex do not.

## Only the preSMA manifold represents tempo continuously

We reconstructed a surface from the three-dimensional trajectories of each area by spatial lofting (Figure 6A; Supplementary Figure 12). Core and belt did not form a coherent surface, whereas parietal cortex, putamen, M1, and preSMA formed truncated cones whose amplitude and speed changed with TI. We then tested three criteria of a two-dimensional manifold: smoothness (local planarity), embedding (absence of self-intersections), and regularity (linearly independent tangent directions). The putamen, M1, and preSMA met all three criteria, parietal cortex was intermediate, and core and belt failed (Figure 6B).

**Figure 6.**
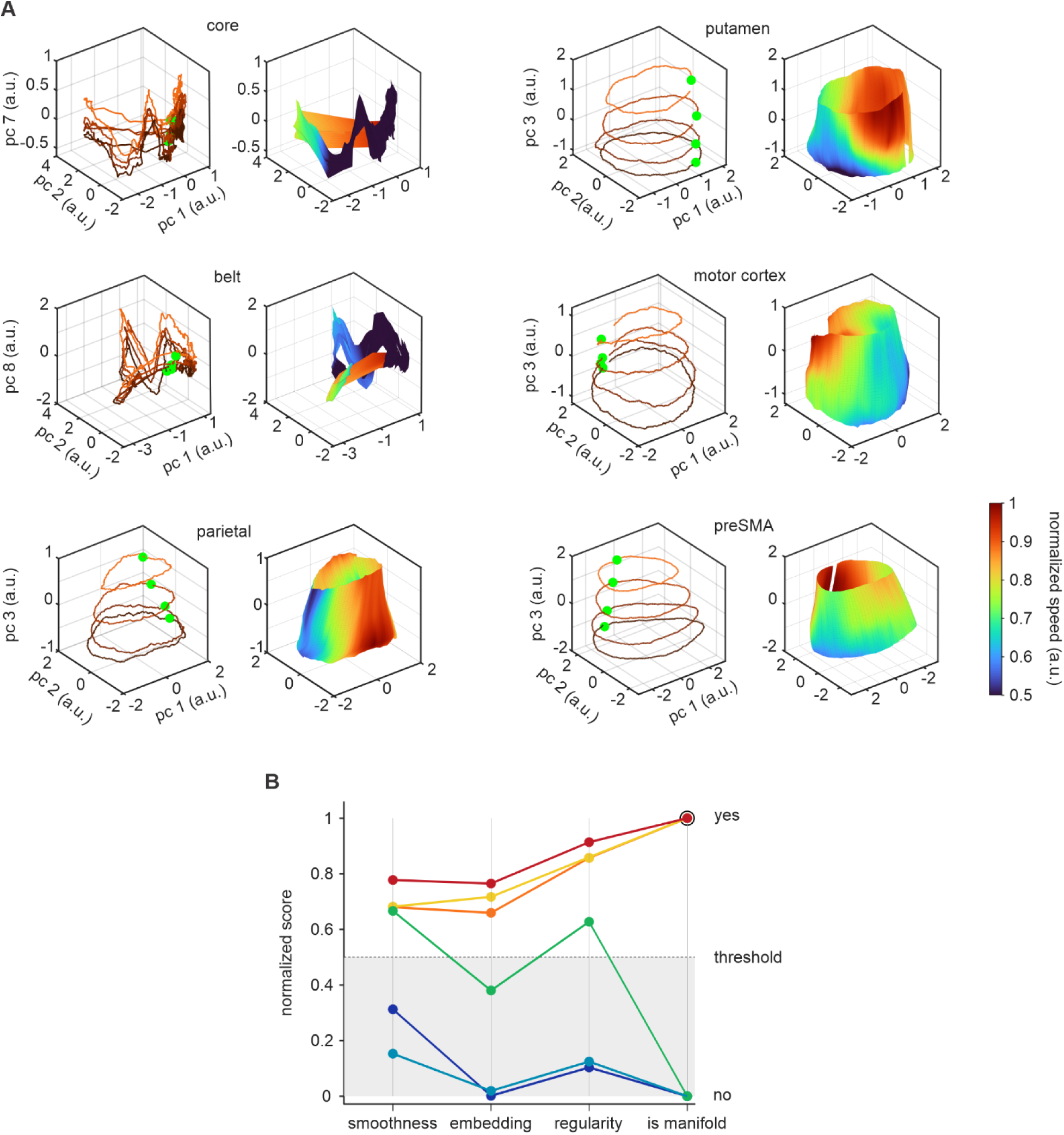
Manifold structure of population trajectories across areas. (A) Three-dimensional neural trajectories in the selected PC space (left) and the corresponding surfaces reconstructed by spatial lofting (right) for each area. Surface color indicates normalized trajectory speed; green denotes the starting point of each trajectory. (B) Manifold-compatibility scores (smoothness, embedding, and regularity) for each reconstructed surface. Scores above threshold indicate that a criterion is satisfied; a surface was classified as a two-dimensional manifold only if all three criteria were satisfied. Color code as in Figure 3D.

**Figure 7.**
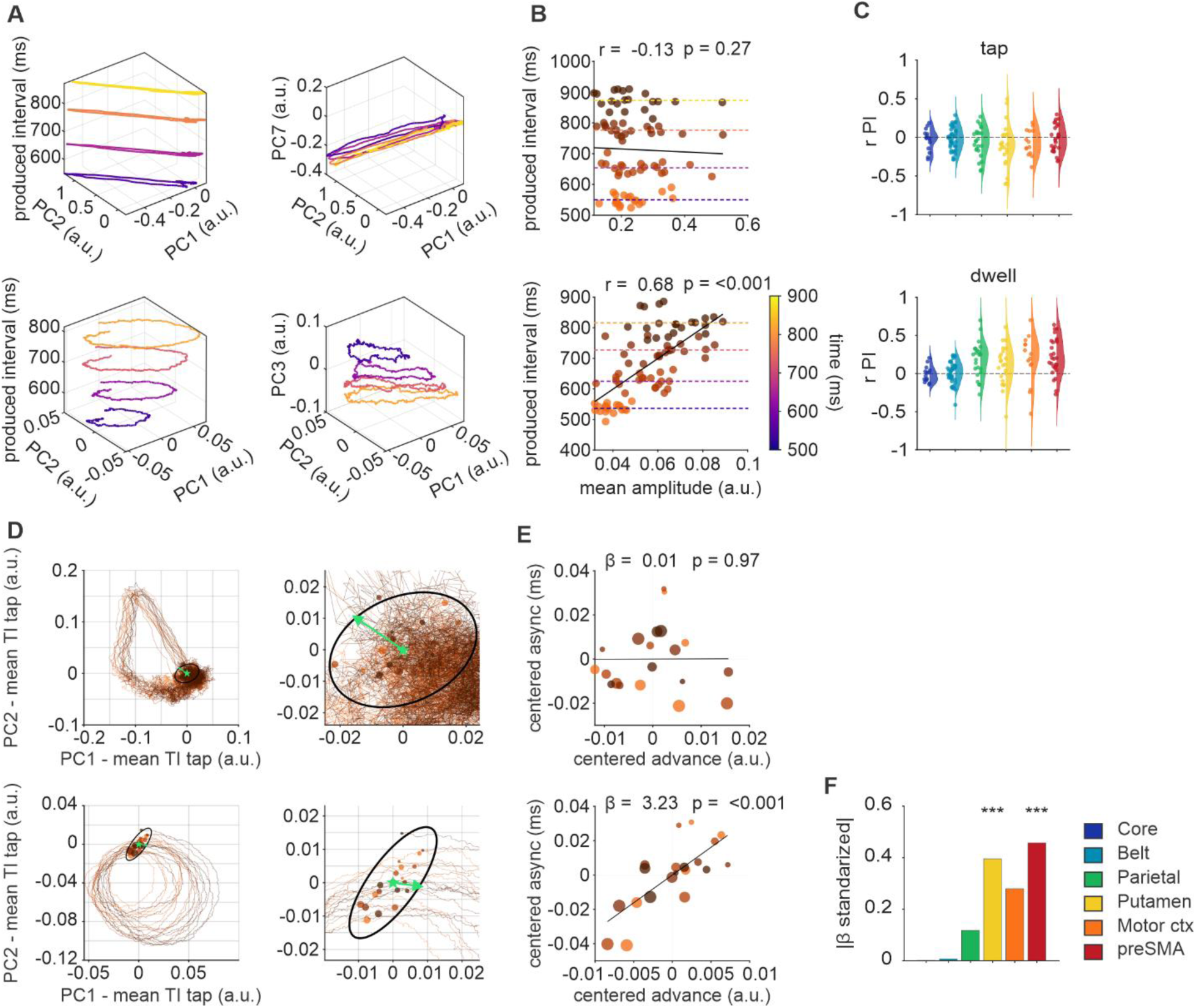
Trajectory geometry encodes the period and phase of tapping. (A) Mean trajectories from representative core (top) and preSMA (bottom) recordings, displayed in the PC1–PC2 plane with the mean produced interval along the vertical axis (left), and in PC1, PC2, and the PC that best separated the TIs (PC3 for preSMA; right). (B) Mean trajectory amplitude versus produced interval for the examples in (A). Each point is one trial, colored by TI (color code as in Figure 5G); dashed lines, mean produced interval for each TI; solid line, least-squares regression. (C) Distributions across sessions and areas (color code as in Figure 1E) of the correlation between trajectory amplitude and produced interval during the movement (top) and dwell (bottom) epochs. Parietal cortex, M1, and preSMA showed stronger coupling during the dwell (paired Wilcoxon test on |r|, p < 0.005); dwell correlations were significantly positive in parietal cortex, putamen, and preSMA (one-sample Wilcoxon test on signed r, p < 0.05). (D) Left, serial-order trajectories for core (top) and preSMA (bottom), centered for each TI on its mean tap position. Circles, tap positions; green star, centroid of the tap distribution; green arrow, estimated direction of progression; black ellipse, covariance of tap positions. Right, enlarged view of the tap region. (E) Centered tap advance versus centered asynchrony after removing the mean of each TI, for core (top) and preSMA (bottom). Points are serial orders within TIs, colored by TI and scaled in size by serial order; solid line, least-squares fit; β, within-condition regression coefficient. (F) Absolute standardized within-condition regression coefficient (|β|) relating centered tap advance to centered asynchrony for each area, after z-scoring both variables within each area. *p < 0.05, **p < 0.01, ***p < 0.001.

We next asked whether the height of the cone represents tempo as a continuous variable. Trial positions along PC3 were clearly ordered by TI in preSMA but blurred in parietal cortex, putamen, and M1 (Supplementary Figures 14A and 14B), and only preSMA showed a discriminability (d′) between neighboring TIs significantly above zero (Supplementary Figure 14C). When the manifold was rebuilt leaving out one intermediate TI (650 or 750 ms), the height of the new cone classified the held-out TI accurately in preSMA, whereas in parietal cortex, putamen, and M1 the jackknife confidence intervals crossed chance level (Supplementary Figure 14D). The rebuilt preSMA cone also showed the largest gain in predicting the amplitude and speed of the held-out rings in the PC1–PC2 plane (Supplementary Figures 14E and 14F). Consequently, only the preSMA manifold represents the TI tempo continuously, through the height of the cone and the amplitude and speed of its rings.

## preSMA trajectory geometry encodes the period and phase of each tap

If population geometry inherits behaviorally relevant information from the neurons that generate it, fluctuations in geometry should covary with fluctuations in the timing of taps ^34,51–53^. We therefore measured, in each recording session, the amplitude of the trajectories and the position of the tap along the direction of trajectory progression and related them to the produced interval (period) and to the asynchrony (phase).

Trajectory amplitude scaled with the produced interval across recording sessions (Figures 7A and 7B), and this coupling was stronger during the dwell than around the tap in parietal cortex, M1, and preSMA (Figure 7C). In contrast, trajectory speed showed no systematic relationship with the produced interval (Supplementary Figure 15). Thus, the amplitude of preSMA trajectories during the dwell encodes the duration of the produced interval, confirming previous findings ^33^.

The position of the tap along the trajectory changed with the asynchrony, namely, taps ahead of the centroid of the tap distribution were associated with positive asynchronies and taps behind it with negative asynchronies (Figures 7D and 7E). This association was significant in preSMA and the putamen but not in core, belt, parietal cortex, or M1 (Figure 7F). Therefore, the position of the tap within preSMA trajectories encodes the phase of synchronization.

## Timing errors are expressed in the geometry of the preSMA manifold

If manifold geometry underlies rhythmic timing, early and late errors should deviate from correct trials in opposite directions ^46,54,55^. We projected the population activity of error trials onto the state space defined by correct trials and examined the two intervals preceding an error (RSOE −2 and −1) and the error interval itself. preSMA activity was lower than in correct trials before early errors and higher before late errors, with an abrupt further change during the error interval (Figure 8A). Trajectories preserved their loop organization but contracted in the PC1–PC2 plane before early errors and expanded before late errors (Figure 8B), and along PC3 they drifted toward the positions of short TIs before early errors and of long TIs before late errors (Figure 8C). Again, among all areas, preSMA was the most tightly related to the duration of error intervals (Supplementary Figure 16A).

**Figure 8.**
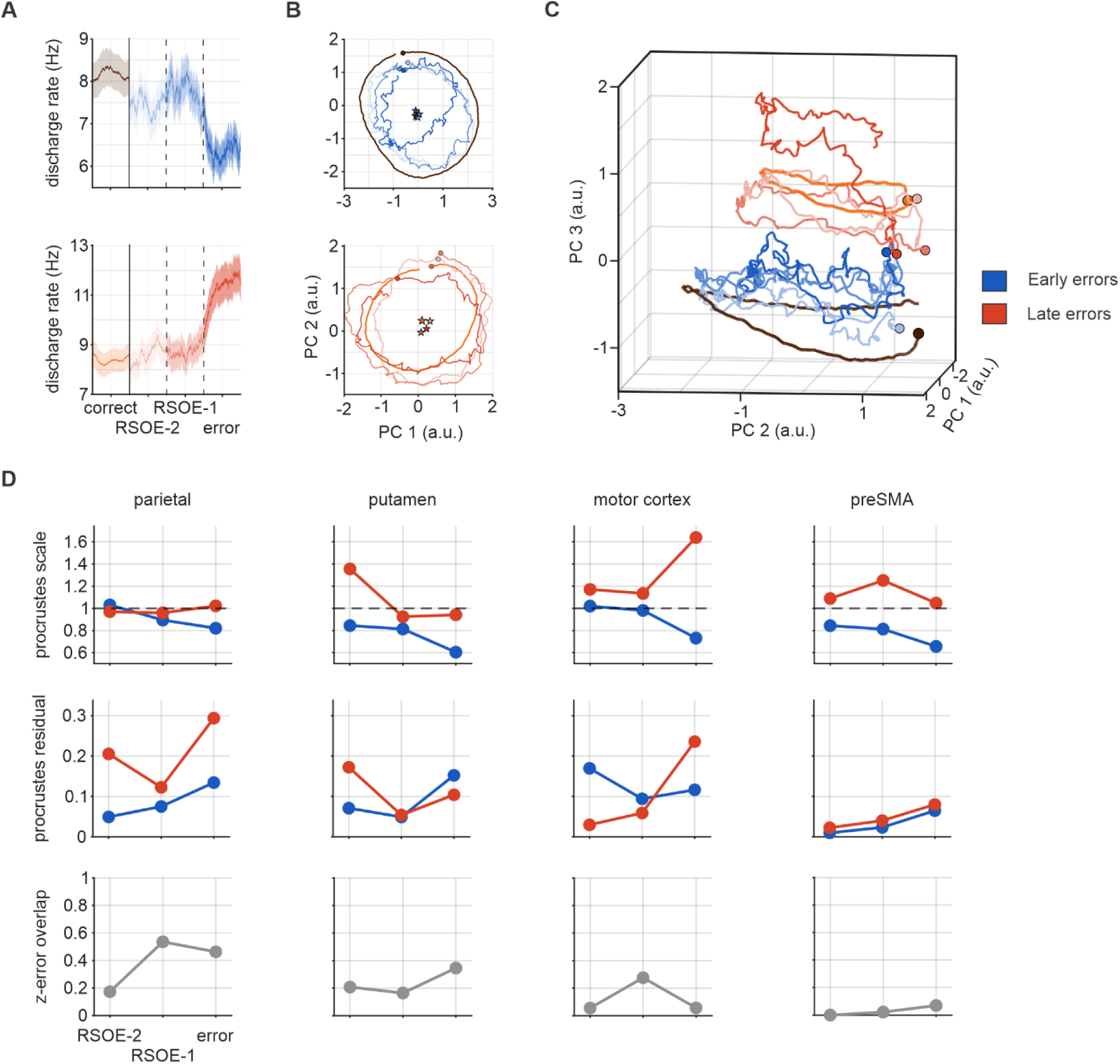
Timing errors are expressed in the geometry of population trajectories. (A) Mean population discharge rate for correct intervals (serial orders 3, 4, and 5), the two correct intervals preceding an error (RSOE −2 and RSOE −1; see Figure 2E), and the error interval, for early (top, blue) and late (bottom, red) errors; color code on the right of (B). (B) Corresponding trajectories projected onto PC1 and PC2, with the tap position (filled circle) and trajectory centroid (star). (C) Three-dimensional trajectories in the common PC space, where the third dimension corresponds to PC3. PCs were estimated exclusively from correct trials, and all error-related trajectories were projected onto the same basis using identical centering and normalization. (D) Geometric changes in parietal cortex, putamen, M1, and preSMA, expressed relative to the correct trajectory of the same population. Top, Procrustes scale factor between early or late errors and the corresponding correct trajectory; middle, residual Procrustes dissimilarity after optimal alignment; bottom, overlap coefficient between the normalized distributions of separator-axis values sampled along each trajectory (0, no overlap; 1, complete overlap). Color code as in (C).

We quantified these changes with Procrustes analysis (Figure 8D). This scale determines the contraction (below one) or dilation (above one) factor used to normalize the size between two geometric shapes without distorting their proportions. Here, we compare the shapes of the trajectories in correct trials with those in different RSOE periods of the incorrect trials. In preSMA, the Procrustes scale was below one for early errors and above one for late errors, both before and during the error interval, and residuals were small, indicating that the loops kept their shape but contracted or expanded relative to correct trials. Parietal cortex, putamen, and M1 showed larger residuals and no systematic scaling, except during the error interval in M1. preSMA was also the only area in which early and late errors remained separated along PC3, with the lowest overlap across the pre-error and error intervals (Figure 8D; Supplementary Figure 16B).

Therefore, preSMA trajectories encode the period of the produced intervals in both correct and error trials, through the size of the loops in the PC1–PC2 plane and their position along PC3, and these signatures emerge in the intervals that precede an error. Parietal cortex, putamen, and M1 carry only part of this information, supporting the idea of a hierarchical organization of the beat clock.

## Discussion

We recorded neurons across six nodes of the macaque audiomotor circuit during beat synchronization and found that the timing properties of single neurons and populations increase gradually from auditory cortex to preSMA. Auditory cortex detected the metronome in a context-dependent manner and contained local clocks for the interval between stimuli; parietal cortex showed intermediate properties; and the putamen, M1, and especially preSMA generated neural sequences that rescaled with tempo. Only in preSMA did population activity form a cone-shaped manifold that jointly encoded the tempo of the metronome, the produced interval, and the phase of each tap, and whose geometry shifted in a predictable direction before and during timing errors. These results support a hierarchical beat clock with preSMA at its top.

## Monkeys set the period and control the phase of synchronization

The monkeys set the period of the produced intervals at the beginning of the trial and adjusted the phase of each tap to maintain synchrony, a strategy probably shaped by our reward rule, which penalized large asynchronies^20^. Earlier versions of the task rewarded accurate produced intervals regardless of the asynchrony and yielded precise periods but positive asynchronies ^19^, which were interpreted as a lack of prediction ^56^. When reward depends on the asynchrony, as here, monkeys synchronize with asynchronies close to zero across a wide range of tempos ^20,22^. Monkeys, like humans, can thus flexibly and predictively control the period or the phase of tapping ^36,57^, but the reward rules determine which of the two parameters they emphasize^4,58^.

## Auditory cortex combines context-dependent detection with local interval clocks

Almost all auditory clusters were tonically inhibited during synchronization relative to perception, while their phase-locked responses were preserved. We propose that this tonic inhibition, probably coming from the premotor system^59,60^, suppresses ongoing activity and increases the signal-to-noise ratio of the responses to the metronome, an ‘iceberg effect’ that could be mediated by local inhibitory circuits^59,61,62^. In humans, moving in time with a periodic stimulus improves the segmentation of sounds and the discrimination of events on the beat^17,63^,more so when motor predictions are better^64^. Phase-locked responses were also larger during synchronization than during passive listening, possibly through attentional gain^17,65,66^, as described for attention to rhythmic sounds in primate auditory cortex^67^ and in humans^15,68,69^.

Beyond detecting the stimuli, core cluster 5 and belt cluster 6 formed neural sequences whose peak activations tiled the interval at every tempo, so that a reader could measure relative elapsed time by identifying the small group of active neurons ^70^. These sequences were robust across trial repetitions and regenerated in every interval, but only during synchronization and not during passive listening ^71^. They extend evidence for sensory clocks in auditory and visual areas ^72–75^ by showing that, within auditory cortex, interval clocks coexist with context-dependent detection of the metronome. Their sparse responses, however, made them less accurate than the scaling sequences of preSMA and probably are used for local auditory elapsed time prediction. Finally, tap-aligned clusters in core and belt could carry a top-down beat signal from the frontal lobe ^76–78^ or tactile and proprioceptive feedback from the tap ^79^; together with stimulus-locked responses, they could support error correction by comparing the internal beat, the actual tap, the auditory input and the stimuli elapsed time quantification.

## A gradient of timing properties along the audiomotor circuit

Temporal scaling, interval tiling, and symmetric recruitment increased from auditory to parietal cortex and peaked in preSMA, whose neural sequences tiled the interval almost perfectly, regenerated in each produced interval, and rescaled across tempos ^34,35,80^. Parietal cortex behaved as a timing interface between auditory and motor areas ^81^, whereas the putamen ^82–84^ and motor cortex^85,86^ were functionally closer to preSMA without matching its timing properties. The passive condition corroborated this hierarchy: auditory areas processed the metronome in both conditions, parietal neurons responded mildly in both, and frontal areas responded only when the metronome guided behavior. Likewise, motor areas of monkeys do not respond to passively heard tone sequences ^87^, but they do when the tones are paired with reward ^87^ or used to predictively drive movement ^88,89^. These results invite a reexamination of the origin of event-related potentials, such as the mismatch negativity, that are elicited by violations of beat regularity in preattentive listeners ^56,90^.

## preSMA trajectories encode the period and phase of the beat

We characterized the geometric properties of the neural trajectories that parametrically encode the TIs, the period of produced intervals, the phase of the asynchronies, as well as the duration of intervals in correct and early and late error trials. Again, we found that preSMA possesses all the properties of the core timing neural chronometer with a tight representation of period and phase in its population dynamics and with a gradual dissipation in the association between neural population responses and rhythmic behavior in the other five areas. These findings support the notion that the medial premotor areas constitute the core timing circuit, using a neural population clock to measure time and maintain the internal beat during rhythmic timing ^30,91^. As previously documented^31,33,34^, preSMA trajectories form a regenerating loop for each produced interval, and taps occur where the loops for all tempos converge^34–36^. Here we show that the position of the tap relative to this zone encodes the asynchrony, providing a mechanism for the tap-by-tap control of phase. Amplitude and speed both changed with tempo, but across sessions only amplitude tracked the produced interval, especially during the dwell. Because a second speed peak near the next tap acts as a control signal that triggers the stereotyped movement^33^, trajectory speed may instead trigger the tapping motor program in motor cortex, in line with the relation between neural trajectory speed in motor cortex and movement speed ^92–94^.

## A cone-shaped manifold for beat timing in preSMA

A key finding is a third neural dimension, PC3, that together with the circular plane defines a two-dimensional manifold shaped as a truncated cone. This manifold is a low dimensional robust mechanism for beat timing, whose properties explain the monkeys’ strategy to solve the task. The angular position on the cone describes the evolution of the population state within each interval and the phase of each tap, whereas the height of the cone sets the size of the circular cross section during the dwell and could act as a parametric control signal for tempo throughout the trial. A similar manifold underlies the reproduction of single intervals in the medial premotor cortex ^47^, where duration is encoded mainly by speed rather than amplitude ^52,95^, and inputs from the motor thalamus can set the speed at which the clock traverses the plane ^28,47^, giving it flexibility to generalize across durations and tasks ^42^. In our data, the tempo control signal was not evident in parietal cortex, putamen, or M1, suggesting that it is integrated within preSMA circuits; recordings in the motor thalamus will be needed to determine its origin. Previously we showed that MPC trajectories gradually increase in amplitude and oscillatory strength during the perception epoch and then shift into a distinct subspace at the onset of tapping^31^. Hence, the height of the cone is probably set while the monkey attends to the metronome with no movement.

Error trajectories remained organized on the manifold: the size of the cross section and the position along the cone shifted toward shorter or longer tempos before and during early and late errors. Correct and erroneous timing are thus organized within a single low-dimensional space, in which deviations in manifold position are accompanied by predictable deviations in produced timing. Only preSMA contained a manifold that captured all these behavioral parameters, supporting its role as the core timing integrator for beat synchronization ^34,96–98^. The graded organization of the other areas is consistent with a flow of information in which the preSMA beat signal is compared with incoming auditory input to adapt the phase of the taps and is sent to the corticospinal system to trigger movement. A similar hierarchy has been observed during visual interval categorization in humans ^73^, in which preSMA prefers durations close to the category boundary ^99^, suggesting that different timing tasks engage hierarchical organizations with preSMA at the top.

Neural manifolds can thus provide a low-dimensional architecture in which behaviorally relevant variables are jointly organized ^100,101^: in preSMA, the period and phase of rhythmic tapping are complementary coordinates of a common population state space.

## Limitations of the study

Our conclusions rest on two monkeys and on four tempos (550–850 ms). Whether the manifold extends to slower or faster tempos, to tapping without a metronome, or to complex musical rhythms remains to be tested. Population analyses pooled neurons recorded across sessions and monkeys, so trajectories were estimated from trial-averaged activity rather than from simultaneously recorded single trials, and error-related geometry was characterized for families of error trials rather than for individual trials. The relation between manifold geometry and behavior is correlational; causal manipulations of preSMA, and of candidate inputs such as the motor thalamus, are needed to establish whether the manifold controls synchronization. Finally, the reward rule emphasized asynchrony, which may have biased the monkeys toward phase control.

## Acknowledgments

We thank Ranulfo Romo, Bruno Averbeck, Luis Carrillo Reid, and Victor de Lafuente for their fruitful comments on the manuscript. We also thank Maria Antonieta Carbajo, Juan Ortíz, and Raul Paulín for their technical assistance. The scientific English of this manuscript was proofread by Claude.

Pablo Marquez is a doctoral student of the Programa de Doctorado en Ciencias Biomédicas at Universidad Nacional Autónoma de México (UNAM) and has received a fellowship (No. 403878) from Secretaría de Ciencias, Humanidades, Tecnología e Innovación (SECIHTI, formerly CONAHCYT).

Hugo Merchant is supported by Secretaría de Ciencias, Humanidades, Tecnología e Innovación (SECIHTI): CBF-2025-G-89, and Programa de Apoyo a Proyectos de Investigación e Innovación Tecnológica (UNAM-DGAPA-PAPIIT): IG200424.

## Data availability

The data used to generate the associated figures will be made available upon request.

## Code availability

The code used to generate the associated figures will be made available upon request.

## EXPERIMENTAL MODEL AND SUBJECT DETAILS

### Animals

Two male rhesus monkeys (*Macaca mulatta*) were used in this study: Monkey 1 (13 years old, approximately 8.5 kg) and Monkey 2 (9 years old, approximately 7.5 kg). The monkeys were obtained from a specialized *Macaca mulatta* breeding company in Mexico City, called Proyecto Camina A.C., which adheres to international standards of reproduction and animal care. The company does not catch animals in the wild and has certified veterinary care.

All animal care, housing, and experimental procedures were approved by the Ethics in Research Committee of the Instituto de Neurobiología, UNAM, protocol 090.A. All experiments followed the 3Rs and conformed to the principles outlined in the Guide for the Care and Use of Laboratory Animals (NIH, publication number 85-23, revised 1985).

The animals were housed in a monkey facility with cages measuring 1.5 m³. They were kept under controlled temperature (22 °C) and humidity, and a 12–12 h day-night cycle. The monkeys were housed separately since they did not show amicable interactions. Our Institute has three certified veterinarians who continuously provide care for the monkeys and perform regular health checkups and medical examinations. Animal care staff keep the facilities clean and provide food for the animals every day of the year. In addition, monkeys were monitored daily by researchers and the animal care staff to check their health and welfare^102^.

The monkeys were fed ad libitum with a special diet from LabDiet (Monkey Diet 5038) and had daily access to raisins and berries. Water was provided five days a week during correct trials in the training and experimental sessions. Monkeys received between 200 and 250 ml of water and/or juice per day. If performance was especially poor or the monkey did not engage much with the task, additional water to reach the minimum 200 ml was provided by soaking dry food pellets in the required amount of water. Following the last session of the week, monkeys were given up to 350 ml of water. The monkeys also received fruit, such as apples and bananas, after experimental sessions and on weekends.

## METHOD DETAILS

### Perception synchronization task

Monkeys were trained in the perception-synchronization task (PST) using operant conditioning techniques^19,24^. During the PST, monkeys were presented with isochronous metronomes composed of brief white-noise bursts (33 ms total duration, including 10-ms cosine onset and offset ramps; 75 dB SPL). The stimuli defined target intervals (TIs) of 550, 650, 750, or 850 ms. The TI remained constant within each trial and was varied pseudorandomly across trials. Each trial consisted of a train of 11 auditory stimuli and 7 movements (taps) produced by the monkey. The monkey initiated each trial by placing its hand on a lever (hold period). A yellow square then appeared on a screen in front of the animal, and the auditory stimuli began at the same time.

The animal was required to keep holding the lever during the first four stimuli (perception epoch). The yellow square then disappeared, and this served as the go signal for the monkey to begin tapping in synchrony with the remaining seven stimuli (synchronization epoch).

The monkeys were seated in a primate chair within a custom setup designed to deliver visual and auditory stimuli and to record motor responses. The non-dominant hand was restrained. Auditory stimuli were delivered through two speakers placed to the left and right in front of the monkey, equidistant from the animal’s head (Figure 1 A,B). Taps were detected when the monkey’s hand crossed a laser sensor while touching an acrylic surface, which eliminated auditory feedback from the movements.

Throughout each trial, the asynchrony between each tap and its corresponding stimulus (paired by serial order) was measured. If the asynchrony remained within a 280 ms threshold for all seven taps, the trial was considered correct and the monkey received a water reward (inter trial interval duration of 2s). If the asynchrony exceeded this threshold at any serial position, the trial was aborted without reward, and the animal had to initiate a new trial.

We also presented a passive perception task where the monkeys listen to the same metronomes used during the PST without access to the key holding, the tapping zone or the reward tube. The beginning of each trial was automatically triggered (2s inter-trial time).

Behavioral control and stimulus delivery were implemented using an RZ6 Multi-I/O Processor (Synapse version 94; Tucker-Davis Technologies, Alachua, FL, USA). The processor was controlled by custom-written programs in MATLAB R2016b (version 9.1.0.441655; MathWorks, Natick, MA, USA) using the Psychophysics Toolbox (Psychtoolbox-3, version 3.0.18)^103,104^.

### Behavioral measures

For the behavioral analysis, the following measures were computed from correct trials, excluding the first tap of each trial: (1) asynchrony, defined as the time difference between each tap and its corresponding stimulus, matched by serial order (tap minus stimulus; negative values indicate that the tap preceded the stimulus); (2) produced interval, defined as the time between two consecutive taps; and (3) constant error, defined as the produced interval minus the target interval. All measures are reported in milliseconds.

Trials were classified as errors when the absolute asynchrony exceeded 280 ms at any serial position: early errors when the asynchrony was below −280 ms and late errors when it exceeded +280 ms. Early errors were analyzed at the 850-ms TI and late errors at the 550-ms TI, where each type was most frequent. Error trials in which the error occurred at serial orders 5–7 were aligned to the error tap (relative serial order to error, RSOE = 0), and the asynchronies from RSOE −3 to 0 were analyzed. As a reference, each correct trial at the same TI contributed four consecutive asynchronies ending at serial order 5, 6, or 7; correct trials were randomly assigned to these end positions in proportion to the frequency of errors at each position (random number generator seed: 42).

### Behavioral variance decomposition

Produced intervals and asynchronies were analyzed separately, using correct trials and excluding the first tap of each trial (five produced intervals and six asynchronies per trial). Each session was included only once, even when it was recorded simultaneously in several areas (n = 220 unique sessions from both monkeys). For each session, the data of each TI were arranged as a trial × serial-order matrix, and each observation was decomposed as

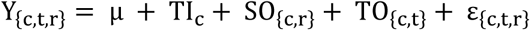

where *Y*_{*c*,*t*,*r*}_ is the value at serial position r of trial t in TI c; μ is the session grand mean; *TI_c_* is the deviation of the mean of TI c from μ; *SO*_{*c*,*r*}_, the serial-order (SO) profile, is the mean of serial position r minus the TI mean; *TO*_{*c*,*t*}_, the trial offset, is the mean of trial t minus the TI mean; and *ε*_{*c*,*t*,*r*}_ is the residual.

Within each TI, the variance components of SO, trial offset, and residual were estimated from expected mean squares, treating trials as random and serial positions as fixed: 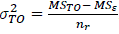, and 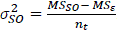, where MS is the mean square of each source, and *n* and *n_t_* are the numbers of serial positions and trials; negative estimates were set to zero. These components were averaged across TIs, weighted by the number of observations per TI. The TI component was computed as the variance of the TI means around μ, weighted by the number of observations per TI. Each component was expressed as a fraction of the sum of the four components and averaged across sessions.

To identify the sources of serial dependence, the lag-1 autocorrelation (ac1) between consecutive serial positions was computed for each TI, pooling all trials, and averaged across TIs. It was computed on the raw data and after successively removing the TI component, the SO profile, and the trial offset (that is, on the residual), and then averaged across sessions.

Because each row of the residual sums to zero by construction, the expected ac1 of a residual without serial dependence is 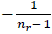, (−0.25 for produced interval and −0.20 for asynchrony), which is shown as the theoretical null.

### Recursive model of asynchrony correction

Tap-to-tap correction during synchronization was quantified separately for each TI using correct trials from the 220 unique sessions of both monkeys. The terminal transition was also excluded, because it showed an edge-related increase in variability and a shift in mean asynchrony, leaving five taps per trial.

Asynchrony obeys the exact identity

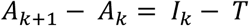

where *A_k_* is the asynchrony at tap k, *I_k_* is the interval produced between taps k and k+1, and T is the TI. This identity was verified as an extraction-integrity check. Corrective behavior was modeled as

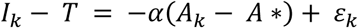

which yields

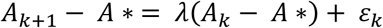

with *λ* = 1 − *α*. Thus, λ represents the fraction of the current deviation carried into the next tap, whereas α represents the corresponding apparent correction gain.

For each TI, the empirical recursion

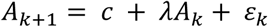

was fit by ordinary least squares to all valid (*A_k_*, *A_k_*_+1_) pairs pooled across trials and serial positions. The equilibrium asynchrony was defined as the fixed point

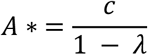

corresponding to the asynchrony for which the expected subsequent asynchrony is unchanged; for |λ| < 1, this point is stable.

To assess whether the fitted tap-to-tap dynamics accounted for the serial evolution of behavior, the recursion was iterated from the observed mean asynchrony at the first retained tap, _Â*k* = *A* ∗ + *λ*_(*k*−1)(Ā1− *A*∗).

Because serial order was not used to estimate λ or A*, the predicted profile introduced no free parameters beyond the initial anchor.

### Neural recordings

#### Recording sites and probes

Neural activity was recorded using high-density silicon microelectrode arrays from five cortical regions (auditory core, auditory belt, presupplementary motor area (preSMA), parietal area 7a, and primary motor cortex) and one subcortical structure (putamen). Five sites were recorded simultaneously in each session of the main recording phase: auditory core, auditory belt, preSMA, parietal area 7a, and a fifth site that alternated across sessions between M1 and putamen. Probe models are listed in the key resources table.

Monkey 1 (107 sessions). Auditory mapping was performed with acute recordings using a 64-channel NeuroNexus probe (V1×64-Edge), while preSMA was recorded simultaneously with a semi-chronic 64-channel NeuroNexus probe (A8×8-Edge; 2022–2023). In the main recording phase (2024), auditory core and belt were recorded with semi-chronic 32-channel NeuroNexus probes (V1×32-Edge), and preSMA and parietal area 7a with semi-chronic 64-channel NeuroNexus probes (Buzsaki64). The fifth site, M1 or putamen, was recorded acutely with a 64-channel NeuroNexus probe (V1×64-Edge).

Monkey 2 (113 sessions). Auditory mapping was performed with acute recordings using a 32-channel NeuroNexus probe (V1×32-Poly2; 2020). Auditory core was recorded with a chronic 64-channel Cambridge NeuroTech probe (ASSY-156-M2; 2020), and auditory belt, preSMA, and parietal area 7a with semi-chronic 64-channel NeuroNexus probes (Buzsaki64; 2020–2021). The fifth site, M1 or putamen, was recorded acutely with a 32-channel NeuroNexus probe (V1×32-Poly2).

#### Signal acquisition

Across experiments, chronic, semi-chronic, and acute recordings were combined. Signals were acquired with two Blackrock Neurotech systems, both using CerePlex M headstages (128 channels each): a Cerebus Neural Signal Processor (128 channels, NSP1.0 with Experiment I/O) running Central Software Suite v7.0.4 (NSP firmware v7.0.0; Hardware Library v3.11), and a CerePlex Direct data acquisition system (256-channel configuration with Experiment I/O) running Central Software Suite v7.0.5 (NSP firmware v7.00.05.00; configuration protocol v3.11). In both systems, continuous broadband signals were digitized at 30 kHz. For online visualization and detection of spiking activity, the acquisition configuration applied a bandpass with a high-pass cutoff of 300 Hz (order 1) and a low-pass cutoff of 7.5 kHz (order 3). Online sorted spike waveforms were stored with a length of 48 samples and a pre-trigger of 10 samples.

### Spike sorting

The data were automatically spike sorted using Kilosort 2.5 (v2.5)^105,106^ and then manually curated using phy (v2.0b6; https://github.com/cortex-lab/phy). For each recording session, the raw extracellular signals from the synchronization task and the passive perception task were concatenated prior to spike sorting to facilitate the identification of units shared across both behavioral conditions. During manual curation, each unit detected by a given template was visually inspected by evaluating the associated voltage waveform/template across channels, and units whose events were consistent with artifacts or noise were discarded. Units exhibiting low-amplitude spikes, unstable or inconsistent waveform shapes over time (drift-related changes), and/or refractory period contamination were classified as multi-unit activity and excluded from subsequent analyses. Finally, candidate units that were spatially adjacent and displayed highly similar waveforms were systematically compared to determine whether they should be merged, using waveform similarity and the absence of refractory contamination in the cross-correlograms between units as decision criteria. Units meeting these criteria were considered to reflect the spiking activity of a single neuron. This procedure yielded 2,918 single units in monkey 1 (auditory core, 255; auditory belt, 644; parietal area 7a, 778; putamen, 503; M1, 68; preSMA, 670) and 1,801 in monkey 2 (auditory core, 206; auditory belt, 589; parietal area 7a, 224; putamen, 215; M1, 257; preSMA, 310).

### Firing rate estimation

Instantaneous firing rates were estimated by convolving each neuron’s spike train with a causal exponential kernel and sampling the result on a time grid with 1 ms resolution (Δt = 0.001 s). Firing rates were reported in spikes/s.

For each neuron and each time sample *t_i_*, the firing rate was computed as the sum of exponentially decaying contributions from spikes *s_j_* that occurred at or before *t_i_* (causality). The kernel was defined as:

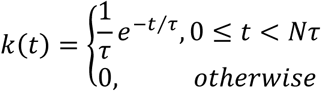

and the firing rate at time *t_i_* was:

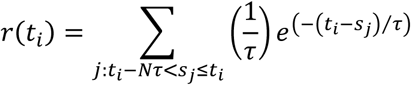

To limit the number of effective time constants contributing after each spike, we truncated the causal exponential kernel to a finite window of Nτ, with N = 4; thus, only spikes within (*t_i_* − *Nτ*, *t_i_*] contributed to *r*(*t_i_*). After 4τ, a spike contributes only 1.8% (*e*^−4^) of its initial value, so its contribution was considered negligible. Different time constants were used depending on the recorded area: auditory areas used τ = 10 ms, whereas all other areas used τ = 30 ms.

### Trial Time Warping

To determine whether the activity of each neuron during the synchronization epoch was better aligned to the sensory events (stimuli) or to the motor events (taps), we adapted the trial time-warping method that we previously proposed^45^. The analysis was performed separately for each monkey using correct trials only. For each neuron and TI, we used the stimulus-aligned spike trains, together with the stimulus and tap times of each trial, within a window extending from 500 ms before the first stimulus to 500 ms after the last (seventh) stimulus of the synchronization epoch. The number of trials was equalized across TIs by keeping, for each TI, only the first trials up to the smallest number of trials available across TIs for that neuron. Neurons were included when their mean firing rate in this window was at least 0.5 spikes/s (0.4 spikes/s in Parietal and 0.3 spikes/s in Putamen) and below 100 spikes/s, and when the coefficient of variation of their firing rate across trials (SD across trials, averaged across TIs, divided by the mean firing rate) was below 0.75.

The warping transformation mapped the spike times of each trial onto a common template. First, the spike times between consecutive taps were linearly rescaled so that each tap fell at the time of its stimulus plus the mean asynchrony of the monkey, while the window edges remained fixed; thus, each produced interval was rescaled to the TI. The warped spike times were then defined as a weighted combination of the rescaled and the original spike times, controlled by the parameter *w*. With *w* = 0 the activity remained aligned to the stimuli, with *w* = 1 it was aligned to the taps, and intermediate values produced alignments between the two. We evaluated *w* from 0 to 1 in steps of 0.1.

For each value of *w* and each TI, we estimated the firing rate as the average across trials of the warped spike trains. The spike train of each trial was modeled as an inhomogeneous Poisson process with this rate, which was estimated from all trials, including the evaluated. The log-likelihood of *w* was the sum of the trial log-likelihoods across all trials and TIs, so that each neuron had a single best alignment. The log-likelihood, evaluated at the 11 values of *w*, was interpolated with a cubic spline at a resolution of 0.01, and *ŵ* was the value of *w* that maximized the interpolated log-likelihood.

Finally, we compared three alignment categories: sensory, motor, and complex. The evidence for the sensory and motor categories was the likelihood at *w* = 0 and *w* = 1, respectively, and the evidence for the complex category was the geometric mean of the likelihood across *w*. We computed the three pairwise ratios between these evidences on a log₁₀ scale. A neuron was assigned to a category when its evidence exceeded that of each of the other two categories by at least 2 log₁₀ units (a factor of 100); otherwise, it was considered indeterminate.

Each included neuron was counted as motor when *ŵ* > 0.5 and as sensory when *ŵ* < 0.5, and an area was considered motor when more than half of its neurons were motor (Supplementary Figure 4). Core and Belt were classified as sensory, and preSMA, Parietal, Putamen, and M1 as motor; accordingly, the activity of the four motor areas during the synchronization epoch was aligned to the taps for the PSTH analysis (see Event-aligned PSTHs per target interval).

### PSTH construction

To compute the PSTH (peri-stimulus time histogram), we extracted, for each neuron, the spikes occurring within each presented interval or within each produced interval. For the Core and Belt areas, neural activity was aligned to stimulus onset. The computation was performed separately for each task epoch. During the perception epoch, we used the three presented intervals delimited by the four perception stimuli. During the synchronization epoch, we used the presented intervals 2–6, excluding the first presented interval. For the control activity, we extracted spikes from the 1 s period preceding the first stimulus. For the passive condition, we used all presented intervals (1–10) from the activity aligned to the presented stimuli.

For the remaining areas, the main difference was that, during the synchronization epoch, spikes were aligned to movement execution using time warping transformation. After selecting the appropriate alignment and identifying the relevant events for each epoch, we computed (across all trials and for each serial order) the spike times relative to the corresponding event. These spike times were then mapped onto the interval-specific window *T*, defined as the time to the next event: *T* corresponded to the time to the next stimulus for stimulus-aligned analyses, and to the time to the next movement for movement-aligned analyses. This yielded relative spike times between 0 and *T*. Importantly, this procedure was performed separately for each target interval duration (550, 650, 750, and 850 ms), thus obtaining a distinct PSTH for each interval.

We then estimated the smoothed firing rate (see Firing-rate estimation). To avoid an edge effect of the causal kernel at the onset of each window, where the firing rate would otherwise start at zero because no preceding spikes were included, the firing rate was computed including the spikes from the 500 ms preceding the window onset, and only the samples within *T* were retained. For time-warped data, this pre-window was defined on the rescaled time axis.

Because the kernel was truncated at 4τ (at most 120 ms), this pre-window included all spikes contributing to the firing rate at the window onset. Finally, for each neuron and for each epoch/condition (control, perception, synchronization, and passive), we averaged the firing-rate profiles across all serial orders and trials, obtaining a representative firing-rate mean trajectory for the time interval between the two relevant events for each of the four interval durations.

### Estimation of activation periods

Starting from the original set of spikes used to compute the PSTH of each neuron, we generated a null distribution by randomly reassigning the time stamp of every spike within the analyzed interval window of duration T, at 1-ms resolution. Concretely, for each of the N spikes, a new time was sampled from a uniform distribution over the interval [0, T] (i.e., over the exact time window used to compute the PSTH), thereby preserving the total spike count while removing any time-locked temporal structure. The permuted spike train was then convolved with the same exponential kernel used for the original firing-rate estimate to obtain a smoothed firing-rate trace. To avoid the edge effect of the causal kernel, this trace was computed with the same correction as the observed firing rate, that is, including the spikes from the 500 ms preceding the window onset and retaining only the samples within [0, T] (see Event-aligned PSTHs per target interval). This procedure was repeated 1,000 times for each neuron, epoch, and TI, yielding, at each 1-ms time bin, a null distribution of firing-rate values.

For each 1-ms time bin, we computed one-sided permutation p-values for positive and negative deviations, using an empirical count with a +1 correction to avoid zero p-values. Let *R_obs_*_(_*_t_*_)_ denote the observed smoothed firing rate at time bin t, and let *R*^(*b*)(*t*)^ denote the smoothed firing rate of the b-th permuted spike train, with b = 1, …, B (B = 1,000). The p-value for a positive deviation was computed as:

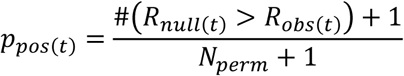

and the p-value for a negative deviation was computed as:

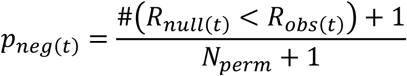

A time bin was considered significant when the corresponding p-value was below 0.01. Significant bins were labeled as positive activity when p_pos(t) < 0.01 (i.e., R_obs(t) exceeded the null distribution) and as negative activity when p_neg(t) < 0.01 (i.e., R_obs(t) fell below the null distribution). Activation periods were defined as runs of at least 15 consecutive significant bins (15 ms) in Core and Belt and of at least 50 consecutive significant bins (50 ms) in the other four areas.

### Cluster identification

Neurons were clustered according to the similarity of their PSTHs, separately for each area and pooling both monkeys. For each neuron, we used the PSTHs of the synchronization epoch (see Event-aligned PSTHs per target interval), averaged across trials and serial orders and concatenated in time across the four TIs, so that each neuron was represented by a single temporal vector capturing its response across durations. As in the PSTH analysis, PSTHs were aligned to the stimuli in Core and Belt and to the taps in preSMA, Parietal, Putamen, and M1. Each concatenated PSTH was z-scored across all its time bins.

For each area, the z-scored PSTHs were embedded in a two-dimensional space using UMAP^107,108^, with 15 neighbors, a minimum distance of 0.1, and the Euclidean distance. The number of clusters K was selected on the UMAP coordinates by evaluating K = 2–10 with four criteria: the Silhouette, Calinski–Harabasz, and Davies–Bouldin indices, computed on k-means solutions (10 replicates per K), and the Akaike information criterion (AIC) of Gaussian mixture models (full covariance, 5 replicates, regularization of 10⁻⁶). Each criterion voted for its optimal K (maximum Silhouette and Calinski–Harabasz indices, minimum Davies–Bouldin index and AIC), and the K with the most votes was selected; ties were resolved in favor of the smallest K. Neurons were then grouped on the UMAP coordinates using k-means (squared Euclidean distance, 20 replicates, maximum of 1,000 iterations, random seed 1).

After the automatic clustering step, cluster assignments were visually inspected by examining both the separation of neurons in the UMAP embedding and the PSTHs of the neurons within each cluster, to verify that neurons grouped in the same cluster had similar PSTHs. When isolated neurons or subgroups showed PSTHs inconsistent with their assigned cluster, they were manually reassigned to the cluster with the most similar PSTH; this curation was applied only when the mismatch was evident from the PSTHs. The final clusters were numbered in order of the peak latency of their mean PSTH and were used in all subsequent analyses.

### Comparison between behavioral epochs

For each cluster (see Cluster identification), we averaged the PSTHs of its neurons separately for the synchronization epoch, the perception epoch, and the passive condition, using only the 850-ms TI (see Event-aligned PSTHs per target interval). As in the PSTH analysis, all PSTHs were aligned to the stimuli in Core and Belt, whereas in preSMA, Parietal, Putamen, and M1 the synchronization PSTHs were aligned to the taps and the perception and passive PSTHs to the stimuli. We then computed pairwise difference traces by subtracting the firing rates point-by-point in time (Synchronization − Perception and Synchronization − Passive). For each difference trace Δr(t), we quantified the total modulation by integrating separately the positive and negative components relative to a zero baseline. Specifically, we computed the area above zero as

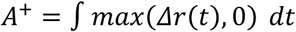

and the area below zero (in magnitude) as

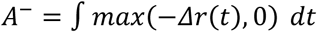

using trapezoidal numerical integration over the time vector t (in seconds, sampled at 10-ms resolution), so that both areas were expressed in spikes. For visualization, the negative area was plotted with a negative sign (i.e., −A⁻) to represent decreases in firing rate relative to the reference epoch or condition.

### Population response reliability

To quantitatively test whether the activation profiles of neurons depicted in Supplementary Figure 5 and Supplementary Figure 6 were robust and consistent within the 20 trial repetitions, we used a permutation test over repetitions using a geometric approach used before on the activation profiles of cell populations^48,71^. First, we constructed the heatmaps of Supplementary Figure 8 using the mean normalized activity for all the repetitions of the neurons constituting a cluster, sorting their time-varying activity by the peak activity latency. For self-sorted neural sequences in preSMA, we observed clear gradual patterns of activation (see Supplementary Figure 8A left). Then, we permuted a trial repetition for each cell and cross-sorted the neurons with the original peak order (Supplementary Figure 8A middle). Next, we computed the Euclidean distance matrix of the activity maps between the original mean repetitions and the cross-sorted permuted repetitions as depicted in Supplementary Figure 8A right. Finally, we identified the bin with the minimum distance, assuming that if the neural responses are consistent across trial repetitions the minimum distance will falls on the diagonal of the distance matrix across bins. The distance matrices shown in Supplementary Figure 8B and their associated minimum distances depicted as circles correspond to the mean of 1000 permutations.

### Temporal scaling index

To quantify whether population activity exhibited temporal scaling (i.e., a time-warped version across interval conditions) rather than a trivial time shift, we compared, for each cluster, the population-mean activity trace for the short interval (550-ms TI, *y_S_*_(_*_t_*_)_), against the population-mean trace for the long interval (850-ms TI, *y_L_*_(_*_t_*_)_), with 1-ms resolution. Each population-mean trace was the average of the firing rates of the neurons of the cluster in a single trial of the synchronization epoch and serial orders, aligned to the stimuli in Core and Belt and to the taps in preSMA, Parietal, Putamen, and M1. The analysis was performed separately for each of the trials (n=19), pairing the trials of the two TIs by their order. Before fitting, traces were mean-centered (*y* = *y* − ⟨*y*⟩).

We evaluated two competing linear models. A scaling (time-warp) model,

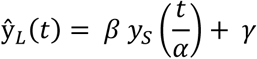

where α is the temporal scaling factor (short→long), and β and γ are a gain and an offset estimated by ordinary least squares after warping *y_S_* onto the long-time axis by shape-preserving piecewise cubic Hermite interpolation (pchip).

A shift (null) model,

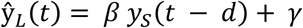

where Δ is a temporal shift, with β and γ again estimated by least squares after shifting *y_S_* and interpolating it onto the long-time axis. In both models, samples of the long-time axis that fell outside the short trace were extrapolated with the same pchip polynomials.

Both models were fit by grid search over their respective warp parameters: α was searched on 161 evenly spaced values spanning ±35% around 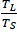, the ratio between the durations of the long and short traces, and Δ was searched from −300 to 300 ms in 2-ms steps. For each candidate parameter value, we computed the sum of squared errors (SSE) between *y_L_* and the fitted prediction ŷ*_L_*, and selected the parameter that maximized R² (equivalently, minimized SSE).

Finally, we computed the scaling index:

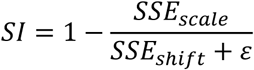

clipped to [0, 1]. Thus, [SI ≈ 1] indicates that scaling explains the long-interval activity substantially better than a simple shift, whereas [SI ≈ 0] indicates no advantage of scaling over shifting. For each cluster, the SI was summarized by its median across trials.

### Peak entropy & temporal sparsity

To quantify how strongly population activity formed a neural sequence (i.e., temporally ordered peaks with limited overlap), we computed the peak entropy (PE) and the temporal sparsity (TS) as defined by Zhou et al.^48^, using their MATLAB function SeqIndexDB^109^. For each cluster, we used the single-trial firing rates of the 850-ms TI described in Temporal scaling index, arranged as a three-dimensional array:

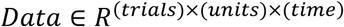

where each entry is the activity of neuron i at time bin t in a given trial. Both metrics were computed separately for each trial and summarized for each cluster by their median across trials.

For PE, we identified the peak time of each neuron, 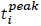, restricting the search to the bins of its activation periods (see Calculation of activation periods); neurons without activation periods were excluded from PE. We then constructed a histogram of the peak times with B bins centered at equally spaced times from 0 to T, where T is the number of time samples. The number of bins B was chosen adaptively as

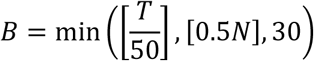

and the resulting value was then bounded below by 8 and above by N, the number of neurons in the cluster. Let *p_k_* be the fraction of the included neurons whose peak occurred in bin k. Peak entropy was

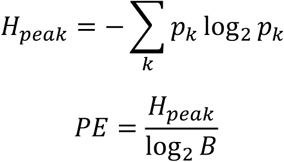

PE is maximal when peak times tile the interval approximately uniformly.

For TS, we normalized the activity of each neuron by the instantaneous population activity at each time bin t:

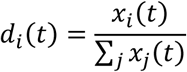

We then computed the entropy across neurons at each time,

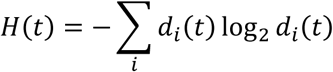

and averaged it across time:

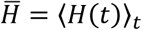

Temporal sparsity was defined as

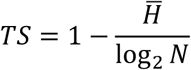

where *N* is the number of units. TS approaches 1 when, at each moment, activity is concentrated in few neurons (minimal overlap) and decreases when many neurons are co-active.

### Intrinsic timescale

To estimate the intrinsic timescale *τ*, we followed the approach of Murray et al. ^49^ and defined *τ* as the decay time constant of the spike-count autocorrelation during baseline activity. We used the control epoch, defined as the 1-s period preceding the first stimulus, pooling the trials of the four TIs. For each neuron, spikes were counted in 50-ms bins, yielding a matrix of trials × time bins. Trials without spikes in the control epoch and time bins with zero mean spike count across trials were excluded, and neurons with fewer than five trials or fewer than three bins with spikes were not analyzed.

For each lag k (corresponding to *Δ* = *k* × 50*ms*), we computed the Pearson correlation across trials between the spike counts of every pair of bins separated by that lag, (*i*, *i* + *k*). Letting *N*_1_ and *N*_2_ be the spike counts in bins *i* and *i* + *k*, the correlation was

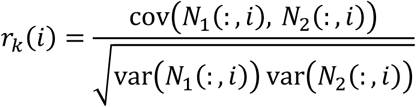

and only finite values were retained. For each area, correlations were pooled across bin pairs and across the neurons of all its clusters from both monkeys, and the autocorrelation function was summarized as the mean correlation per lag, *R_mean_*(*Δ*).

Finally, *τ* was obtained by fitting an exponential decay with an offset to *R_mean_*(*Δ*) for lags from 100 ms onward,

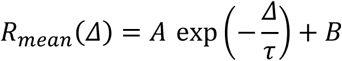

weighting each lag by its number of correlation values. The fitted *τ* was taken as the intrinsic timescale of each area, with larger values indicating a slower decay of the correlations in baseline activity.

### Peak-time distributions

This analysis was performed on the synchronization epoch for clusters 1 and 5 of Core, cluster 6 of Belt, and cluster 3 of preSMA, with activity aligned to the stimuli in Core and Belt and to the taps in preSMA (see Event-aligned PSTHs per target interval). For each target interval, trials were averaged separately for each neuron and serial order. Neurons with at least one significant bin within the corresponding target interval (see Calculation of activation periods) were retained. For each neuron and serial order, the peak time, *t_peak_*, was defined as the unrestricted global maximum of the mean firing-rate profile within the target interval. To prevent noise-driven maxima from contributing to the analysis, a peak was accepted only when its time bin was significant for that neuron and target interval; the maximum was not re-estimated within significant bins. Each neuron therefore contributed at most one peak per serial order and target interval.

Peak-time distributions were estimated separately for each target × serial-order combination by kernel density estimation (Gaussian kernel, 20-ms bandwidth) on a 1-ms temporal grid.

Densities were restricted to the corresponding target interval and normalized to unit probability mass, yielding one population distribution for each target × serial-order state.

Temporal separation between all pairs of target × serial-order distributions was quantified using the first-order Wasserstein distance,

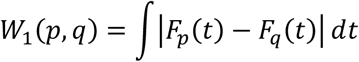

where *F_p_* and *F_q_* are the cumulative distribution functions of p and q, and the integral was evaluated numerically on the 1-ms grid. Because the underlying variable is time, Wasserstein distances are expressed in milliseconds. The resulting distances were grouped according to the absolute difference between target intervals, |ΔT| = 0, 100, 200, or 300 ms. The mean distance at |ΔT| = 0, [W0], quantified the dispersion among serial orders of the same target, whereas the mean distances at |ΔT| > 0 quantified the separation between different target intervals. This separation was summarized as the slope of the linear regression of the mean Wasserstein distance on |ΔT|, expressed in ms.

### Population trajectories in PCA space

For each area, we estimated a principal component analysis (PCA) space from the trial-averaged activity of the synchronization epoch, with activity aligned to the stimuli in Core and Belt and to the taps in preSMA, Parietal, Putamen, and M1 (see Event-aligned PSTHs per target interval). Neurons with at least 19 trials in each TI were included, and 19 trials were randomly selected for each neuron and TI. Neurons recorded in different sessions and in both monkeys were pooled into a pseudo-population for each area. For each neuron and TI, firing rates (1-ms resolution) were averaged across the 19 trials and the serial orders, so that each time point corresponded to a population activity vector whose elements were the firing rates of the individual neurons.

The PCA coefficients were estimated from the concatenation of the four TIs, yielding a single observations × neurons matrix. Before PCA, the activity of each neuron was soft-normalized by dividing it by its firing-rate range across all concatenated observations plus 10 spikes/s^110^, to reduce the influence of neurons with larger firing-rate ranges on the resulting population axes and then was mean-centered. Because longer TIs contain more time bins, concatenating them in absolute time would give them greater weight in the PCA. To avoid this, all TIs contributed the same number of time bins to the estimation of the coefficients: in Core and Belt, all TIs were truncated in absolute time to the shortest one (0–550 ms, 551 bins), and in the remaining areas, each TI was resampled by shape-preserving piecewise cubic Hermite interpolation (pchip) to 851 points on a normalized time axis. These procedures were used only to estimate the coefficients. The resulting coefficients defined orthonormal population axes; all available principal components were retained, and the variance explained by each component was quantified.

For all subsequent analyses, activity was projected in absolute time onto these fixed coefficients, after applying the same normalization factors and means used to estimate them. The projection was applied either to the trial-averaged activity or, for single-trial analyses, to each trial, with serial orders averaged within the trial; because neurons were pooled across sessions, single-trial trajectories combined the i-th selected trial of each neuron (pseudo-trials). For cluster-level analyses, the activity of the neurons of each cluster was projected onto the same axes using only their coefficients. Thus, the truncation or normalization of the time axis affected only the estimation of the PCA basis, whereas the analyzed trajectories preserved their absolute temporal structure, yielding time-resolved PC trajectories for each TI or trial within the same reference space.

To analyze these trajectories in three dimensions, we then selected, for each area, a subspace formed by three of these PCs, one of which was the component that best separated the TIs.

This separating axis was identified from the trial-averaged trajectories: for each PC, the offset of each TI was the mean of its projection over its whole trajectory in absolute time, and the separation score was the mean absolute difference between the offsets of all pairs of TIs. The PC with the highest score was selected as the separating axis; PCs whose score was within 1% of the maximum were considered tied, and ties were resolved in favor of the PC with the largest F statistic of a one-way ANOVA of the projected values grouped by TI and, if still tied, in favor of the PC with the largest explained variance. The other two axes of the subspace were the PCs with the largest explained variance among the remaining components, and the variance explained by the subspace was the sum of that of its three PCs.

### Participation ratio

Linear dimensionality was quantified from the PCA eigenspectrum of each recorded area using the participation ratio. After PCA was applied to the trial-averaged population activity, all principal components were retained and their explained variances were used to estimate the effective dimensionality of the population response.

The participation ratio was computed as:

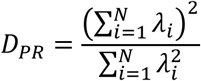

where *λ_i_* is the variance explained by the (i)-th principal component and (N) is the total number of principal components. The participation ratio estimates the effective dimensionality of the population response, with larger values indicating that variance is distributed across a broader set of PCA dimensions.

### Trajectory amplitude and speed

To quantify how trajectories evolved from the alignment event, we computed their amplitude and speed in the plane of the first two principal components (PC1–PC2). For a trajectory with coordinates [x(t), y(t)] in this plane, the reference point was its position at the alignment event:

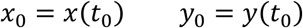

where [t₀] is the time of the tap in preSMA, Parietal, Putamen, and M1, and of the stimulus in Core and Belt. Amplitude was defined as the Euclidean distance from the reference point:

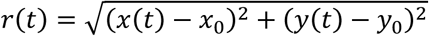

Instantaneous planar speed was computed from the time derivatives of the coordinates,

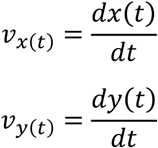

And the speed magnitude was defined as:

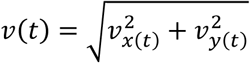

Derivatives were computed by central differences (one-sided at the trajectory edges) on a time vector in seconds with 1-ms resolution, spanning the duration of each TI, so that speed was expressed in arbitrary units per second; speed was then smoothed with a centered 51-ms moving average. Both measures were computed for the trial-averaged and the single-trial trajectories of each TI, for the whole population of each area and for each cluster.

### Ordinal trend of amplitude and speed

To quantify whether population trajectory features followed the ordinal structure of the TIs, we computed ordinal trend slopes separately for trajectory amplitude and speed in each area.

Amplitude and speed were obtained from the single-trial population trajectories (see Trajectory amplitude and speed). To compare the four TIs bin by bin, their traces were first placed on a common time axis. In Core and Belt, traces were analyzed in absolute time within the common window of the shortest TI (0–550 ms, 551 bins), so that each bin corresponded to the same time after the stimulus in all TIs. In preSMA, Parietal, Putamen, and M1, traces were resampled by shape-preserving piecewise cubic Hermite interpolation (pchip) to 851 points on a normalized time axis, so that each bin corresponded to the same fraction of the interval in all TIs.

The analysis was performed separately for each trial, pairing the i-th trial of the four TIs, which yielded 19 slopes per area and feature. At each time bin, the four TIs were ranked by the value of the feature at that bin: the four values were sorted in ascending order, and each TI received a rank from 1 (lowest value) to 4 (highest value). The rank of each TI was then averaged across all time bins, giving a mean rank that reflects the typical position of that TI relative to the other three throughout the trajectory. Finally, the mean ranks were fitted with a linear function of TI order (550, 650, 750, and 850 ms, coded 1–4), and the slope of this fit was taken as the ordinal trend slope. Positive slopes indicate that the feature increased with TI duration and negative slopes that it decreased; the slope reaches 1 or −1 when this ordering held at every time bin, and is close to 0 when the feature had no monotonic relationship with TI duration.

### Trajectory amplitude and produced interval

We evaluated the relationship between trajectory amplitude and the produced interval separately for each recording session and area. For each session, the activity of the simultaneously recorded neurons was projected onto the PCA basis of its area, computed from the neurons of all sessions (see Population trajectories in PCA space), using the coefficients of those neurons. Projections were obtained in absolute time, so that each TI kept its own duration, with activity aligned to the stimuli in Core and Belt and to the taps in preSMA, Parietal, Putamen, and M1. Sessions with fewer than 10 simultaneously recorded neurons. For each trial of the synchronization epoch, trajectory amplitude (see Trajectory amplitude and speed) was averaged across serial orders, and the produced interval was the mean of the intervals produced in that trial.

We compared two windows within each interval (1-ms bins). The tap window comprised the first and last 100 ms after the alienation event; the remaining bins formed the dwell window. Because the dwell window contained more bins than the tap window, it was reduced to the number of bins of the tap window of its area (200), so that differences between windows did not depend on the number of bins averaged. For each TI, a single contiguous block of that length was selected at a random position within the dwell window, whereas the tap window was always used in full.

For each trial, amplitude was averaged over the bins of each window. For each session and window, we pooled the trials of the four TIs and computed Pearson’s correlation coefficient (r) between mean amplitude and produced interval. The selection of the dwell block was repeated 50 times per session, and the median r across repetitions was retained as the dwell r of the session.

### Neural tap advance and asynchrony

We analyzed the relationship between neural tap advance and behavioral asynchrony separately for each area. We used the single-session projections described in Trajectory amplitude and produced interval (sessions with at least 10 simultaneously recorded neurons), without averaging across serial orders, for the five serial orders of the synchronization epoch (intervals 2–6), with activity aligned to the stimuli in Core and Belt and to the taps in preSMA, Parietal, Putamen, and M1. For each TI, a session was included when it contributed at least 15 trial × serial-order combinations. For each TI and serial order, we averaged the projected trajectories in the plane of the first two principal components (PC1 and PC2) across all trials of the included sessions, pooling both monkeys, together with the asynchrony of the tap at the end of each interval. This yielded 20 mean trajectories per area (four TIs × five serial orders), each with its mean asynchrony.

The tap position of each mean trajectory was its first time point, which corresponded to the tap in preSMA, Parietal, Putamen, and M1 and to the stimulus in Core and Belt. For each TI, the initial tangent of its mean trajectory across serial orders was defined as the unit vector from the first time point to the mean position over the following 10% of the trajectory, and the advance direction of the area was the normalized mean of the four tangents. Neural tap advance was then quantified as the projection of each tap position, relative to the centroid of the 20 tap positions, onto the advance direction.

For visualization, the trajectories of each TI were translated so that the centroid of its tap positions was at the origin, and advance and asynchrony were centered within each TI and plotted with the four TIs together.

### Spatial lofting and surface parametrization

To characterize the geometry defined by the condition-specific population trajectories, we constructed a continuous surface using a spatial lofting procedure. For each area, the mean trajectory of each TI (see Population trajectories in PCA space) was treated as a three-dimensional curve in the space defined by the first two principal components and the separating axis (PC7 in Core, PC8 in Belt, and PC3 in preSMA, Parietal, Putamen, and M1), and the surface was generated by interpolating between homologous points across trajectories rather than by fitting an explicit analytic surface model. Correspondence between trajectories depended on the temporal organization of each area. In preSMA, Parietal, Putamen, and M1, trajectories were linearly resampled to 180 points over normalized time, such that homologous points represented the same relative position within the interval. In Core and Belt, where population dynamics were analyzed in absolute time, trajectories were truncated to the duration of the shortest TI (550 ms), and homologous points corresponded directly to the same absolute-time bin across conditions.

The condition-specific trajectories were ordered according to the position of their geometric centers (the mean of the points used to build the surface) along the separating axis; no monotonic ordering with respect to TI was imposed. These centers were used to define a longitudinal coordinate, s, based on the cumulative Euclidean distance between consecutive trajectory centers. For each temporal coordinate, the corresponding points across conditions were then interpolated along s at 120 evenly spaced values using shape-preserving piecewise cubic interpolation (PCHIP), producing a continuous two-dimensional surface embedded in the three-dimensional PCA space. Thus, the resulting surface was parametrized by a longitudinal coordinate describing progression across condition-specific trajectories and a second coordinate representing either normalized time or absolute time within each trajectory.

Trajectory speed (see Trajectory amplitude and speed) was mapped onto this surface using the same correspondence and interpolation procedure used to construct its geometry. The instantaneous speed magnitude was first assigned to each point of the original trajectories and resampled or truncated along the temporal coordinate in the same way as the trajectories. Speed values at homologous points were then interpolated across the longitudinal surface coordinate, yielding a continuous scalar field. Consequently, the speed map did not constitute an independent fit, but rather a pointwise dynamical property of the neural trajectories transported onto the reconstructed surface according to the same spatial lofting coordinates.

### Assessment of manifold compatibility

To evaluate whether the reconstructed neural surface was geometrically compatible with an embedded two-dimensional manifold, we assessed three complementary properties of its representation in three-dimensional principal component space: local smoothness, regularity of the parametrization, and absence of self-intersections. For each area, the analysis was performed on the parametric surface constructed as described in Spatial lofting and surface parametrization,

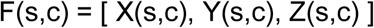

where [s] is the longitudinal coordinate across the interpolated trajectories and [c] the temporal coordinate along each trajectory. The surface was evaluated on its interpolation grid, with 120 rows along [s] and 180 columns along [c] in preSMA, Parietal, Putamen, and M1, or one column per 1-ms bin of the 550-ms TI in Core and Belt, and all distances in parameter space are expressed in grid steps. Because the local-planarity residual is curvature-driven, the row nearest to each original trajectory and the rows immediately before and after it were excluded from the interior points, so that the discrete change in curvature at the nodes would not be scored as a departure from smoothness. The outermost rows and columns were also excluded, since symmetric neighborhoods and central finite differences are undefined there.

Local smoothness was quantified by the extent to which surface neighborhoods were approximated by a local tangent plane. For neighborhood radii [R ∈ {1, 2, 4, 6, 8, 10}] grid steps, each neighborhood comprised the [(2R + 1) × (2R + 1)] grid points centered on an interior point. Singular value decomposition was applied to the centered three-dimensional coordinates within each neighborhood, and deviation from planarity was expressed as the fraction of variance orthogonal to the best-fitting plane,

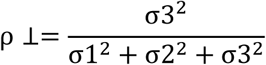

where [σ1 ≥ σ2 ≥ σ3] are the singular values. The median residual was computed over all interior points at [R = 1] and, at larger radii, over 400 regularly spaced interior points located at least 10 grid steps from the edges, so that all neighborhoods were complete. The dependence of the median residual on neighborhood size was quantified by the slope [β] of a linear fit of log₁₀ of the median residual on log₁₀ of [R]. The criterion was based on this scaling rather than on the residual itself, because at small radii [ρ⊥] is dominated by discretization and interpolation texture rather than by geometry. For a surface of finite curvature [κ], a neighborhood of extent [R] departs from its tangent plane by order [κR²], so that [ρ⊥ ∼ κ²R²] and [β ≈ 2] is expected. Slopes were fitted over the large-scale range [R ≥ 6] and over the full range of radii, and a surface was classified as locally smooth when [β_large ≥ 1.0] or [β_full ≥ 1.5], both conservative relative to this expectation.

Regularity of the parametrization requires the Jacobian of the map [(s, c) → ℝ³] to retain rank two throughout the surface. Tangent vectors [*T_s_*] and [*T_c_*] were estimated along the two parametric directions by finite differences with a step of one grid point, central in the interior and one-sided at the edges, and their linear independence was quantified by

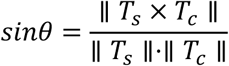

which approaches zero when the tangents become collinear. Because [sin θ] does not depend on the magnitude of the tangents, collapsed tangents were detected separately (norm below 10 times machine precision). Surfaces passed this criterion when [min(sin θ) > 0.05] (approximately 2.9°) over interior points and no tangent vector was degenerate. Degeneracies were screened over the complete grid, including interpolation nodes, since a collapsed tangent is a failure of parametrization wherever it occurs.

Embedding was evaluated by searching for pairs of surface points that were spatially close but distant in parameter space. Taking [*d_mes_*_ℎ_] as the median Euclidean distance between adjacent grid points along both parametric directions, candidate pairs closer than 3 [*d_mes_*_ℎ_] were identified over the complete grid and classified according to their parameter separation, with more than three grid steps treated as nonlocal. Longitudinal contacts ([Δs > 3]) indicate approach between distinct regions of the surface, whereas within-trajectory contacts ([Δs ≤ 3, Δc > 3]) indicate self-crossing along the trajectory direction. Seam contacts, between points located in the first two and the last two columns, which represent opposite sides of the open seam of a nearly closed trajectory, are expected and were excluded from the criteria. A surface satisfied the embedding criterion when the minimum distance in both contributing classes exceeded 0.5 [*d_mes_*_ℎ_]; classes with no detected contact could not cause failure.

A reconstructed surface was considered compatible with an embedded two-dimensional manifold only when it simultaneously satisfied the smoothness, regularity, and embedding criteria, the three outcomes being retained separately because they diagnose distinct geometric limitations.

### Sepparating-axis discriminalbility

To quantify whether the separating axis preserved condition-specific population states at the trial level, we analyzed the pseudo-trial projections (see Population trajectories in PCA space) in Parietal, Putamen, M1, and preSMA, in which the separating axis was PC3. Each pseudo-trial trajectory was summarized by its mean position along the separating axis across all time bins of the interval, in absolute time. This yielded one scalar position per pseudo-trial and, for each TI, a distribution of 19 positions.

Discriminability between adjacent TIs was quantified using the sensitivity index d′:

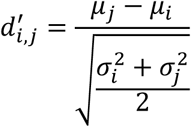

where µ*_i_* and 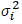 denote the mean and variance (with n − 1 in the denominator) of the pseudo-trial positions of TI i, and j is the next longer TI. d′ was computed for the 550–650, 650–750, and 750–850 ms pairs, and the minimum absolute value across the three adjacent comparisons was used as the summary measure for each area, providing a conservative estimate of separating-axis discriminability based on the least separated pair of neighboring TIs. The absolute value was used because the sign of d′ depends on the arbitrary sign of the principal component.

For visualization (Figure 15A,B), positions were z-scored within each area across all pseudo-trials, and the sign of the axis was set so that shorter intervals had higher values.

### Leave-one-condition-out classification

To determine whether the reconstructed population surface generalized to neural states not used during its construction, we performed a leave-one-condition-out (LOCO) analysis independently in Parietal, Putamen, M1, and preSMA. In each fold, one of the two intermediate TIs (650 or 750 ms) was held out, and the PCA basis was recomputed using only the remaining three TIs, with the same preprocessing as in Population trajectories in PCA space. Mean trajectories and individual pseudo-trial trajectories from all four TIs were then projected onto the first three principal components of this independently trained PCA space, and the population surface was reconstructed exclusively from the mean trajectories of the three included TIs, as described in Spatial lofting and surface parametrization.

Four candidate rings, corresponding to the four TIs, were defined on the reconstructed surface. The three rings associated with the included TIs corresponded to the longitudinal positions used to construct the surface. For the held-out TI, the longitudinal position was estimated by linear interpolation from its mean produced interval, using the relationship between mean produced interval and longitudinal coordinate defined by the three included TIs. The mean produced interval of each TI was computed from the first 19 trials of each session that contributed neurons to the area and had at least 19 trials in that TI, averaged across those sessions. The corresponding candidate ring was then obtained point-by-point from the continuous surface using PCHIP interpolation. This procedure used only the behavioral estimate of the held-out TI and no neural data from that TI. Thus, the candidate ring of the held-out TI was a genuinely interpolated trajectory that had contributed neither to PCA estimation nor to surface construction.

Each pseudo-trial trajectory of the held-out TI was linearly resampled to the 180 points of the rings and compared with all four candidate rings. For candidate ring i, trajectory-to-ring dissimilarity was quantified as the mean pointwise Euclidean distance across the parametrized cycle:

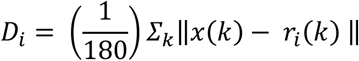

where *x*(*k*) and *r_i_*(*k*) are the k-th of the 180 points of the pseudo-trial trajectory and of candidate ring i in the space of the first three principal components, and ‖·‖ denotes the Euclidean norm.

Each pseudo-trial was assigned to the ring yielding the minimum mean distance. Classification accuracy was calculated as the proportion of held-out pseudo-trials assigned to the candidate ring of their true TI. With four candidate rings, chance performance was 25%.

### Prediction of held-out trajectory dynamics

To test whether the interpolated surface captured the internal dynamics of neural trajectories not used during its construction, we quantified how well the surface predicted the amplitude and speed profiles of held-out intermediate TIs in Parietal, Putamen, M1, and preSMA. The analysis was performed independently for the 650– and 750-ms TIs, each held out in turn with the procedure described in Leave-one-condition-out classification: the PCA basis was recomputed without the held-out TI, the mean trajectories of all four TIs were projected onto its first three principal components, and the surface was reconstructed from the three included TIs. The expected longitudinal position of the held-out TI was determined from its mean produced interval as in that analysis, and the surface prediction was the ring obtained at that position by PCHIP interpolation along the surface.

Prediction performance was evaluated against a condition-blind baseline. The baseline trajectory was obtained by averaging the parametrically corresponding points of the three included TI trajectories. Thus, whereas the surface prediction used the behaviorally estimated longitudinal position of the held-out TI, the baseline represented the generic mean trajectory of the area without information about the held-out TI’s position on the surface.

Both predictions were compared with the observed mean trajectory of the held-out TI, projected onto the same PCA basis. All trajectories were linearly resampled to 180 points in normalized time ([τ ∈ [0, 1]]), so that corresponding points could be compared across TIs of different duration, and dynamic profiles were computed in the PC1–PC2 plane. Amplitude was defined as the distance of each point from the first point of the same trajectory, which corresponded to the population state at the tap. Speed was quantified as the norm of the displacement between consecutive points, smoothed with a 25-point moving average (the first value was repeated so that the profile kept 180 points); it was therefore expressed per step of normalized time rather than per millisecond.

For each profile, prediction error was quantified as the mean absolute deviation from the observed held-out profile, normalized by the mean magnitude of that observed profile:

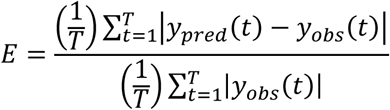

where *y_obs_*(*t*) denotes the observed profile (amplitude or speed) at point t, *y_pred_* (*t*) the corresponding prediction, and T = 180 the number of points of the normalized cycle. Equation E was evaluated twice for each profile: with the surface prediction as *y_pred_*(*t*), yielding *E_surface_*, and with the condition-blind mean trajectory as *y_pred_*(*t*), yielding *E_mean_*. Prediction gain was then defined as

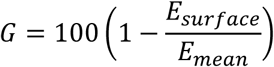

Positive values therefore indicate that knowledge of the held-out TI’s position on the reconstructed surface reduced prediction error relative to the condition-blind mean trajectory, whereas negative values indicate poorer prediction than the baseline. Errors from the two intermediate held-out TIs were averaged before computing the reported gain; that is, G was obtained as a ratio of average errors rather than as the average of the two per-TI gains.

### Error-related population trajectories

Error-related population trajectories were analyzed in Parietal, Putamen, M1, and preSMA, separately for early and late errors (see Behavioral measures). Early-error trajectories were constructed from the longest TI (850 ms), whereas late-error trajectories were constructed from the shortest TI (550 ms). For each error family, the analysis included only neurons with available data in both correct and error trials that were represented in the PCA space described in Population trajectories in PCA space. Thus, each error family and its corresponding correct reference were constructed from the same neuronal population, pooling both monkeys.

Because these neurons were not recorded simultaneously, the resulting trajectories represent trial-averaged pseudo-population activity rather than instantaneous population states.

Activity was aligned to the taps, with the same time window as for correct trials (see Event-aligned PSTHs per target interval). For each neuron, firing-rate profiles were averaged across error trials separately for each relative serial order with respect to the error (RSOE): the two cycles preceding the behavioral error (RSOE−2 and RSOE−1) and the error cycle itself. The corresponding correct reference was obtained by averaging correct trials of the same TI and then averaging across serial orders 3–5. These population firing-rate matrices were normalized and centered using the same preprocessing parameters as the original PCA decomposition and were subsequently projected onto the PCA coefficients of the neurons included in each family.

The resulting time-resolved projections defined, for each error family, one population trajectory for the correct reference and one trajectory for each of the three error-related cycles (RSOE−2, RSOE−1, and the error cycle).

### Geometry of error-related trajectories

Changes in population trajectory geometry were quantified separately for early and late error families across RSOE−2, RSOE−1, and the error cycle. Each error trajectory was compared with the correct trajectory constructed from the same neuronal population and TI (see Error-related population trajectories), thereby avoiding comparisons across populations with different neuronal compositions. Before all measurements, trajectories were smoothed with a 6-sample moving average.

Changes in trajectory shape within the principal population plane were quantified using Procrustes analysis in the PC1–PC2 plane. Before comparison, correct and error trajectories were linearly resampled to 850 equally spaced points in normalized cycle time to establish pointwise correspondence. Each error trajectory was then fitted to its correct reference, allowing translation, rotation, and uniform scaling, but not reflections. Two quantities were retained. The Procrustes scale was defined so that values above 1 indicate that the error-related trajectory was larger than its correct reference. The Procrustes residual dissimilarity was the residual sum of squares after the fit, divided by the sum of squared deviations of the correct trajectory from its centroid, and quantified the remaining difference in trajectory shape after accounting for position, orientation, and scale. Procrustes analysis was restricted to the PC1–PC2 plane so that changes in cycle geometry were evaluated independently of displacement along the separating axis.

Position along the separating axis was evaluated at the beginning of each cycle (its first sample), corresponding to the tap. Because the direction of PCA axes is arbitrary, the sign of the separating axis was standardized within each area such that shorter TIs occupied higher values. The sign was chosen from the linear trend of the mean separating-axis position across the four correct TIs (serial orders 3–5), computed from the neurons with data in all four TIs. To permit comparisons across areas, tap positions were then centered at the midpoint between the tap positions of the two correct references (the 850-ms reference of the early-error population and the 550-ms reference of the late-error population) and divided by their separation. Under this normalization, the two correct reference positions lie at ±0.5, and error-related values indicate their relative displacement along the separating axis.

Finally, separation between early– and late-error trajectories along the separating axis was quantified for each of the three cycles from the distributions of their time-resolved separating-axis values across all samples of the cycle. Both distributions were discretized using 40 common bins spanning their combined range, normalized as probability distributions, and their overlap coefficient was computed as the sum across bins of the minimum probability shared by the two distributions. Thus, values approaching 0 indicate little overlap, whereas values approaching 1 indicate highly overlapping distributions.

## QUANTIFICATION AND STATISTICAL ANALYSIS

### Behavioral statistics

Behavioral data were analyzed separately for each monkey, pooling correct trials across sessions; the trial was the unit of observation (Monkey 1, n = 8,610; Monkey 2, n = 9,009 correct trials). For each correct trial, asynchrony, produced interval, and constant error were averaged across serial positions, and the effect of TI on each measure was assessed with a one-way ANOVA, followed by pairwise post hoc comparisons with Bonferroni correction.

Asynchronies of correct and error trials were compared with a two-way mixed ANOVA, with trial type (correct or error) as a between-trial factor and RSOE (−3 to 0) as a within-trial factor, separately for early and late errors; p values were Greenhouse–Geisser corrected. Statistical significance was set at α = 0.05. F statistics, degrees of freedom, and p values are reported in the Results and figure legends; measures of center and dispersion are defined in the figure legends. Analyses were performed in MATLAB R2026a (version 26.1.0.3251617, Update 2; MathWorks) with the Statistics and Machine Learning Toolbox (anova1, multcompare, fitrm, and ranova).

Variance fractions and lag-1 autocorrelations from the variance decomposition were computed per session and are reported as means across the 220 unique sessions. Confidence intervals (95%) for λ and A* were obtained from 5,000 bootstrap resamples of sessions (with replacement; random seed 42), refitting the recursion to the pooled pairs of each resample and taking the 2.5th and 97.5th percentiles.

### Cluster-based permutation test

We assessed whether the differences between conditions (see Comparison between behavioral epochs) deviated from zero over time using a cluster-based sign-flip permutation test^111^, separately for each cluster and each comparison. For each cluster, we constructed a matrix D (N × T), where N is the number of neurons in the cluster and T is the number of 1-ms time bins, and each element represented the difference between conditions for one neuron at one time bin. At each bin, a one-sample t-statistic against zero was computed across neurons. Time bins exceeding an uncorrected two-sided threshold (p < 0.05, t distribution with N − 1 degrees of freedom) were grouped into temporal clusters of contiguous bins, and the mass of each temporal cluster was defined as the sum of the absolute t-values within it.

Significance was determined by sign-flip permutation^112^. On each permutation, the full temporal profile of each neuron was multiplied by +1 or −1 with equal probability, preserving within-neuron temporal structure while disrupting the consistency of the population-level effect under the null hypothesis. The t-statistic was then recomputed at each time bin, temporal clusters were re-identified using the same threshold, and the maximum temporal-cluster mass was retained. Repeating this procedure 5,000 times generated a null distribution of maximum temporal-cluster mass values. For each observed temporal cluster, the p-value was computed as the proportion of permutations in which the maximum null mass was greater than or equal to the observed mass, using an empirical count with a +1 correction to avoid zero p-values. Temporal clusters were considered significant when p < 0.05, thereby controlling the family-wise error rate across time.

Because temporal clusters were defined from the absolute t-statistic, this test established the presence of a significant temporal effect irrespective of sign. The direction of each temporal cluster was then assigned from the sign of the mean difference across its bins. For each cluster and each direction, we selected the longest temporal cluster lasting at least 50 bins (500 ms), and the effect was considered significant when its p-value was below 0.05. For this test, n is the number of neurons in each cluster, pooling both monkeys. Because the null distribution was obtained by permutation, the test did not require normally distributed differences. The areas and the significant effects of each cluster are shown in (Fig 2A, Supplementary figure 9A).

### Peak-time permutation tests

Statistical inference was performed independently for each cluster using 5,000 permutations (random seed fixed). Target-dependent changes in the full peak-time distribution were assessed with a Cramér–von Mises statistic comparing the empirical cumulative distribution of the accepted peaks of each target with the pooled distribution across the four target intervals,

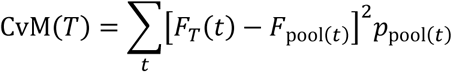

The null distribution was obtained by permuting target labels across the accepted neuron × serial-order peaks while preserving the original number of observations per target. For the Wasserstein distances, target labels were permuted across the target × serial-order population distributions while keeping the Wasserstein matrix fixed; for each permutation, the mean W was recomputed for each |ΔT|, together with the contrasts [W(ΔT) − W0] and the Wasserstein slope. Pairwise Wasserstein distances were therefore used descriptively and were not treated as independent replicates for inference. All permutation tests were one-sided in the direction of the predicted increase above the shuffled distribution, with empirical p given by

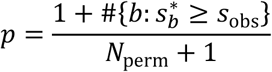

where *s_obs_* is the observed statistic, *s_b_* ∗ the statistic obtained in permutation b, and #{·} the number of permutations satisfying the condition. Benjamini–Hochberg FDR correction was applied separately to the peak-distribution tests (across clusters and targets), the W(ΔT) − W0 tests (across clusters and |ΔT|), and the slope tests (across clusters), and tests were considered significant at q < 0.05. For the Cramér–von Mises test, n is the number of accepted neuron × serial-order peaks of each target; for the Wasserstein tests, n is the number of target × serial-order distributions.

### Sign-flip tests of ordinal slopes

Amplitude and speed slopes were evaluated with sign-flip permutation tests^112^, using the mean of the 19 trial slopes as the test statistic and 10,000 permutations. The slopes of each feature were first tested against zero to determine whether they expressed a significant ordinal trend; amplitude and speed were then compared within each area by sign-flip permutation of their trial-wise differences. All tests were two-sided, with p-values computed with a +1 correction to avoid zero values, and were corrected with the Benjamini–Hochberg FDR procedure separately for the zero-slope tests (amplitude and speed across all areas) and for the paired amplitude–speed comparisons (across areas). Effects were considered significant at q ≤ 0.05. Here, n is the number of trials (19) per area.

### Permutation tests of amplitude and speed

We assessed whether trajectory amplitude and speed differed across TIs over time using cluster-based permutation tests^111^, separately for each area and each measure. The tests were applied to the amplitude and speed traces of the individual pseudo-trials (see Trajectory amplitude and speed), on the same time axes as in Ordinal trend of amplitude and speed: absolute time (0–550 ms) in Core and Belt, and normalized time (851 bins) in preSMA, Parietal, Putamen, and M1.

We first applied an omnibus test across the four TIs. At each time bin, Welch’s one-way ANOVA^113^ was computed across pseudo-trials, with TI as the factor. Time bins exceeding an uncorrected threshold (p < 0.05) were grouped into temporal clusters of contiguous bins, and the mass of each temporal cluster was defined as the sum of the F-values within it. On each of 5,000 permutations, the TI labels were randomly reassigned across the 76 pseudo-trials of the area, temporal clusters were re-identified with the same threshold, and the maximum temporal-cluster mass was retained. For each observed temporal cluster, the p-value was computed as the proportion of permutations in which the maximum null mass was greater than or equal to the observed mass, using an empirical count with a +1 correction to avoid zero p-values. Temporal clusters were considered significant when p < 0.05, thereby controlling the family-wise error rate across time.

When the omnibus test yielded at least one significant temporal cluster, the six pairs of TIs were compared with the same procedure, using Welch’s t-test^114^ at each time bin. Time bins exceeding an uncorrected two-sided threshold (p < 0.05) were grouped into positive and negative temporal clusters according to the sign of t, and the mass of each temporal cluster was defined as the sum of the absolute t-values within it. On each of 5,000 permutations, the TI labels were randomly reassigned across the 38 pseudo-trials of the two TIs, and the maximum masses of the positive and negative temporal clusters were retained separately. Thus, the p-value of each observed temporal cluster was computed, as in the omnibus test, from the null distribution of its own sign. Pairwise temporal clusters were considered significant when p < 0.0083 (0.05/6; Bonferroni correction for the six comparisons).

For these tests, n is the number of pseudo-trials (19 per TI). Welch statistics do not assume equal variances across TIs, and their parametric p-values were used only to form temporal clusters. Because significance was assessed by permutation, the tests did not require normally distributed data. The significant temporal clusters of the pairwise comparisons in preSMA are shown in Supplementary Figure 11.

### Tests of amplitude–interval correlations

The per-session r values (see Trajectory amplitude and produced interval) were tested separately for each area. Differences in coupling strength between the dwell and tap windows were assessed with a paired two-sided Wilcoxon signed-rank test on the absolute r values (|r|), and dwell correlations were compared with zero using a one-sample two-sided Wilcoxon signed-rank test on the signed r values. All valid sessions were included, without prior selection based on the significance of their correlations. For each test, p-values were corrected across the six areas with the Benjamini–Hochberg procedure^115^, and effects were considered significant at a false discovery rate of 0.05. For these tests, n is the number of sessions in each area, pooling both monkeys (sessions per area: Core, 15; Belt, 41; Parietal, 24; Putamen, 23; Motor Ctx, 14; preSMA, 28). Because the Wilcoxon tests are nonparametric, they did not require normally distributed r values. The distributions of r across sessions are shown in (Figure 7C).

### Tests of advance–asynchrony coupling

The within-TI relationship between advance and asynchrony was assessed separately for each area with a linear model that included TI as a categorical fixed effect (asynchrony ∼ advance + TI), so that the advance slope (β) was estimated from the covariation across serial orders within each TI. For comparisons across areas, advance and asynchrony were standardized within each area (z-scores across its 20 points) and the same model was refitted; the resulting standardized slope (βz) quantified coupling strength. Because the orientation of the advance direction is specific to each area, coupling was summarized as |βz|, whereas its SE and p-value were taken from the signed coefficient (two-sided t-test), which yields the same p-value as the unstandardized model. For these tests, n = 20 TI × serial-order combinations per area, pooling the trials of the included sessions of both monkeys (sessions per area: Core, 15; Belt, 41; Parietal, 24; Putamen, 23; Motor Ctx, 14; preSMA, 28). Significance was defined as p < 0.05, without correction across areas. |βz| for the six areas in [Figure 7F].

### Session-level jackknife

Uncertainty in the minimum absolute [d′] (see Separating-axis discriminability) was estimated with a session-level leave-one-out jackknife (Efron and Tibshirani, 1993). In each iteration, all neurons recorded in one session were removed, and the position of each pseudo-trial was recomputed as the sum of the contributions of the remaining neurons to its projection onto the original separating axis, without re-estimating the PCA. The minimum absolute adjacent [d′] was then recomputed from these positions. Because [d′] does not depend on the scale of the projection, the restricted axis was not renormalized. The jackknife standard error was calculated as

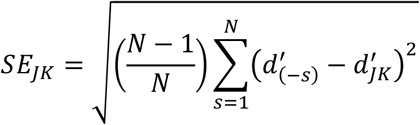

where N is the number of sessions that contributed neurons to the area, d′(−s) is the minimum absolute adjacent d′ obtained after excluding session s, and *d*^′^ is the mean of the leave-one-out estimates. The value calculated from the complete dataset is reported together with a 95% confidence interval defined as [d′ ± 1.96 *SE_jack_*]. For this analysis, n is the number of pseudo-trials per TI for d] (n = 19) and the number of sessions per area for the jackknife (Parietal, 24; Putamen, 23; Motor Ctx, 14; preSMA, 28). The minimum absolute d′ and its 95% confidence interval are shown for each area in Supplementary Figure 15C.

For the LOCO classification accuracy (see Leave-one-condition-out classification), the complete LOCO procedure, including PCA estimation, surface reconstruction, candidate-ring generation, and pseudo-trial classification, was repeated in each jackknife iteration after removing all neurons from one session. The mean produced intervals used to position the held-out ring were computed once from all sessions. Because accuracy is bounded between 0 and 1, jackknife accuracies were variance-stabilized using the angular transform

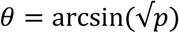

where p is the accuracy, and the jackknife standard error was computed in the transformed space with the same expression as for d′. A 95% confidence interval was constructed around the transformed accuracy of the complete dataset as [θ ± 1.96 SE_θ], constrained to the natural interval [0, π/2], and back-transformed to the accuracy scale as p = sin²θ. A confidence interval was obtained for each held-out TI. For this analysis, n is the number of pseudo-trials of the held-out TI (n = 19), and the number of sessions per area is the same as for d′. Accuracies, their 95% confidence intervals, and the chance level are shown in [Supplementary Figure 15 D].

For the prediction gains (see Prediction of held-out trajectory dynamics), the complete procedure, including PCA estimation, surface reconstruction, and the surface and baseline predictions, was repeated in each jackknife iteration after removing all neurons from one session, with the mean produced intervals computed once from all sessions. The jackknife standard error of G was computed with the same expression as for d′, directly on the gain scale, and the 95% confidence interval was defined as [G ± 1.96 SE_jack] around the gain of the complete dataset, without transformation. Gains were obtained separately for amplitude and speed, with one value per area, and the number of sessions per area is the same as for d′. Gains and their 95% confidence intervals are shown in Supplementary Figure 15 E,F.

