## Supplementary Information for "PreSMA population dynamics keep the beat atop the primate audiomotor hierarchy"

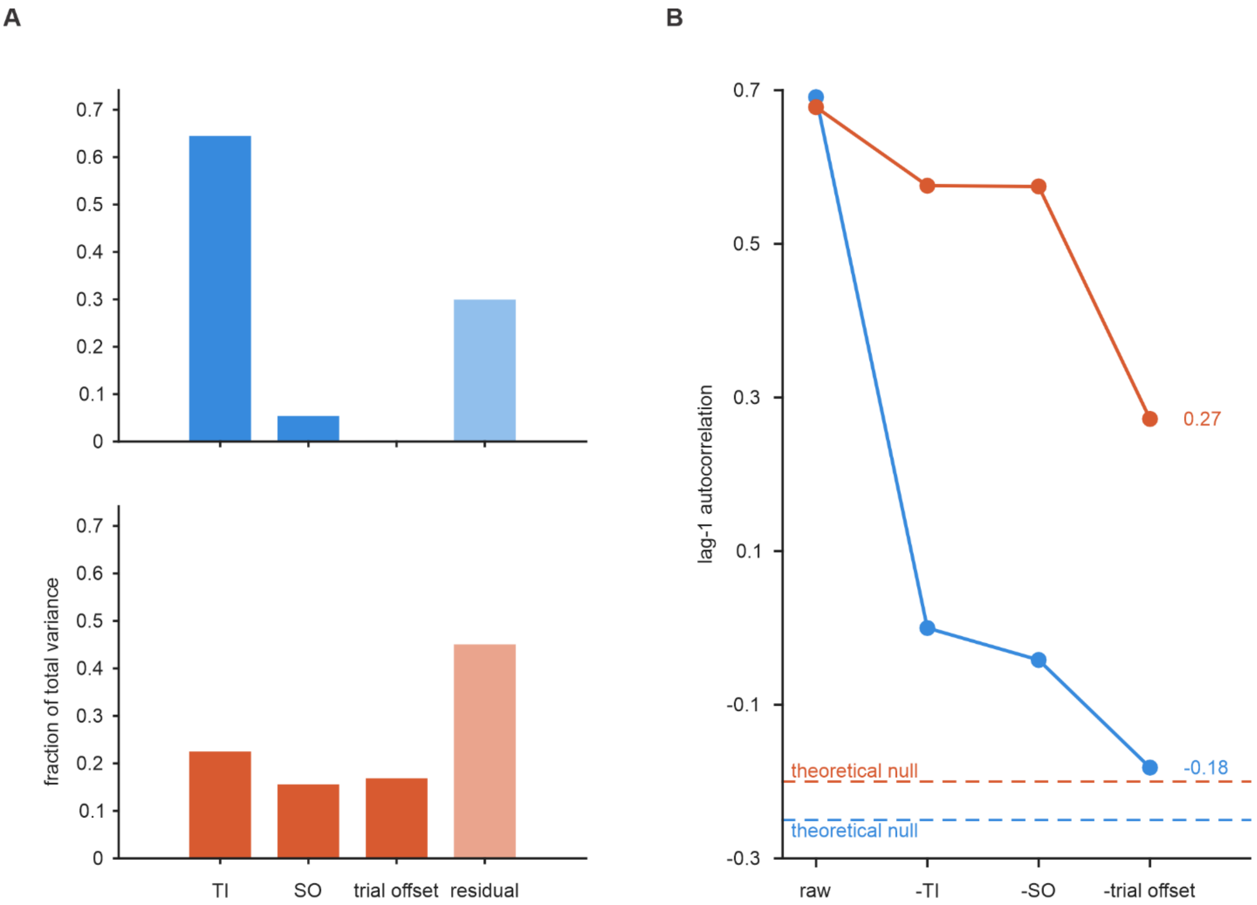


**Figure S1. Variance partition and residual variance structure of rhythmic timing during the PST, related to Figure 2. (A)** Variance partitioning of produced intervals and asynchronies across single recording sessions. Behavioral variance was decomposed into contributions from the target interval (TI), serial order (SO), trial-to-trial offset fluctuations, and residual variability. The produced interval variance was dominated by the TI, whereas the asynchrony showed larger serial-order, tempo, and residual components. **(B)** Progressive removal of deterministic behavioral structure from the lag-1 autocorrelation. Autocorrelation was calculated for the raw data and after sequentially removing target interval effects (-TI), serial-order profiles and trial-wise tempo. Produced intervals showed no remaining positive lag 1 dependence after correction, indicating that the sequential lag-1 structure of the produced intervals depends strongly on these parameters, especially the TI. In contrast, the asynchrony retained a large positive lag-1 autocorrelation, consistent with serial drift across tapping movements within all the sequential structure of the trial. Horizontal dotted lines indicate the theoretical null autocorrelation imposed by a within-trial mean centering of $-1/(nSO - 1)$. See also Figure 2G.

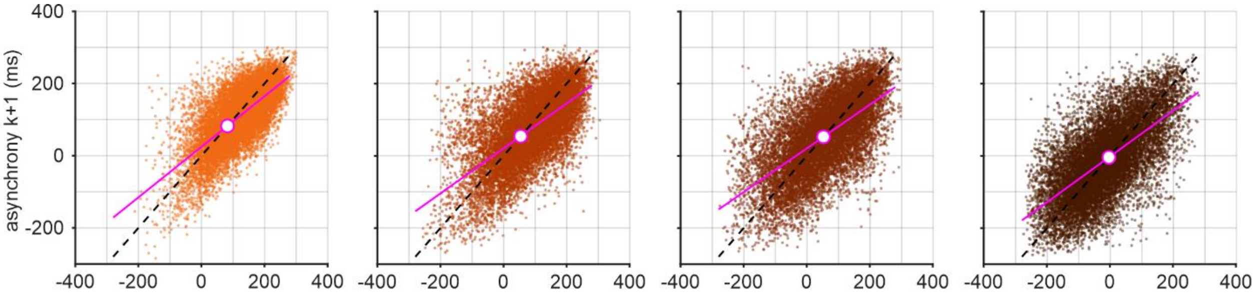


**Figure S2. Recursive correction of asynchrony, related to Figure 2.** Tap-to-tap asynchrony dynamics for each target interval, from correct trials of 161 sessions. Dashed lines are identity lines with a slope of one and indicate synchronization persistence without correction ($async_{k+1} = async_k$). Magenta lines show the fitted autoregressive relation, $async_{k+1} = c + \lambda \cdot async_k$. White circles depict the equilibrium point $A^* = \frac{c}{1-\lambda}$, where the fitted dynamics intersect the identity line.

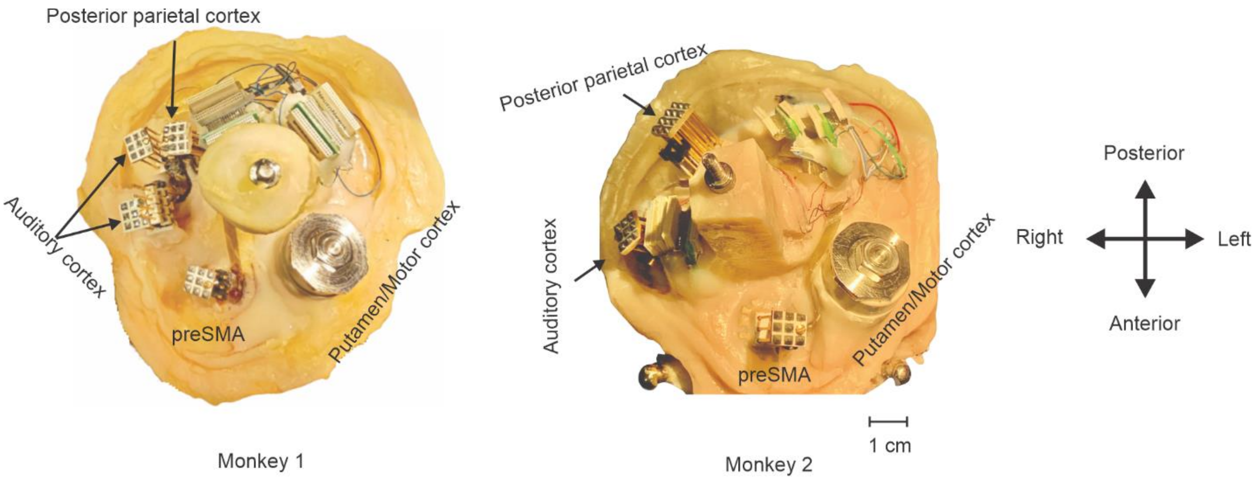


**Figure S3. Recording implants, related to Figure 1.** Photographs of the implants of monkey 1 (left) and monkey 2 (right). The small towers are screw micromanipulators used to drive high-density silicon probes along the z axis. Buzsaki-64 probes were used for preSMA and posterior parietal cortex (area 7a), and linear array electrodes for deep recordings in auditory cortex. A recording chamber was also placed for acute recordings in the putamen and motor cortex with linear array electrodes. The large post with a screw on top of the chamber supported a protective cap that maintained the sterility of the recording sites between experiments.

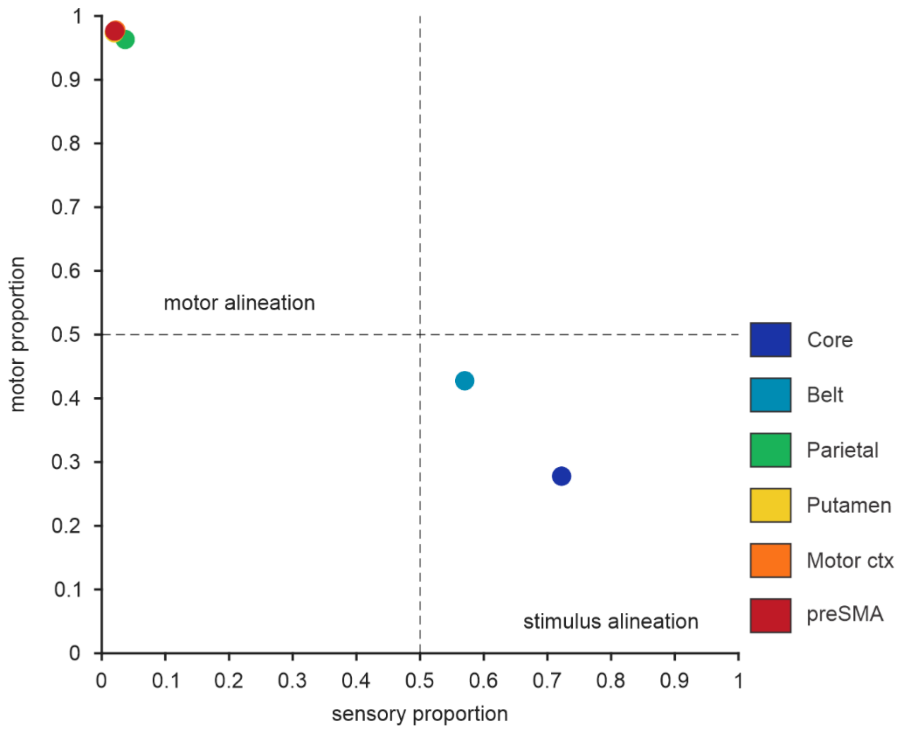


**Figure S4. Warping alignment of neuronal activity to stimuli or taps across areas, related to Figure 3.** Proportion of neurons classified as sensory (stimulus-aligned) versus motor (tap-aligned) in each area (one dot per area), relative to the total of both categories, determined with a warping analysis. The red dot (preSMA) overlaps those of the putamen and motor cortex.

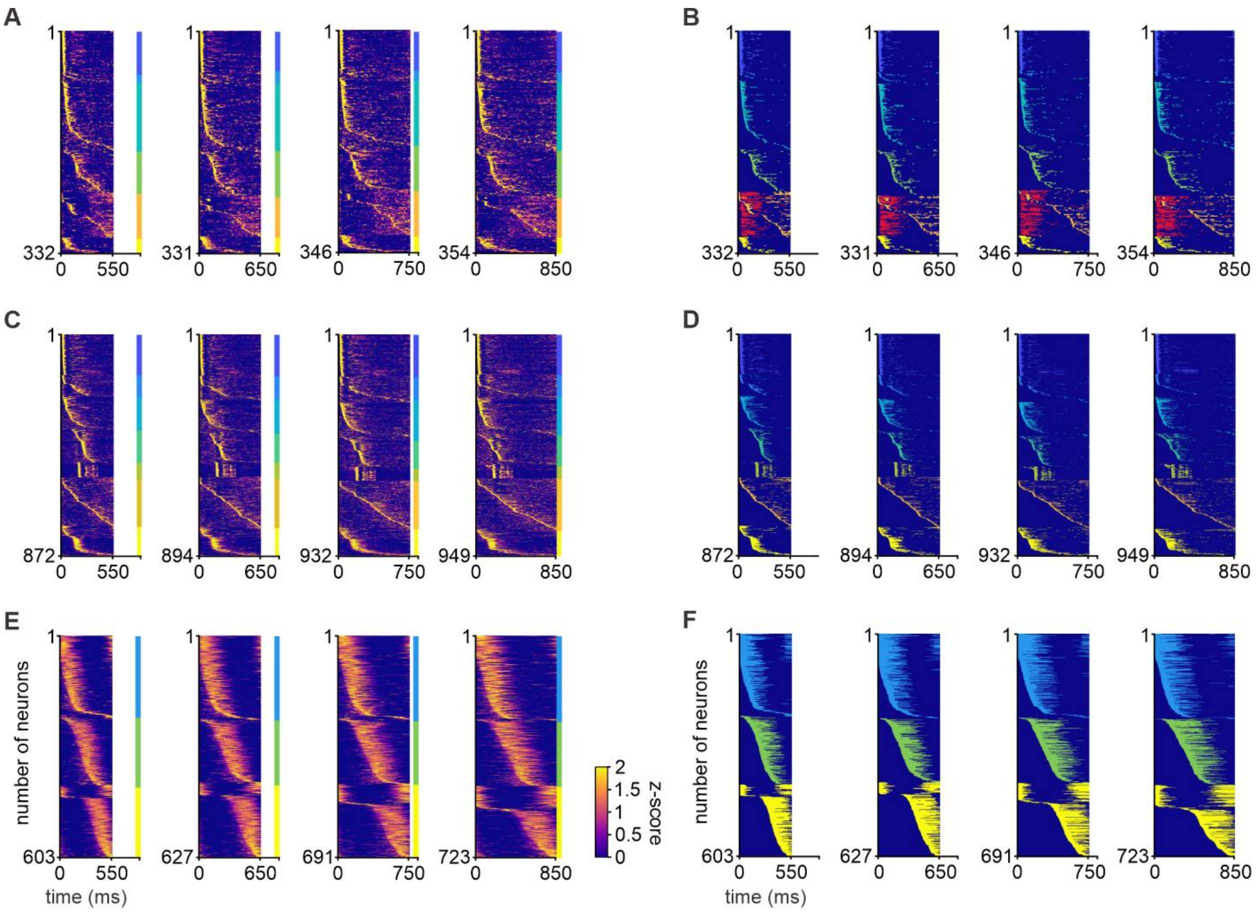


**Figure S5. Clustering and activation profiles of core, belt, and preSMA neurons during the PST, related to Figure 3. (A)** Heatmaps of the activation profiles of core neurons averaged across serial-order elements of the PST sequence, for each of the four TIs. The color bar (bottom right) shows the z-scored discharge rate. Neurons (y axis; total number at the bottom) are plotted as a function of time (x axis) and sorted into six clusters by UMAP followed by k-means clustering (see Methods); the colored vertical line identifies the clusters. Clusters 1–5 are aligned to stimulus onset and cluster 6 to the tap. Cluster 1 (blue) shows stereotyped, short-latency, short-duration responses phase-locked to the stimulus. Cluster 2 (cyan) shows an initial short-duration response followed by a response whose peak covers a substantial portion of the interval. Cluster 3 responds with longer latencies than cluster 1, and some of its neurons form a neural sequence that tiles the interval independently of the TI. Cluster 4 (green) shows short-duration responses with onset latencies longer than those of cluster 3. Cluster 5 (orange) shows a short-latency inhibition after stimulus onset (red inhibition periods) followed by a rebound of activity during the interval (orange activation periods) that forms a neural sequence covering the entire interval, including the 850-ms TI. Cluster 6 (yellow) is aligned to the tap, with responses after the tap and, in some neurons, long-latency responses closer to the next tap than to the previous one. **(B)** Activation periods of the neurons in (A). Each colored horizontal line marks the periods in which the discharge rate of a neuron was above baseline (permutation test; see Methods).

**(C and D)** Same as (A) and (B) for belt neurons, which formed seven clusters: clusters 1–6 aligned to the stimulus and cluster 7 aligned to the tap. Cluster 1 (blue) resembles core cluster 1, with short-latency and short-duration responses in phase with the metronome. Clusters 2–5 show phasic activity with onset latencies increasing from cluster 2 to cluster 5, without robust neural sequences. Cluster 6 (light orange) forms a neural sequence that covers the entire interval across TIs with a uniform recruitment rate. Cluster 7 (yellow) shows a burst just after the tap, a second response with a longer latency from the previous tap, and a third response within the produced interval until the next tap.

**(E and F)** Same as (A) and (B) for preSMA neurons, which formed three tap-aligned clusters that divided the interval between taps into roughly three equal parts: cluster 1 starting after the tap, cluster 2 with activity covering the entire interval independently of the TI, and cluster 3 with activity before the next tap. Numbers of neurons per cluster are given in Tables S2 and S3.

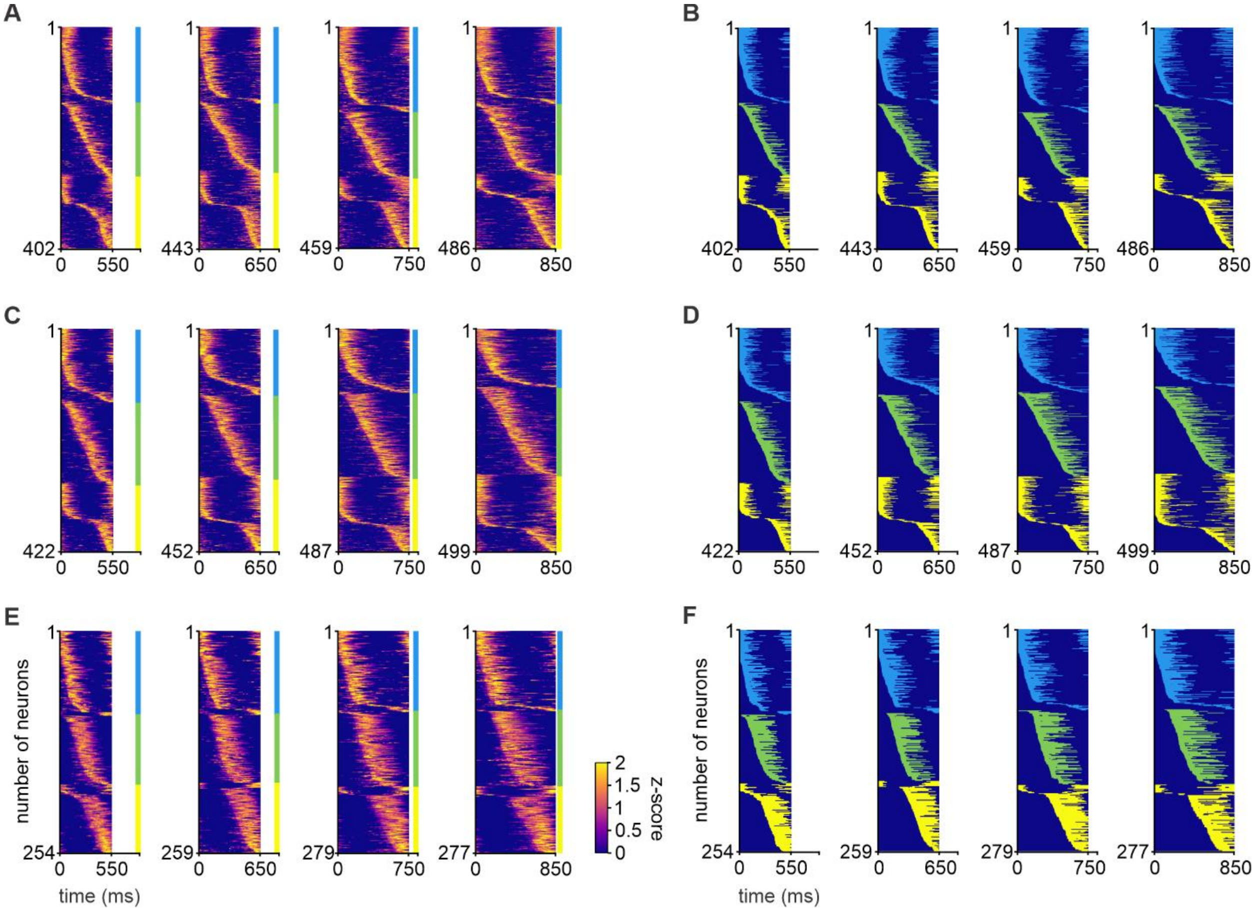


**Figure S6. Clustering and activation profiles of parietal, putamen, and motor cortex neurons during the PST, related to Figure 3.** Same organization as Figure S5. **(A)** Heatmaps of the activation profiles of parietal (area 7a) neurons averaged across serial-order elements of the PST sequence, for each of the four TIs. The color bar (right) shows the z-scored discharge rate. Neurons (y axis; total number at the bottom) are plotted as a function of time (x axis) and sorted into three tap-aligned clusters by UMAP followed by k-means clustering (see Methods); the colored vertical line identifies the clusters. **(B)** Activation periods of the neurons in (A), defined as in Figure S5B. **(C and D)** Same as (A) and (B) for putamen neurons, with three tap-aligned clusters. **(E and F)** Same as (A) and (B) for motor cortex neurons, with three tap-aligned clusters. Numbers of neurons per cluster are given in Tables S2 and S3.

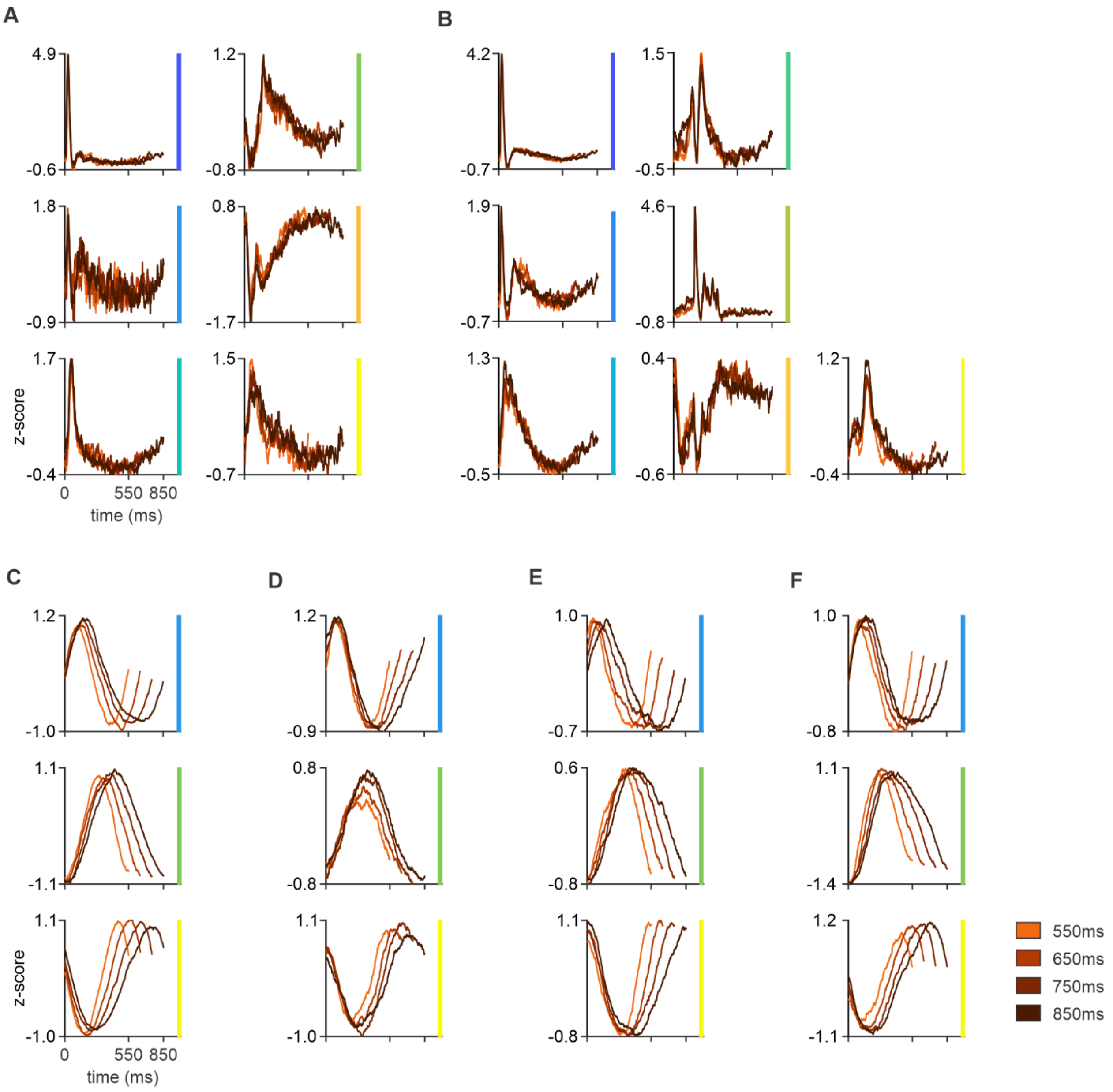


**Figure S7. Mean responses of neural clusters during synchronization, related to Figure 3.** Mean z-scored spike-density functions for the clusters identified independently in each area: core (A), belt (B), preSMA (C), parietal area 7a (D), putamen (E), and motor cortex (F). Core and belt responses are aligned to the stimulus or the tap according to the best alignment of each cluster, whereas preSMA, area 7a, putamen, and motor cortex responses are aligned to the taps. Within each cluster, traces correspond to the four TIs (color code on the right). Numbers of neurons per cluster are given in Table S2.

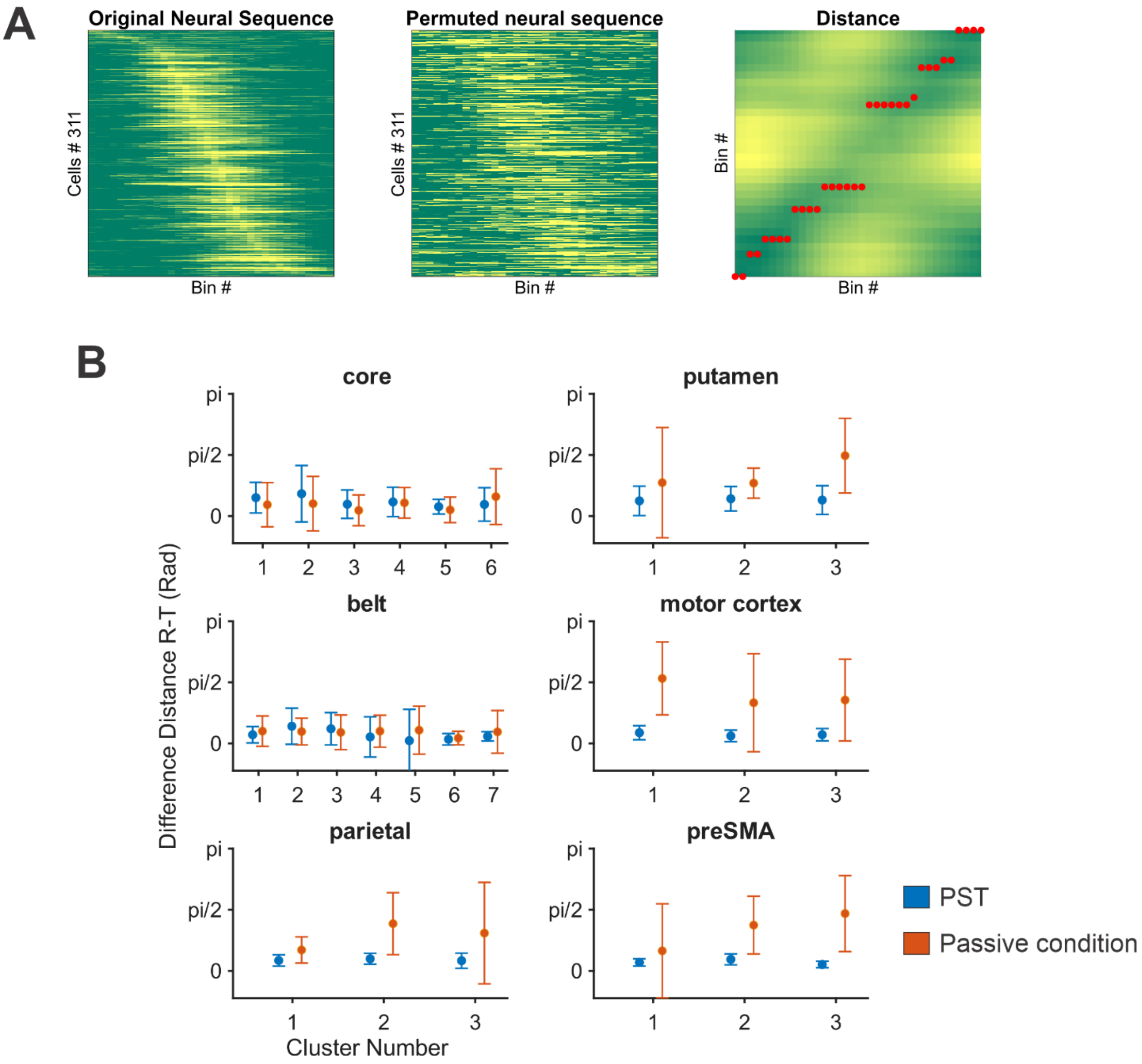


**Figure S8. Permutation test to measure consistency of neural sequences across trial repetitions, related to Figure 3. (A)** Left, neural sequence in preSMA: mean normalized activity of 311 neurons (y axis) across 20 repetitions, sorted by peak latency and plotted as a function of time (34 bins of 25 ms) for the 850-ms TI; color scale from green (low) to yellow (high activity). Middle, activity of the same neurons for one permuted trial, cross-sorted with the order of the left panel. Right, Euclidean distance matrix between the activity profiles of the left and middle panels; red circles mark the minimum distance in each bin. Most minima lie on or near the diagonal, indicating consistent response profiles across repetitions. **(B)** Circular mean ± SD of the difference (in radians) between the position of the minimum distance averaged across the 34 bins of 1,000 permuted trials (R, real) and the diagonal in (A, right) (T, theoretical), for the PST (blue) and the passive condition (orange). Differences between conditions were significant for all clusters of parietal cortex, putamen, motor cortex, and preSMA ($p < 0.001$), but not for all clusters of core and belt.

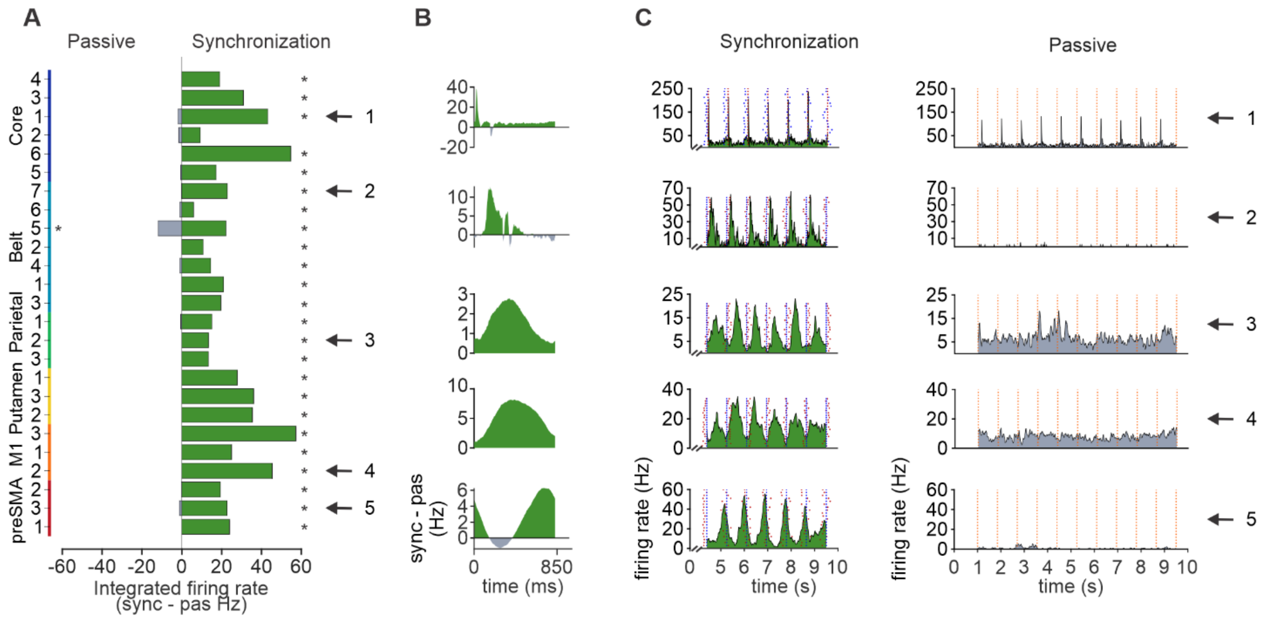


**Figure S9. Activity during synchronization versus passive listening across clusters and areas, related to Figure 3.** Same conventions as in Figures 3A–3C. **(A)** Difference in integrated firing rate between synchronization and the passive condition. The green bars of all clusters and areas lie on the synchronization side, indicating larger responses during the active task. **(B)** Synchronization-minus-passive difference in mean discharge rate as a function of time for the 850-ms TI, for the clusters marked by arrows in (A). **(C)** Discharge rate of prototypical neurons from the clusters marked in (A) during synchronization (left) and the passive condition (right) for the 850-ms TI. Significance was assessed with a temporal cluster-based sign-flip permutation test, as in Figure 3A.

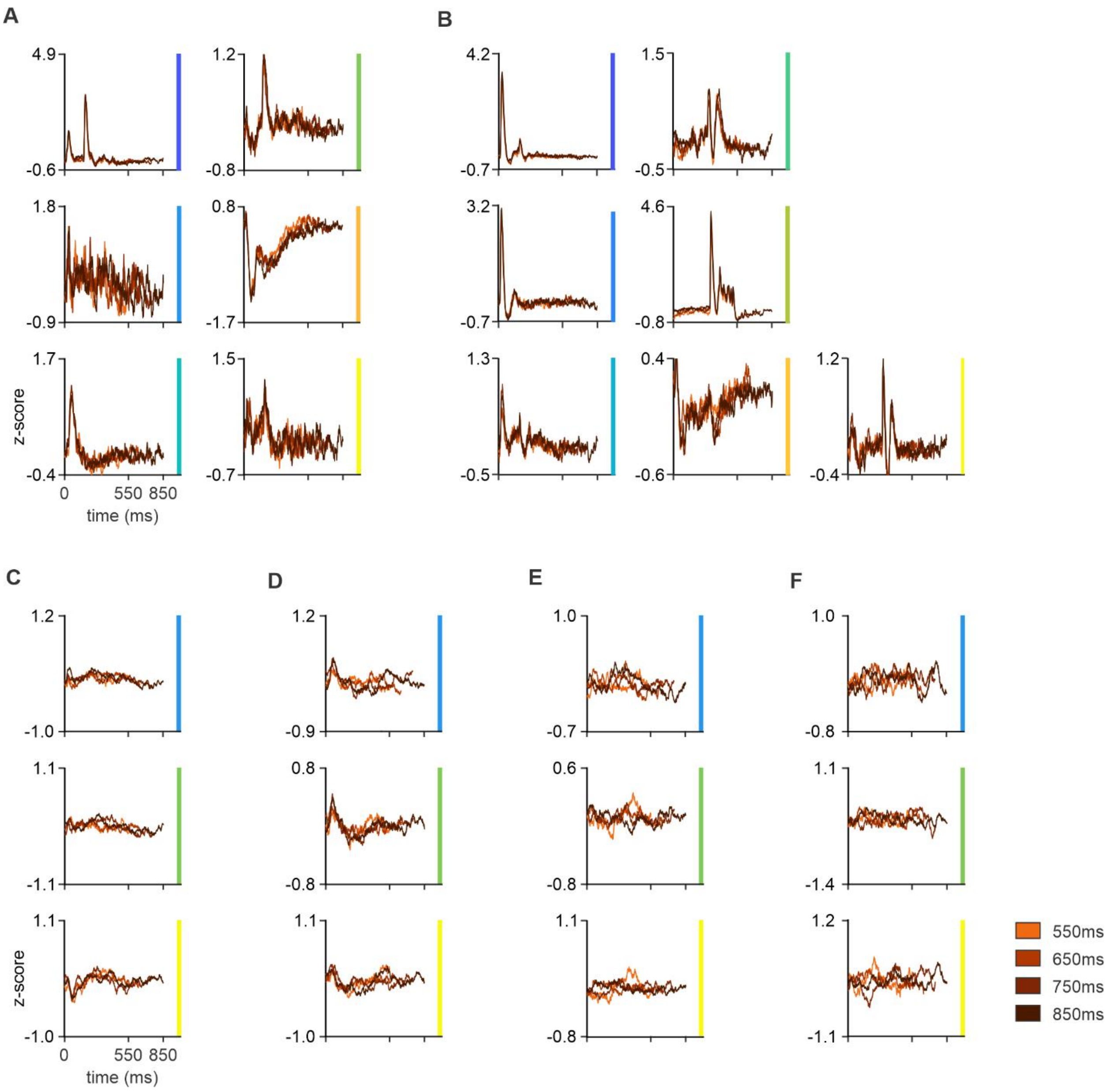


**Figure S10. Mean responses of neural clusters during passive listening, related to Figure 3.** Mean z-scored spike-density functions for the clusters identified independently in each area. Same format as Figure S7.

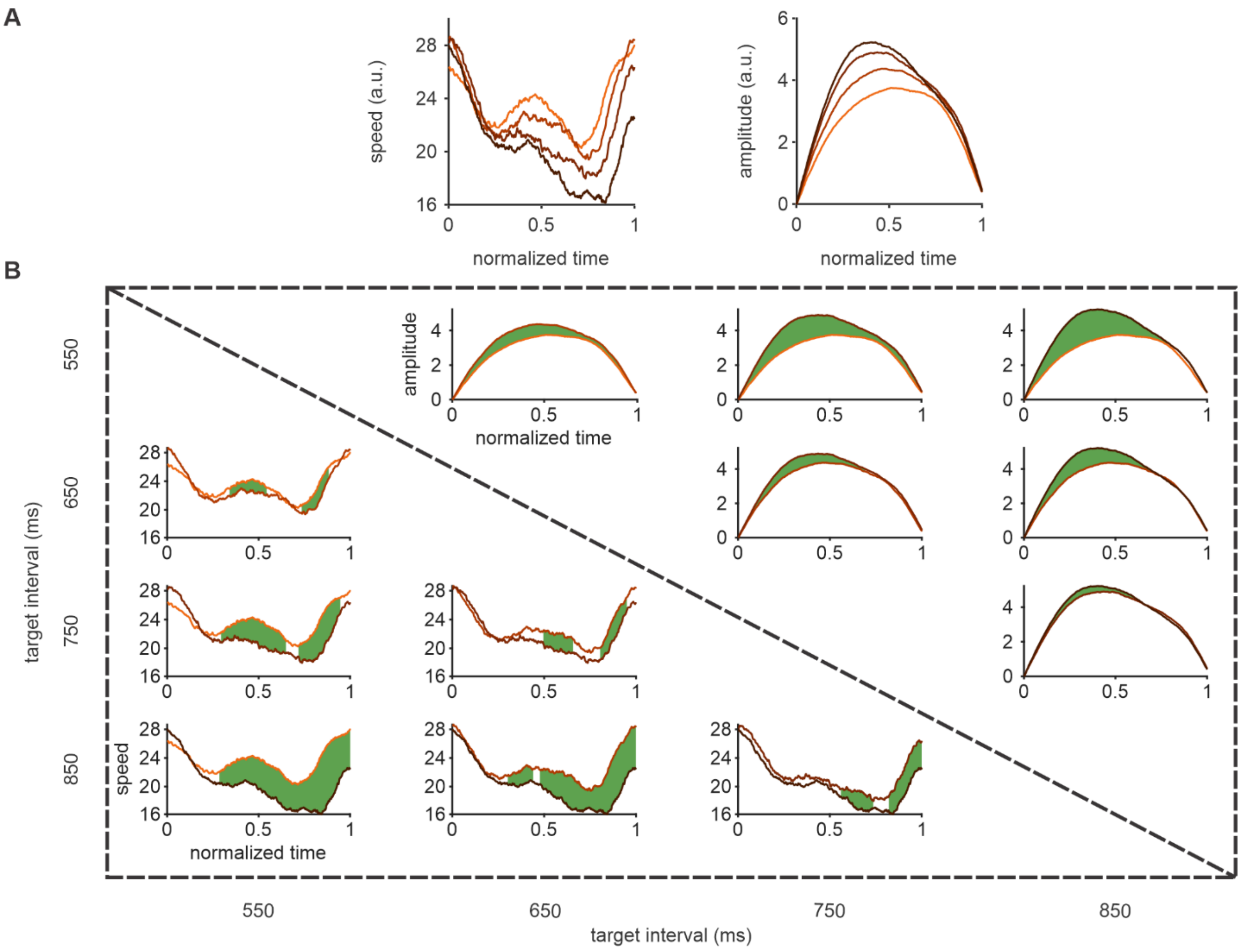


**Figure S11. Pairwise temporal differences in trajectory amplitude and speed across target intervals, related to Figure 5. (A)** Mean trajectory speed (left) and amplitude (right) of the complete preSMA population (980 neurons) for each TI. **(B)** Pairwise comparisons between TIs, with speed comparisons in the lower-left triangle and amplitude comparisons in the upper-right triangle. In each panel, the two curves correspond to the TIs indicated by the row and column. Green shaded regions mark temporal clusters with significant differences (pairwise cluster-based permutation tests with Welch's t test, 5,000 permutations, Bonferroni corrected across the six comparisons). Trajectories are displayed over normalized trial duration.

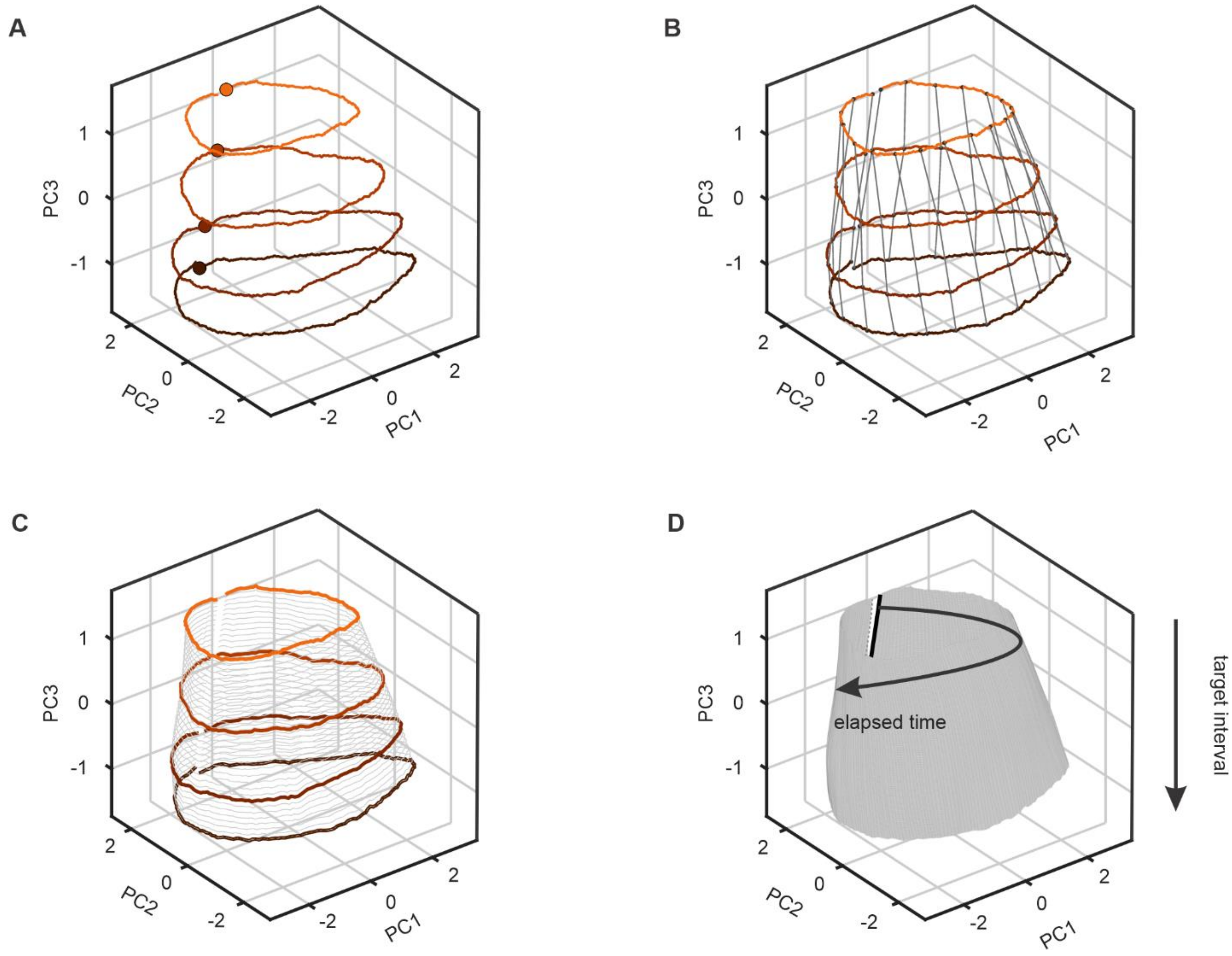


**Figure S12. Construction of the neural state manifold, related to Figures 5 and 6. (A)** Mean population trajectories for the four TIs projected into principal component space; filled circles indicate the tap. **(B)** Homologous points were established across conditions by resampling each trajectory to a common normalized time coordinate; gray lines connect corresponding points across trajectories. **(C)** The observed trajectories (colored) provide the constraints used to interpolate intermediate rings (gray), yielding a continuous surface across TIs. **(D)** Final open surface. The black seam marks the tap shared by all conditions. The surface represents the hypothesized continuum of population states between the sampled TIs; labeled arrows indicate the directions of elapsed time within an interval and of increasing TI. Pointwise trajectory variables, including amplitude and speed, are mapped onto the surface with the same parameterization.

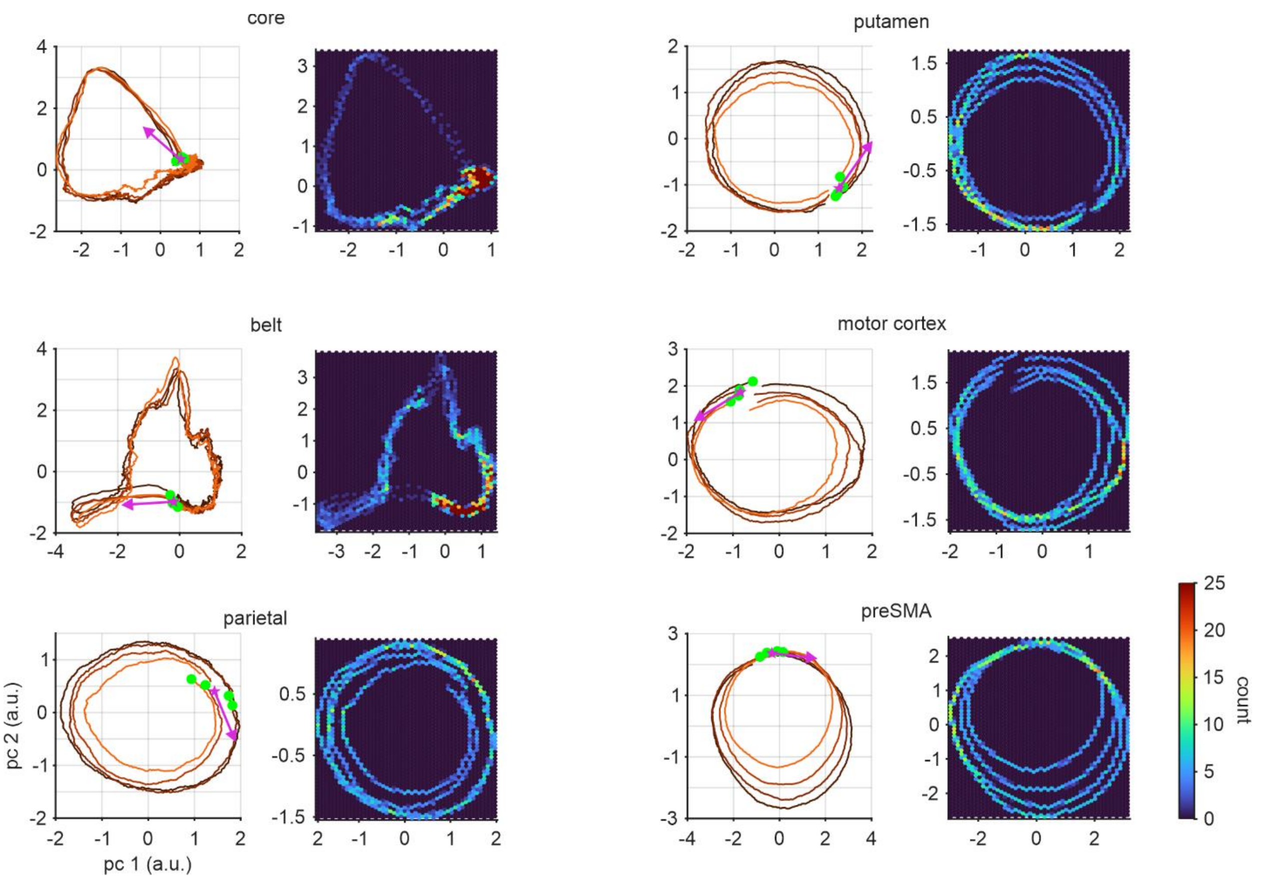


**Figure S13. Two-dimensional projections and density maps of population trajectories, related to Figure 5.** Left, population trajectories projected onto the first two principal components for each area. Each trajectory corresponds to one TI, and green dots indicate its starting point (stimulus for core and belt, tap for the other areas). Right, density maps obtained by pooling the trajectories of all TIs; color indicates the number of trajectory samples within each hexagonal bin. Higher-density regions correspond to portions of state space in which the trajectories spend more time, whereas lower-density regions indicate faster transitions.

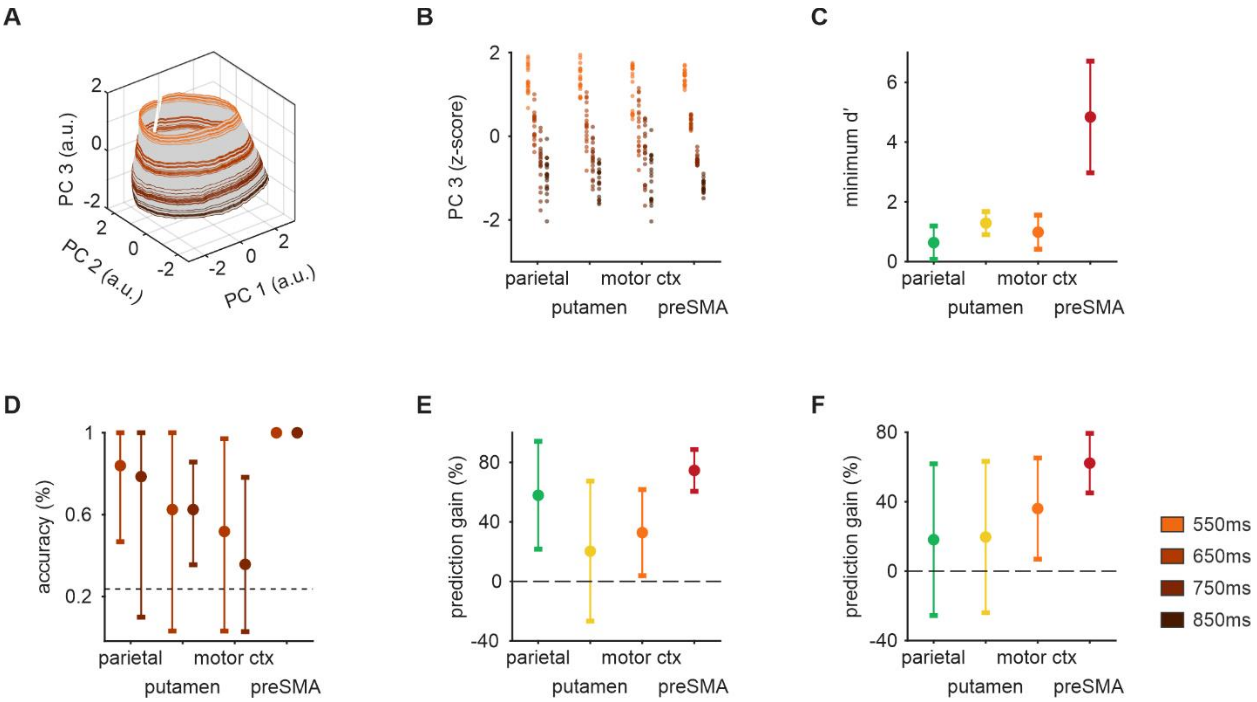


**Figure S14. Geometric validation of the population manifold, related to Figure 6. (A)** preSMA population surface reconstructed from the first three principal components (PCs); colored trajectories show trial rings for the four TIs (550–850 ms). **(B)** Distribution of trial positions along PC3 (separator axis) by area and TI. Each point represents the temporally averaged PC3 position of one trial, expressed as a Z score. **(C)** Minimum adjacent-condition d′ along PC3, quantifying worst-case trial-level discriminability between neighboring TIs. Points indicate values obtained using all sessions; error bars show 95% confidence intervals from a session-level jackknife. **(D)** Leave-one-condition-out (LOCO) classification accuracy for the intermediate TIs. For each held-out pseudo-trial, the complete trajectory projected into the LOCO PCA space was compared with the four candidate rings, and the trial was assigned to the ring with the smallest mean pointwise Euclidean distance. Accuracy denotes the proportion assigned to the ring of the held-out condition; the dashed line indicates chance performance (25%). Error bars show session-jackknife 95% confidence intervals computed after an arcsine-square-root transformation and back-transformed to the accuracy scale. **(E and F)** Prediction gain for the amplitude (E) and speed (F) profiles of held-out intermediate conditions. The interpolated surface ring and a condition-blind mean ring were independently compared with the observed mean population trajectory of the held-out condition. Prediction gain was defined as 100 × [1 − Esurface/Emean], where E is the relative mean absolute profile error; positive values therefore indicate lower prediction error for the surface than for the condition-blind mean trajectory. Profiles were computed in the PC1–PC2 plane. Points show values obtained using all sessions; error bars indicate 95% confidence intervals from a session-level jackknife. Dashed lines indicate zero prediction gain.

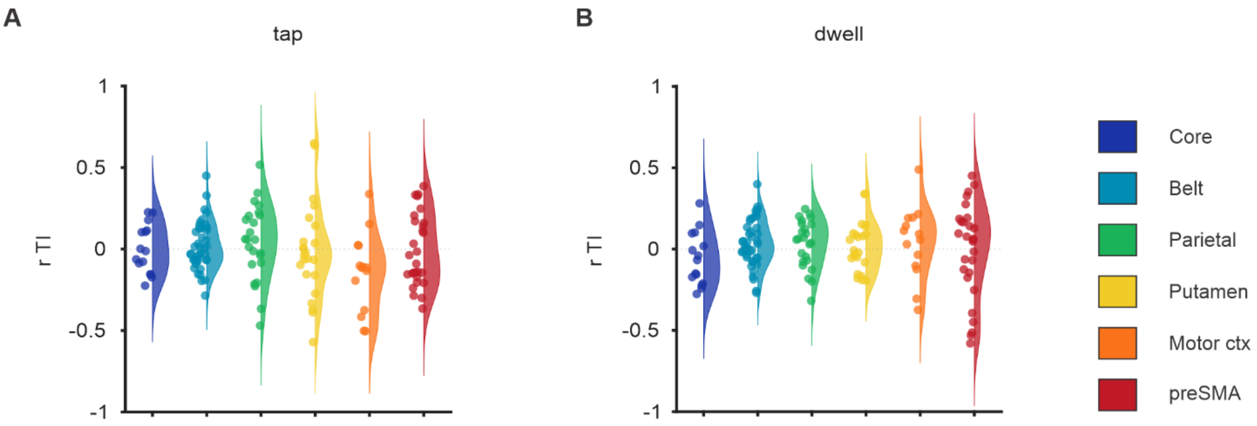


**Figure S15. Correlation between trajectory speed and produced interval, related to Figure 7.** Distribution across recording sessions and areas of the Pearson correlation between trajectory speed and produced interval during the movement (A) and dwell (B) epochs. One-sample Wilcoxon signed-rank tests showed no significant population-level correlations in either epoch.

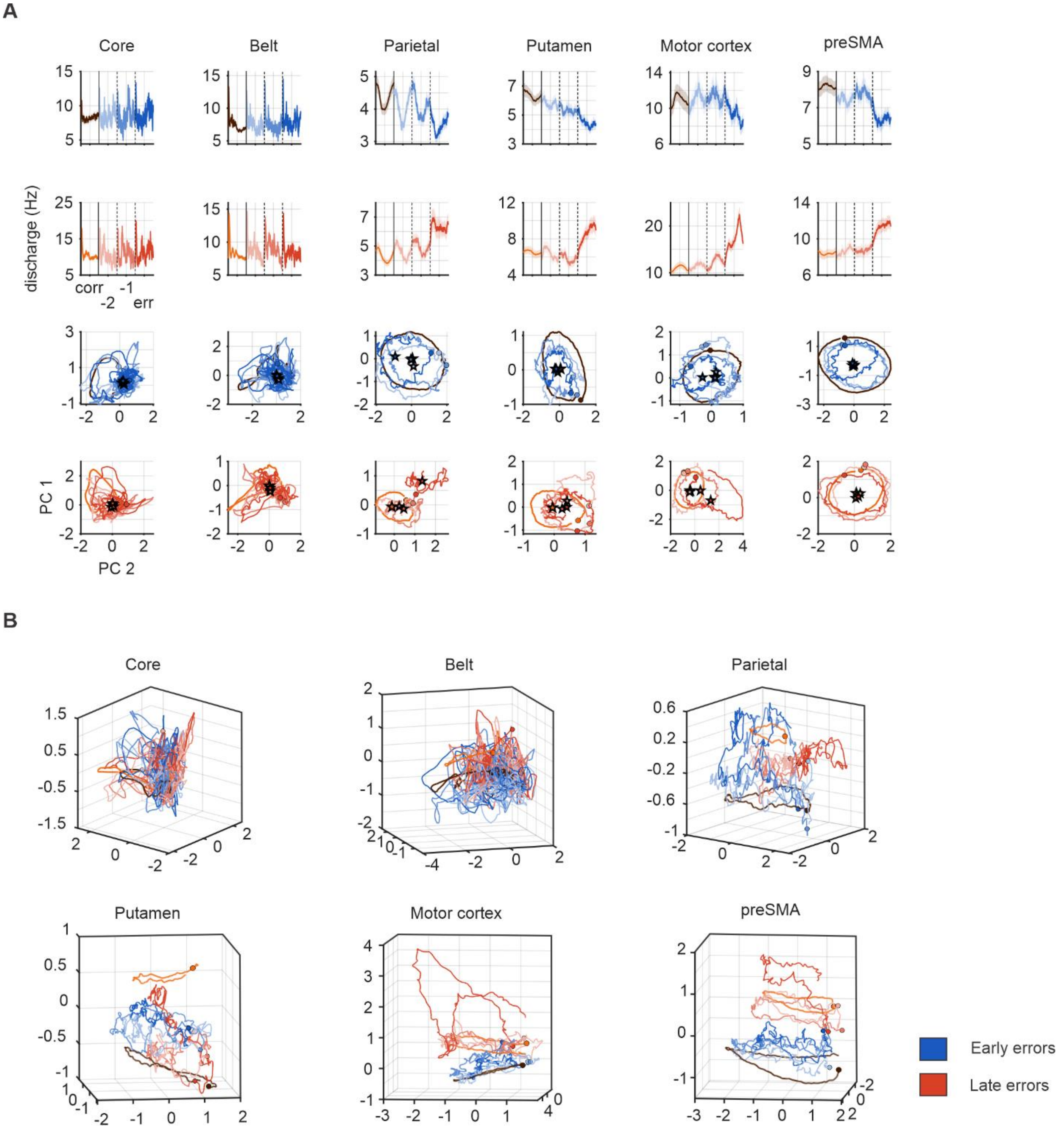


**Figure S16. Population trajectories are associated with timing errors across the audiomotor circuit, related to Figure 8. (A)** Population-average firing-rate profiles and corresponding trajectories in the PC1–PC2 plane for early (blue) and late (red) errors in core, belt, parietal cortex, putamen, motor cortex, and preSMA. Early-error activity is shown together with correct trials of the longest TI (850 ms; brown), and late-error activity with correct trials of the shortest TI (550 ms; orange). Shades indicate the serial order relative to the error (RSOE −2, RSOE −1, and the error cycle). Stars mark trajectory centroids. Dots indicate stimulus times in core and belt and tap times in the other areas. **(B)** Three-dimensional visualization of the corresponding population trajectories, showing early and late errors together with the correct long- and short-interval trajectories in population state space for each area.

**Table S1. Number of recorded neurons per area and monkey, related to Figure 1.**

| Total number of recorded neurons | | | | | | | |
|---|---|---|---|---|---|---|---|
| | Core | Belt | Parietal | Putamen | Motor Cortex | preSMA | Total |
| Monkey 1 (Tomas) | 255 | 644 | 778 | 503 | 68 | 670 | 2918 |
| Monkey 2 (Gil) | 206 | 589 | 224 | 215 | 257 | 310 | 1801 |
| Total | 461 | 1233 | 1002 | 718 | 325 | 980 | 4719 |

**Table S2. Number of neurons per area and cluster, related to Figure 3.**

| Total number of neurons by cluster | | | | | | | | |
|---|---|---|---|---|---|---|---|---|
| Area | Cluster number | | | | | | | Total |
| | 1 | 2 | 3 | 4 | 5 | 6 | 7 | |
| Core | 69 | 23 | 153 | 96 | 92 | 28 | | 461 |
| Belt | 197 | 136 | 218 | 178 | 65 | 316 | 123 | 1233 |
| Parietal | 312 | 336 | 354 | | | | | 1002 |
| Putamen | 214 | 269 | 235 | | | | | 718 |
| Motor Cortex | 124 | 110 | 91 | | | | | 325 |
| preSMA | 388 | 303 | 289 | | | | | 980 |

**Table S3. Number of neurons with significant activation periods per area, cluster, and target interval, related to Figure 3.**

| Neurons with activation periods | | | | | | |
|---|---|---|---|---|---|---|
| Area | Cluster number | Target interval | | | | Total Significative neurons |
| | | 550ms | 650ms | 750ms | 850ms | |
| Core | 1 | 63 | 62 | 60 | 65 | 65 |
| | 2 | 11 | 18 | 14 | 15 | 23 |
| | 3 | 104 | 100 | 105 | 108 | 143 |
| | 4 | 63 | 67 | 69 | 74 | 88 |
| | 5 | 68 | 59 | 72 | 67 | 87 |
| | 6 | 23 | 25 | 26 | 25 | 27 |
| Belt | 1 | 158 | 167 | 179 | 175 | 188 |
| | 2 | 83 | 96 | 106 | 95 | 134 |
| | 3 | 131 | 130 | 145 | 157 | 194 |
| | 4 | 133 | 124 | 126 | 135 | 167 |
| | 5 | 61 | 60 | 59 | 59 | 62 |
| | 6 | 190 | 202 | 199 | 214 | 296 |
| | 7 | 116 | 115 | 118 | 114 | 121 |
| parietal | 1 | 137 | 153 | 176 | 169 | 251 |
| | 2 | 132 | 136 | 135 | 153 | 244 |
| | 3 | 133 | 154 | 148 | 164 | 263 |
| Putamen | 1 | 139 | 135 | 140 | 131 | 187 |
| | 2 | 154 | 167 | 184 | 195 | 239 |

| | | | | | | |
|---|---|---|---|---|---|---|
| | 3 | 129 | 150 | 163 | 173 | 215 |
| Motor Cortex | 1 | 97 | 98 | 102 | 100 | 120 |
| | 2 | 79 | 79 | 94 | 94 | 106 |
| | 3 | 78 | 82 | 83 | 83 | 91 |
| preSMA | 1 | 223 | 239 | 262 | 277 | 350 |
| | 2 | 187 | 181 | 203 | 216 | 271 |
| | 3 | 193 | 207 | 226 | 230 | 267 |

**Table S4. Duration of activation periods per area, cluster, and target interval, related to Figure 3. Values are mean (SD) in milliseconds for the neurons in Figure 3 and Figure S7, for the six areas and their clusters at each of the four TIs. Asterisks in the last column indicate a significant effect of the TI (repeated-measures ANOVA, α = 0.01).**

| area name | cluster | 550ms | 650ms | 750ms | 850ms | Sign Diff |
|---|---|---|---|---|---|---|
| core | 1 | 50.3(26.5) | 49.2(24.7) | 48.8(21.5) | 50.6(23.5) | |
| | 2 | 33.8(17.6) | 38.3(26.5) | 46.8(32.4) | 42.2(33.2) | |
| | 3 | 41.0(26.2) | 48.4(31.3) | 51.1(31.0) | 57.9(36.2) | * |
| | 4 | 53.3(36.5) | 51.6(38.4) | 54.5(37.4) | 57.4(42.9) | |
| | 5 | 58.2(41.8) | 63.3(48.9) | 48.1(35.9) | 61.5(52.4) | |
| | 6 | 64.7(27.5) | 67.7(29.4) | 75.0(32.7) | 82.2(44.0) | |
| Belt | 1 | 52.9(28.4) | 50.3(28.9) | 53.5(37.5) | 58.0(44.4) | * |
| | 2 | 38.4(23.6) | 34.9(20.7) | 34.8(24.8) | 39.3(31.6) | |
| | 3 | 54.5(37.2) | 60.8(41.6) | 62.7(43.2) | 64.9(44.8) | * |
| | 4 | 45.1(25.5) | 44.5(25.7) | 48.3(29.6) | 57.1(41.0) | * |
| | 5 | 68.4(27.9) | 74.1(26.4) | 84.7(29.9) | 101.7(35.3) | * |
| | 6 | 51.9(42.7) | 46.7(40.0) | 50.2(45.6) | 54.3(50.0) | * |
| | 7 | 80.0(37.8) | 91.6(41.0) | 98.8(46.5) | 112.3(65.8) | * |
| Parietal | 1 | 134.2(65.4) | 164.4(86.1) | 185.6(104.3) | 198.0(117.9) | * |

| | | | | | | |
|---|---|---|---|---|---|---|
| | 2 | 113.1(59.7) | 135.5(68.7) | 148.5(80.5) | 153.8(96.1) | * |
| | 3 | 116.0(60.7) | 135.2(82.2) | 159.2(92.9) | 154.3(101.6) | * |
| **Putamen** | 1 | 147.5(64.5) | 168.3(73.7) | 176.0(80.4) | 198.8(95.7) | * |
| | 2 | 148.3(62.4) | 163.0(79.3) | 193.4(96.3) | 207.5(108.5) | * |
| | 3 | 132.7(55.2) | 152.3(77.3) | 180.0(85.2) | 203.7(103.9) | * |
| **Motor cortex** | 1 | 162.1(66.1) | 187.0(78.4) | 196.4(90.8) | 230.3(91.6) | * |
| | 2 | 182.7(71.4) | 196.5(85.7) | 224.7(101.1) | 261.8(120.0) | * |
| | 3 | 191.2(58.3) | 197.3(75.7) | 242.8(87.0) | 266.5(97.9) | * |
| **preSMA** | 1 | 164.1(77.8) | 192.6(92.2) | 205.3(104.9) | 223.7(106.9) | * |
| | 2 | 153.6(73.4) | 174.0(80.7) | 200.3(97.3) | 225.6(114.3) | * |
| | 3 | 172.1(65.3) | 191.9(82.1) | 212.4(91.8) | 236.9(111.6) | * |
